# Evidence for correlated evolution of sleep architecture in chronotype populations of *Drosophila melanogaster* created by artificial selection

**DOI:** 10.64898/2026.11.20.768060

**Authors:** Mansi Rathi, Neeti Badigannavar, Prerana Bhat, Vasu Sheeba

## Abstract

Chronotypes are circadian variants in timing of a behaviour. In humans, some studies have associated differences in sleep timing and quality with underlying circadian clock driven chronotypes although confounding genetic, environmental and social factors complicate the interpretation of these relationships. Here, we present evidence from artificially selected ‘early’ and ‘late’ eclosion chronotype populations of *Drosophila melanogaster* which suggest that divergence in chronotype can lead to correlated evolution of changes in sleep architecture. We find that *early* chronotype flies exhibit greater sleep fragmentation and shorter sleep latency during night, whereas *late* flies showed weaker and less consistent changes in the same direction. These differences were broadly seen for both sexes. We also show that environmental cues like temperature cycles can modulate sleep timing and enhance chronotype differences, while increasing light intensity delayed nighttime sleep timing similarly across chronotypes. Additionally, when sleep was disrupted by mechanical perturbations, *late* flies could not recover lost sleep to the same degree as *early* and *control* in a time-of-day dependent manner. In contrast, following simulated jetlag-induced circadian misalignment, *early* flies showed a greater increase in daytime sleep after phase advances. Our findings suggest that sleep and circadian traits can coevolve, offering insights into the complex interactions between chronotypes and sleep architecture. These populations provide an evolutionary model to investigate how circadian timing, sleep architecture and homeostatic regulation may be linked.

**Statement of significance:** Chronotype, the final manifestation of circadian clock regulation of a specific behaviour such as sleep onset, may influence not only timing of that behaviour but also how various aspects of underlying physiology of that organism is organised and regulated. We demonstrate that significant changes have evolved in sleep architecture, waveform, rebound and responses to circadian misalignment in divergent *early* and *late* chronotypes of *Drosophila melanogaster* populations that were created by artificial selection on the timing of a critical developmental stage under strong circadian control. The most consistent differences involve nighttime sleep organization, particularly increased sleep fragmentation in *early* chronotypes. Because these populations share ancestry and are studied under controlled conditions, they allow chronotype–sleep relationships to be examined without the confounds inherent to studies involving human subjects. Our findings suggest that selection on behavioural timing can have broader consequences for sleep, providing an evolutionary framework to investigate how circadian clocks and sleep characteristics are correlated.

## Introduction

A central role of the circadian clock is to time behaviour and physiological processes to occur at appropriate times of the day i.e., maintain a stable phase relationship with environmental cycles [1]. However, individuals within a population exhibit variation in this phase relationship, thus giving rise to chronotypes – a phenomenon documented across taxa and across variety of rhythmic behaviours [2–7]. Several studies suggest that the timing of a process can be as important as the behaviour itself, allowing forces of natural selection to act on chronotypes [3,7]. In humans, the timing of sleep (onset or midpoint) is primarily the basis of their classification into morning or evening chronotypes, and it is typically assessed using questionnaires like Morningness–Eveningness Questionnaire (MEQ) and the Munich Chronotype Questionnaire (MCTQ) [6,8]. Beyond differences in timing of sleep onset/offset, timing of physiological rhythms like melatonin, body temperature, heart rate and cortisol as well as clock gene expression rhythms in fibroblasts and peripheral clocks also differ between chronotypes, suggesting variations in the underlying circadian clock itself [9–13]. Chronotypes are also correlated with several health-parameters, and differences in sleep quality and architecture have emerged as potential contributors to these associations [14–22].

While many studies suggest that evening chronotypes are associated with poor sleep quality and sleep insufficiency, a few others suggest association of morning chronotypes with poor sleep quality, and sleep architecture differences such as more time in lighter sleep stage [16–20,23,24]. Interestingly, yet another study reports that sleep architecture is unchanged across chronotypes [21]. Along with sleep architecture, a few studies report variations in sleep homeostasis among chronotypes, with evening types exhibiting slower buildup of subjective sleepiness as well as sleep pressure decay rate, than the morning types [16,25]. On the other hand, chronotype differences in sleep were suggested to be reproducible with identical homeostatic parameters implying that chronotype differences may not accompany sleep homeostatic differences [26]. Such contrasting observations and inferences regarding associations between chronotypes and sleep architecture and homeostasis across studies may partly reflect variability in cohort composition, genetic and environmental heterogeneity, shift work, light exposure and lifestyle differences as well as method of assessment of chronotype and sleep quality [27–31]. One major confounding factor is social jetlag, i.e., chronic misalignment between internal circadian time and socially imposed schedules [27]. Social jetlag is reported to disproportionately affect evening chronotypes accounting for much of their reported sleep debt and increased sleepiness [23,24,27,32]. Difference in environmental exposure experienced by the extreme chronotypes provides another source of variation. Light being a dominant entraining cue for the circadian clock and can directly influence sleep [33–37]. GWAS studies have shown enrichment of variants in phototransduction pathways in chronotypes suggesting variations in photosensitivity [6,22,33,38,39]. Seasonal variation adds another layer of complexity as evidenced by evening chronotypes showing greater variation in sleep duration across the year [31,40]. The independent contribution of temperature cycles which co-vary with seasons remains largely unexplored. Broadly, the environmental, social and genetic factors that differ between human chronotypes are deeply intertwined and contribution of each to variation in sleep quality and architecture cannot be isolated in observational settings with limited experimentation. Hence, a model organism where these correlations can be studied systematically in controlled conditions is greatly advantageous in determining relationships between circadian clock variation and sleep properties and enables us to test how specific environmental inputs may modulate it.

The availability of experimentally evolved chronotype populations of *Drosophila melanogaster* in our laboratory provided a unique opportunity to study this relationship. These populations are a result of artificial selection on the timing of adult emergence from pupa (eclosion) which is a circadian clock-controlled population rhythm. Flies emerging in a four-hour window, around lights-ON (morning), denoted as *early*, and those emerging around lights-OFF (evening), denoted as *late* chronotypes have given rise to the divergent chronotype populations [41]. As a response to selection, these populations have diverged not only in the phase and period of the eclosion rhythm but also that of their locomotor activity rhythms and core clock gene expression rhythms as well as in their sensitivity to environmental time cues [41,42]. It has been demonstrated that these *early* flies have a faster running clock which assumes an advanced phase under cyclic conditions whereas *late* flies have a slower running clock with delayed phase as compared to their ancestor *control* populations [42]. With a replicate-population design of four independently evolving populations per chronotype, derived from a common ancestor and reared under similar environmental conditions we have control for genetic and environmental heterogeneity, and it strengthens our confidence on the correlations observed [43]. Additionally, unlike single-gene clock mutants, which produce chronotype-like shifts through disruption of specific molecular components, these experimentally evolved populations achieve chronotype divergence through polygenic changes in standing genetic variation, a mechanism more closely resembling the genetic architecture of chronotypes in natural human populations [44,45]. Hence, these populations provide a powerful tool for studying – A) correlations between chronotypes and sleep, asking not merely whether clock variation affects sleep, but whether sleep traits co-evolve when circadian timing is under selection, and B) how environmental inputs and perturbations modulate sleep of chronotype populations using specific experimental paradigms.

We performed comprehensive characterisation of various sleep parameters in experimentally evolved chronotype populations of *Drosophila melanogaster* using traditional sleep definition [46,47] as well as recently identified sleep states classification [48] to obtain a richer picture of sleep architecture and to test whether chronotype selection has altered the organisation of sleep states in both sexes. We then examined how sleep of these populations is modulated by varying two important environmental cues that impact sleep timing and quality – light, to which chronotypes are known to differ in sensitivity [49,50], and temperature, whose effects on sleep across chronotypes remain unexplored. Additionally, we examined if these chronotypes respond differentially to perturbations like sleep disruption and jetlag conditions to probe homeostatic differences that studies on humans have suggested but cannot isolate. We found that indeed sleep architecture showed correlated evolution with chronotype, primarily through changes in how sleep was organised, and *early* and *late* populations departed from *control* in distinct ways, with differences evident across both circadian and homeostatic aspects of sleep regulations.

## Methods

### Fly strains

#### Chronotype populations derived by long-term laboratory selection

All studies reported here were carried out on four replicate *early*_1-4_ and *late*_1-4_ populations that were originally were derived from common ancestral *control*_1-4_ populations by applying selection pressure on the timing of adult eclosion [41]. The selection protocol involves collecting flies emerging in a 4-hour morning window (ZT21 - ZT1) designated as *early* and flies emerging in 4-hour evening window (ZT9 - ZT13) designated as *late*. For *control* no selection for eclosion time was applied, i.e., flies where any phase of eclosion contributed to the gene pool were collected each generation (Figure S1A). All twelve of these populations have been maintained under identical conditions - Light-Dark cycles (LD 12:12), constant temperature of 25 ± 0.5⁰ C and 70% ± 5% relative humidity. Common subscripts across *early, control* and *late* represent common ancestry and are referred as blocks as part of our experimental and statistical hypothesis-testing design. The populations are maintained on a 21-day discrete generation cycle in large numbers (∼1200 flies, enabling random mating) on banana-jaggery food medium. Before each experiment, one generation of relaxation of selection pressure was provided to obtain standardized populations (also referred to as ‘common-garden’ conditions [51]. Progeny of these standardized populations were used as experimental flies to minimize maternal and any non-genetic inheritance effect on measured traits [51]. These populations continue to remain under active selection at the time of writing this manuscript, and all experiments described here have been carried out between generation numbers 363 to 420 of selection (see Supplementary Table S1 for details). Figure S1B represents percentage emergence of flies during morning and evening windows across generations. Generation-wise trajectories indicate continuous divergence of chronotype populations in eclosion timing.

### Sleep recording

Sleep recordings were carried out using Drosophila Activity Monitors (DAM systems by Trikinetics) system. Unless stated otherwise, ∼32 five-day old virgin males per population were singly housed in 5mm glass tubes with Sucrose-Agar medium (5% sucrose, 2% agar and benzoate) and placed in DAMs. The monitors were placed in Percival incubators (DR-36VL, Percival Scientific, Perry, USA) for the course of the study. Light intensity was adjusted using Lee filters over white, fluorescent lamps and was estimated using LI-COR luxmeter. All blocks were assayed under LD12:12, 500 ± 10 lux intensity, 25 ± 0.5°C and 70 ± 5% RH (standard lab conditions) for three to four days, unless mentioned otherwise. One day of acclimatization into respective environmental regimes was provided before the start of recording for each experiment. Data was recorded in 1-minute intervals. Details of each environmental regime are discussed below.

#### Baseline LD12:12 recordings

Both virgin males and virgin females were recorded under standard lab conditions for 3-4 days.

#### Light intensity experiments

Flies were recorded under different light intensities – 1, 10, 100, or 1000 lux (0.022±0.015, 0.113±0.08, 1.342±0.25, 9.152±1.37 W/m² respectively) during daytime under LD 12:12 cycles. Light intensity was adjusted using LED strips and Lee filters. Horizontal DAM IV monitors were used to ensure each fly receives uniform light intensity. Replicate blocks were assayed independently in separate experiments.

#### Temperature cycles experiments

Flies were recorded under temperature cycles with constant darkness (DDTC), 28°:18°C thermophase: cryophase. Sleep was quantified for 3 days post establishment of stable phase relationship with the zeitgeber.

#### Jetlag experiment

Flies were recorded under LD12:12 for 6 days before being subjected to either a 9 h phase advance or a 9 h phase delay. Recordings continued for 10 days following the shift. Light intensity during this experiment was maintained at 0.1 lux to minimize masking effects to bright light.

#### Sleep deprivation experiments

Flies were subjected to sleep deprivation for different durations using a vortexer (Multitube vortexer, VWR, Radnor, PA, USA) by securely clamping DAMs to the vortexer which rotates (Vortexer Mounting42 Plate, Trikinetics), at a force of 1-g (500 RPM, 3.6mm eccentric orbit). The vortexer was programmed using a light controller (LC4, Trikinetics) to provide 2-s long rotational stimuli every 3 min for 12 hours during nighttime, following baseline recordings under standard lab conditions. Undisturbed flies that did not receive any mechanical stimuli were also recorded in the same incubator. Replicate blocks were assayed independently in separate experiments.

### Sleep quantification and analysis

PhaseR, an R-based shiny app [52], was used to obtain sleep data (sleep per 30 mins) and to estimate the following sleep parameters-number of bouts, median bout duration (since bout duration is highly skewed), sleep latency and sleep duration during daytime and nighttime, using DAM text files. In all experiments, sleep was defined as inactivity of 5 mins or more (standard sleep definition), unless specified otherwise [46,47]. For analysis of sleep states, inactivity durations of 5 – 29 mins, 30 - 59 mins, and ≥ 60 mins were classified as short, intermediate and long sleep respectively [48]. Average across days and then across individuals for a population were calculated after removing data from dead individuals for each sleep parameter.

#### Probability of first sleep bout for each sleep state

To estimate which sleep state flies first enter after lights OFF, we identified, for each fly on each LD 12:12 cycles, the first sleep bout that began during the night (ZT12–24) and recorded its state. Bouts already in progress at lights-off were not counted. For each fly, the probability of each state was the number of nights on which the first bout was that state, divided by the total number of nights. Probabilities were averaged across flies within each replicate block and then across the four blocks. Analyses were performed in R using custom scripts (R version 4.6.1).

#### Midpoint of rise in sleep in DDTC

Night sleep onset phase was determined per individual using an inflection-point method using custom-written R code (R version 4.6.1, *signal* package). Sleep profiles generated using phase R for each sleep definition, were smoothed using a Savitzky-Golay filter (polynomial order 3, window length 7-time bins) to reduce noise while preserving the shape of the waveform [53]. The daytime sleep trough i.e., the point of minimum sleep within ZT 6–12 time window (ZT 0 – start of thermophase; ZT 12 – start of cryophase) was identified on the smoothed profile. First and second derivatives of the smoothed time series were computed numerically using finite differences [53]. The onset of the night sleep bout was defined as the first zero-crossing of the second derivative from positive to negative after the daytime trough up to ZT 18, corresponding to the inflection point of the rising limb where sleep was increasing at its maximum rate. In cases where no clean zero-crossing was detected, the time point of maximum first derivative in the post-trough rising window was used. Mean and SEM of phase of sleep onset were calculated across individuals for each population.

#### Change in sleep post jetlag

Change is all sleep parameters after shift was calculated by subtracting each fly’s average sleep parameter over the three pre-shift baseline days (Day −3 to Day −1) from its value on each day post-shift (Day 3 to Day1) under the new LD schedule [54]. Day 0 being the cycle in which shift occurred was excluded from analysis. The change from baseline was then averaged over individuals of a block.

#### Sleep rebound post sleep deprivation estimation

Sleep rebound was calculated in time windows - 0 to 12h, 12 to 24h and 0 to 24h post sleep deprivation-for both sleep deprived and undisturbed flies by subtracting post-deprivation sleep from baseline sleep in corresponding time duration. To correct for day-to-day variations, sleep rebound of each sleep deprived fly was further subtracted from average sleep rebound of undisturbed flies of the same population. This corrected sleep rebound was then averaged over individuals of a population. Sleep rebound using both standard sleep and long sleep definition were calculated for each time window.

#### Locomotor activity quantification

Raw activity counts were summed in 30 mins bin and plotted across the day to get activity profiles of chronotype flies. Each individual profile was averaged across over 4 days and then across individuals. Total activity counts during day, and night were calculated and compared across chronotype flies.

### Statistical analysis

Randomized block ANOVA model (Selection – fixed factor and block – random factor) was used to compare measured parameters across chronotypes using Statistica v.7 (StatSoft, Tulsa, USA) unless stated otherwise. Tests were performed separately for each sleep state. For light intensity experiments, light intensity was used as another fixed factor in the randomized block ANOVA model. For jetlag experiment, and for the number of bouts and median bout duration of sleep-deprived flies before (Day 1) and after (Day 3) deprivation, a linear mixed-effect model was used, with days as a repeated measure, in addition to block as random factor, and selection as fixed factor using a custom written R code (lme4/lmerTest and emmeans packages in R) and significance of fixed effects was tested by Type III ANOVA with Satterthwaite’s approximation for denominator degrees of freedom. All post-hoc multiple comparisons were made using Tukey’s HSD test. Significance level, α = 0.05 was used. Letters on top of bar plots represent differences across chronotypes unless mentioned otherwise in figure legends.

#### Effect sizes

Cohen’s *d* was calculated for each sleep parameter for *early - control* and *late - control* to measure effect sizes of *early* and *late* selection across replicate populations and replicate experiments. For each experiment, the four block-wise estimates were combined into an inverse-variance-weighted (IVW) average, which weights each estimate by its precision [55]. All calculations were done with custom R code. GraphPad Prism v.8 was used for all data plotting. For raster plots, custom written R code was used.

## Results

### Night sleep is more fragmented in both male and female flies of *early* chronotype

We asked whether sleep architecture has evolved in our artificially selected chronotype populations under controlled conditions. We comprehensively characterized multiple aspects of sleep architecture in both sexes under laboratory conditions similar to their rearing conditions (LD12:12, 25°C ± 0.5°C, 70% ± 5% RH). We measured sleep using standard sleep definition i.e., inactivity of 5 min or more [46,47]. Flies exhibit a mid-day siesta, and their nighttime sleep is known to be architecturally as well as functionally different from day sleep [56,57]. Therefore, we quantified sleep parameters for daytime and nighttime separately.

The sleep profiles of males were broadly similar between chronotypes (Figure 1A), with *late* chronotypes tending towards slightly lower sleep during day and *early* showing lower sleep during night consistently across replicate experiments (Table S2 and S3). However, we did not detect statistically significant differences in daytime or nighttime sleep duration (Figure 1B, Table S2A) using the standard definition for fly sleep. To assess sleep architecture, we calculated other parameters such as number of bouts, median bout duration, and sleep latency during daytime and nighttime for standard sleep. During nighttime, sleep latency of *early* flies was significantly shorter than *late* (Figure 1C), suggesting that *early* chronotype flies fall asleep sooner than *late* during night. Interestingly, *early* flies also showed more number of sleep bouts and shorter median bout duration i.e., more fragmented sleep than *control* during night while *late* showed an intermediate phenotype being statistically not different from either *early* or *control* (Figure 1C). Sleep fragmentation across night can be better visualized in representative raster plots where each row represents sleep architecture of individual flies on a given night (Figure S2B). These results suggest that selection of *early* chronotype led to correlated evolution of more fragmented sleep during nighttime with no difference in total sleep amount in males. Interestingly, no differences were found among the chronotypes in any of the sleep parameters during daytime (Figure S2A).

**Figure 1.**
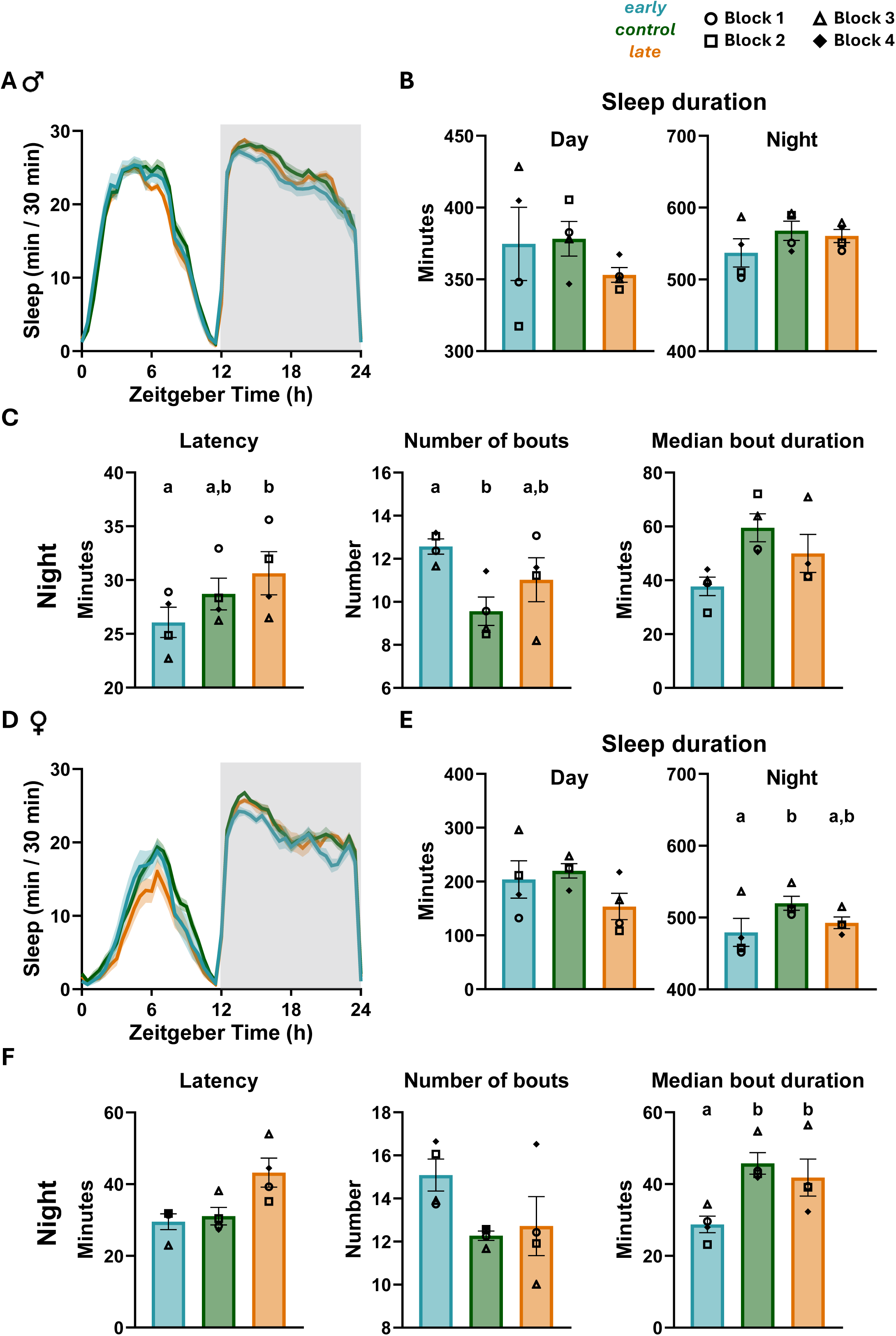
Sleep in male and female chronotype populations under light dark (LD) cycles. (A-C) Standard sleep of virgin male chronotype flies under LD 12:12– (A) Sleep profile; Zeitgeber time 0 represents lights ON. (B) Sleep duration during daytime (F_(2,6)_ = 0.53, *p* = 0.609) and nighttime (F_(2,6)_ = 2.03, *p* = 0.211). (C) Nighttime sleep parameters-latency (F_(2,6)_ = 8.04, *p* = 0.02), number of bouts (F_(2,6)_ = 6.73, *p* = 0.029) and median bout duration (F_(2,6)_ = 4.03, *p* = 0.077). (D-F) Same as A-C for virgin female chronotype flies. (E) Sleep duration during daytime (F_(2,6)_ = 1.97, *p* = 0.219) and nighttime (F_(2,6)_ = 7.17, *p* = 0.034). (F) Night sleep parameters - latency (F_(2,6)_ = 5.11, *p* = 0.051), number of bouts (F_(2,6)_ = 4.91, *p* = 0.054), median bout duration (F_(2,6)_ = 20.77, *p* = 0.002). All sleep profiles are averaged across 4 replicate populations, error bands are ±SEM across 4 populations and grey shaded region represents lights OFF. Bar plots show means ± SEM across 4 populations, with symbols representing individual blocks. A two-way randomized block ANOVA (selection as a fixed factor and block as a random factor) followed by Tukey’s HSD post hoc test, was used for statistical comparisons. Letters on top of bar plots represent statistically significant differences (*p <* 0.05). No letters are displayed in case of non-significant ANOVA results. ALT TEXT: Graphs showing standard sleep in male (A-C) and female (D-F) chronotype flies under LD cycles. Panels A and D show 24h sleep profiles and bar plots show day and night sleep duration and nighttime sleep latency, number of bouts and median bout duration. *late* chronotypes have longer night sleep latency than *early. early* flies show more number of bouts and shorter median bout duration during night than *control* suggesting greater night sleep fragmentation.

As fly sleep is known to be sexually dimorphic [57], we then asked whether the differences between chronotypes seen in males were also present in females (Figure 1D). Female flies broadly recapitulated all the differences. During day, *late* flies showed a trend of reduced sleep duration with no difference in any other sleep parameter (Figure 1D and S2A). During night, *early* flies slept significantly less than *control* and had a significantly shorter median bout duration with a trend of increase in number of bouts (Figure 1E, 1F, S2B and Table S2B). Trend of increase in night sleep latency of *late* flies than others was also observed (Figure 1F). This suggests that effect of selection on sleep is qualitatively consistent across both sexes and across replicate experiments (Table S3).

Since there was no difference in waveform or number of activity counts during day or nighttime among the chronotypes, these differences are specific to sleep organization rather than a reflection of changes in locomotion (Figure S2C, Table S4).

### *early* flies show shift in sleep states organisation during night

Recent reports characterising fly sleep have identified distinct sleep states based on both physiological and behavioural signatures [48,58–62]. More recently, Abhilash et al. 2026, reported that locomotor activity counts when binned appropriately can represent biologically meaningful states that differ in how strongly they are shaped by the circadian clock and the sleep homeostat [48]. We used this approach and estimated short, intermediate or long sleep based on inactivity duration (see methods). A detailed characterisation of the time spent in these states and the latency of each state provided a richer picture of sleep architecture and allowed us to test whether chronotype evolution has altered the organisation of sleep states.

The sleep profiles of each state for male flies suggested that at night, short and intermediate sleep was higher for *early* chronotypes whereas long sleep was lesser (Figure 2A). This difference was observed throughout the night. For *early* chronotypes, the duration of short sleep was significantly higher than *control* while long sleep was significantly lesser (Figure 2B, Table S5). Consistent with this result they also tended to have more number of short and intermediate sleep bouts (Figure S3A, Table S5). Together these results suggest that *early* flies exhibit robust alteration in night sleep architecture with redistribution of sleep towards short, more fragmented bouts at the expense of long consolidated bouts. *late* flies show less distinct changes in sleep architecture although they tend in the same direction as *early* flies.

**Figure 2.**
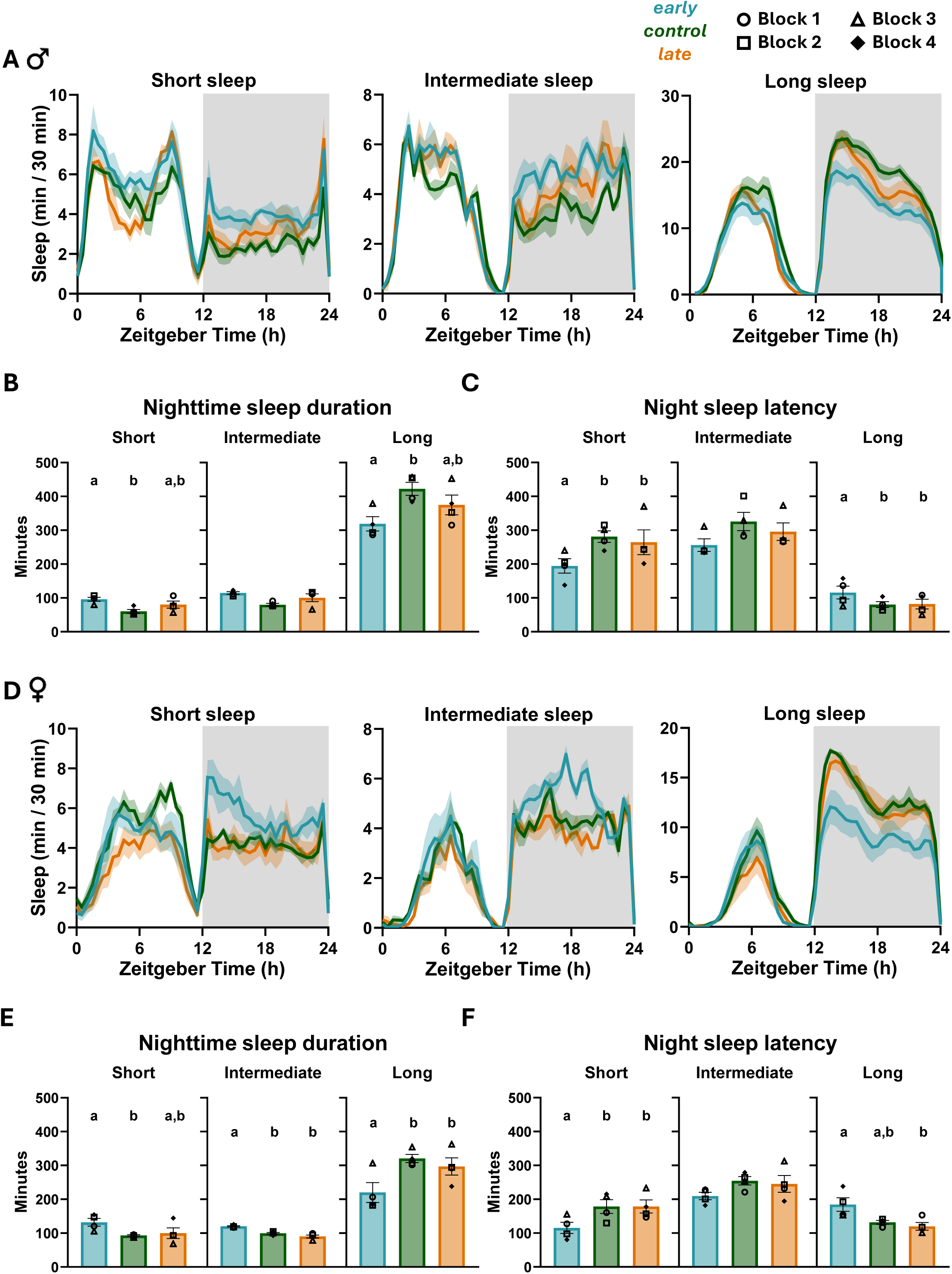
Sleep states in male and female chronotype populations under LD cycles. (A-C) Sleep states of virgin male chronotype flies under LD cycles – (A) Short (left), Intermediate (middle) and Long (right) sleep profiles, Zeitgeber time 0 represents lights ON. (B) Nighttime sleep duration of each sleep state-Short (*F_(2,6)_* = 8.66, *p* = 0.017), intermediate (*F_(2,6)_* = 4.85, *p* = 0.056) and long sleep (*F_(2,6)_* = 13.54, *p* = 0.006). (C) Nighttime sleep latency for each sleep state-Short (*F_(2,6)_* = 8.67, *p* = 0.017), intermediate (*F_(2,6)_* = 2.98, *p* = 0.126) and long sleep (*F_(2,6)_* = 7.71, *p* = 0.022). (D-F) Same as A-C for virgin female chronotype flies. (E) Short (*F_(2,6)_* = 6.93, *p* = 0.028), intermediate (*F_(2,6)_* = 21.79, *p* = 0.002) and long sleep (*F_(2,6)_* = 17.16, *p* = 0.003). (F) Short (*F_(2,6)_* = 7.24, *p* = 0.025), intermediate (*F_(2,6)_* = 1.96, *p* = 0.22) and long sleep (*F_(2,6)_* = 5.60, *p* = 0.042). All sleep profiles are averaged across 4 replicate populations, error bands are ±SEM across 4 populations and grey shaded region represents lights OFF. Bar plots show means ± SEM across 4 populations, with symbols representing individual blocks. A two-way randomized block ANOVA (selection as a fixed factor and block as a random factor) followed by Tukey’s HSD post hoc test, was used for statistical comparisons. Analyses were performed separately for each sleep state. Letters on top of bar plots represent statistically significant differences (*p <* 0.05). No letters are displayed in case of non-significant ANOVA results. ALT TEXT: Graphs showing short, intermediate and long sleep states in male (A-C) and female (D-F) chronotype flies under LD cycles. Panels A and D show 24h sleep profiles for each sleep state, and the bar plots show nighttime sleep duration and latency. In both sexes, *early* flies have more short sleep and less long sleep at night than *control*. *early* flies also show less latency for short sleep and more latency for long sleep as compared to other chronotypes.

**Figure 3.**
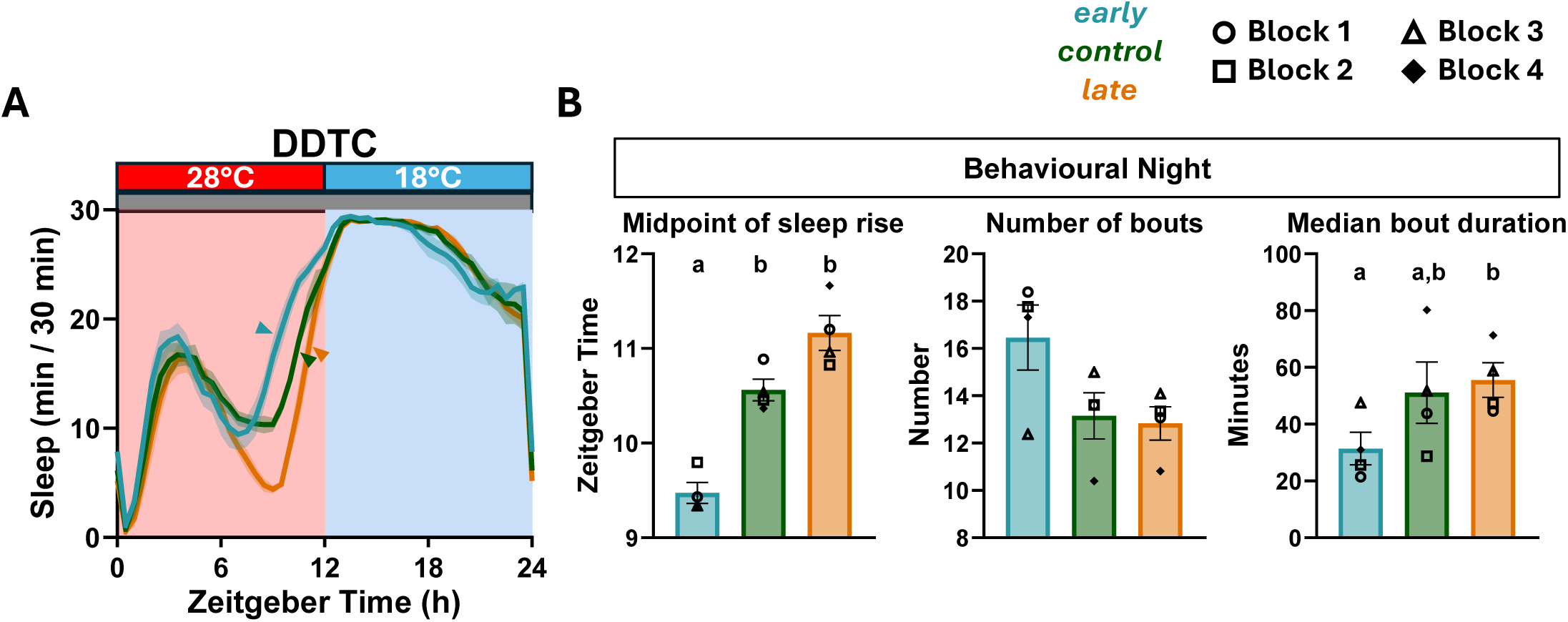
Effect of temperature cycles on sleep in fly chronotype populations. (A) Sleep profile of male chronotype flies under DDTC (temperature cycles in constant darkness; 28 °C thermophase, 16 °C cryophase), Zeitgeber time 0 represents onset of thermophase. Y-axis is sleep per 30 min. Arrowheads indicate the midpoint of rise in sleep for each chronotype, used as an estimate of behavioral night onset. (B) Sleep parameters during behavioral night under DDTC-Midpoint of rise in sleep (*F_(2,6)_* = 28.5, *p* < 0.001), Number of bouts (*F_(2,6)_* = 3.15, *p* = 0.115) and Median bout duration (*F_(2,6)_* = 5.34, *p* = 0.046). Sleep profiles is averaged across 4 replicate populations, error bands are ±SEM across 4 populations. The grey bar above the sleep profile represents constant darkness; red and blue bars and shaded regions represent thermophase and cryophase, respectively. Bar plots show means ± SEM across 4 populations, with symbols representing individual blocks. A two-way randomized block ANOVA (selection as a fixed factor and block as a random factor) followed by Tukey’s HSD post hoc test, was used for statistical comparisons. Letters on top of bar plots represent statistically significant differences (*p <* 0.05). ALT TEXT: Graphs showing sleep in fly chronotype populations under temperature cycles (DDTC). Panel A shows the sleep profile, where sleep rises before the switch to cold phase, earliest in *early* flies and latest in *late* flies. Panel B show bar plots of midpoint of sleep rise, number of bouts and median bout duration during behavioural night where *early* flies have an advanced midpoint of sleep rise and sleep in shorter bouts than *late* flies.

Additionally, we quantified night sleep latency for each sleep state (Figure 2C, Table S5). Consistent with the previous report [48], long sleep had shortest latency at night suggesting that it is the first state that occurs during night. Interestingly, *early* chronotype flies had shorter latency for short sleep and longer latency for long sleep as compared to other chronotypes (Figure 2C). These state-specific differences raised the question of whether the shorter, standard sleep latency observed in *early* flies (Figure 1C) reflects differences in the type of sleep state initiated at sleep onset. We therefore asked which sleep state has higher probability to occur first during night for each chronotype. We found that *early* flies showed similar probability of initiating sleep with either a short sleep bout or long sleep bout in males, whereas *control* and *late* were more likely to initiate sleep with long sleep bouts (Figure S3C). Thus, the shorter overall latency observed for *early* flies may reflect a shift in initial sleep state. Similar results were observed for female flies (Figure 2D-F, S3 and Table S5).

Interestingly, we did not observe any chronotype specific changes for any sleep state parameter for either sex during day (Table S6). Together, these results suggest that selection of chronotypes led to shifts in how sleep states are organised particularly during the night and that this effect occurred independent of sex.

### High light intensity delays night sleep across chronotypes

Light is a dominant time cue that affects sleep timing and human sleep chronotypes are reported to achieve differential light exposure and have differential photosensitivity [6,22,39]. Our fly chronotype populations have also been demonstrated to exhibit differential photosensitivity [49,50]. This led us to ask whether chronotypes show differential light responses in terms of sleep properties, using a range of light intensities-1 lux, 10 lux, 100 lux and 1000 lux under LD12:12 conditions.

We first asked how light intensity affects sleep in *control* flies, as to the best of our knowledge, no study has reported this. We found that daytime sleep duration increases with increase in light intensity in *control* (Figure S4, top, green curve). This matches earlier work that reported longer midday siesta when light intensity is increased [63]. We also observed an increase in night sleep latency with increase in light intensity (Figure S4, green curve) suggesting that high intensity during day leads to delay in night sleep. Next, we asked if chronotypes respond differently to increase in light intensity. No significant interaction effect was detected for any sleep parameter, suggesting all chronotypes responded similarly to increase in light intensity (Figure S4, Table S7). Only day sleep duration and night sleep latency increased steadily with intensity. A few other parameters like daytime sleep latency (shortest at 10 lux and longest at 1000 lux) and number of bouts (lowest at intermediate intensities) showed minor, non-monotonic variation across intensities. Night sleep duration also showed a small, non-monotonic dip at the higher intensities. Sleep state specific analysis did not reveal any chronotype-dependent differences across intensities. Overall, light intensity mainly influenced how much flies slept during the day, and how quickly they fell asleep at night with similar trends across *early, control* and *late* flies.

### Night sleep latency difference across chronotype flies increases under temperature cycles

Temperature is another important time cue especially for poikilotherms like *Drosophila* and it is known to modulate sleep timing and quality in both flies and humans [64–66]. This is especially relevant since our fly chronotype populations have been demonstrated to have co-evolved differential thermosensitivity such that in presence of temperature cycles (warm Thermophase: cool Cryophase; TC) the extent of chronotype (phase) divergence between *early* and *late* populations is enhanced for both eclosion rhythm and locomotor rhythm parameters [67–70]. However, it has been unclear whether temperature cycles also differentially influence sleep of these chronotypes.

We measured sleep parameters under temperature cycles of 28°:18°C without any light cue (DDTC). Under this condition, we observed that the second prolonged sleep period which continued into the cold phase (cryophase or night) was initiated several hours before the transition / temperature dip at ZT12 (Figure 4A). Since the conventional measurement of night sleep latency was not appropriate here, we estimated the midpoint of rise in sleep by first estimating the inflection point where sleep levels increased sharply for each of the chronotypes (see methods). We found that *early* had significantly advanced midpoint of rise in sleep than both *control* (by ∼1.08 hour) and *late* (by ∼1.68 hour) (Figure 4B, Table S8). This suggests that sleep timing differences enhance under temperature cycles when masking effect of light is not present. We also applied this method to calculate sleep timing on LD dataset for consistency and the results were consistent with sleep latency results obtained by the conventional method i.e., *early* had shorter latency than *late* although marginally significant (Table S9).

**Figure 4.**
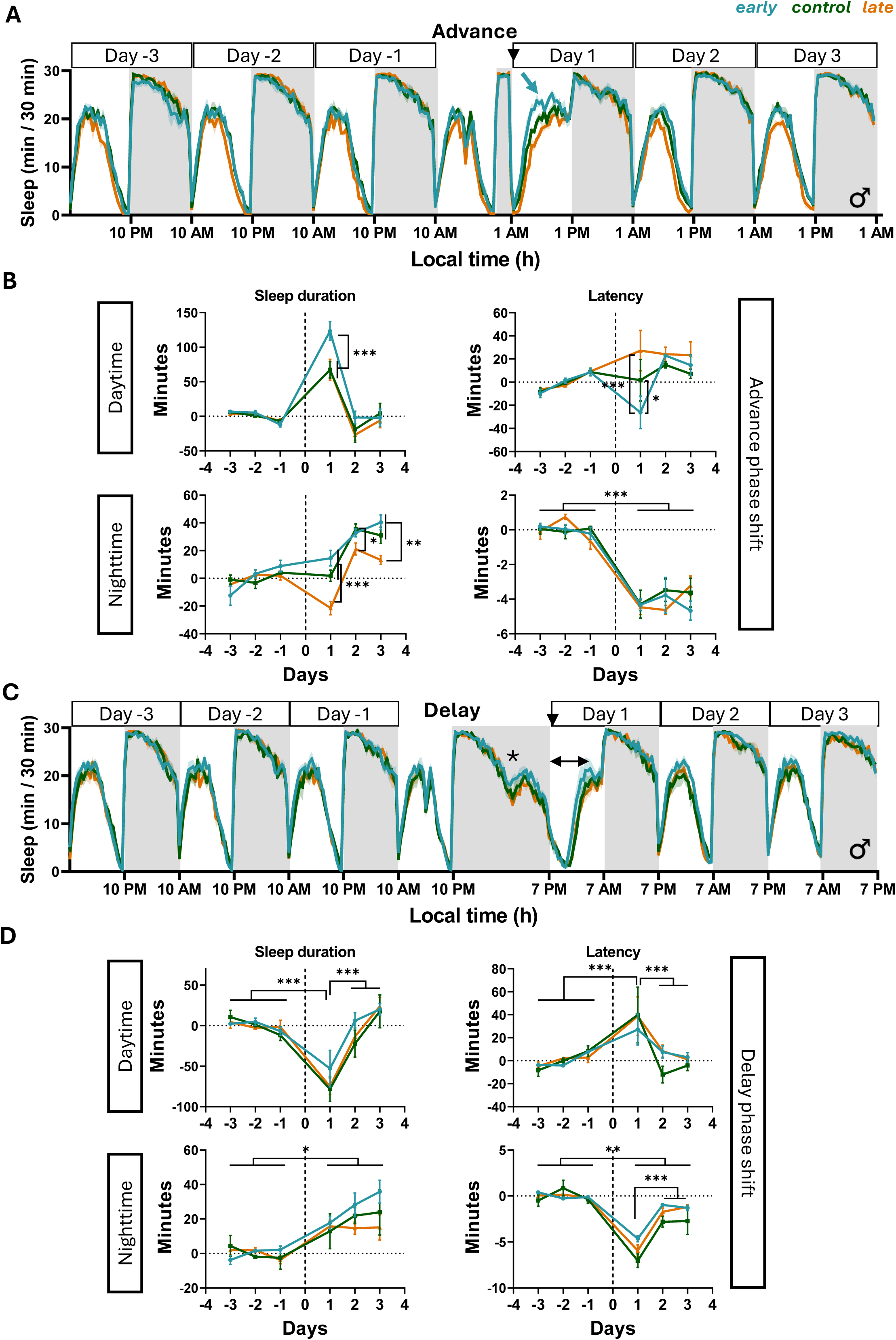
Effect of simulated jetlag on sleep in fly chronotype populations. (A) Standard sleep profile of male chronotype populations; the black arrowhead indicates the 9h advance phase shift of the LD cycle and blue arrow indicates increased sleep of *early* flies on first day (Day1) post shift. (B) Change from each fly’s baseline mean in daytime sleep duration (top left), daytime sleep latency (top right), nighttime sleep duration (bottom left) and nighttime sleep latency (bottom right) post advance shift. (C) Same as A for the 9h phase delay; the asterisk indicates attenuation in sleep due to the LD transition as per the previous unshifted LD regime and double arrowhead represents the delay in sleep latency post shift. (D) Same as B for the delay. Sleep profiles are averaged across 4 replicate populations, error bands are ±SEM across 4 populations and grey shaded rectangles represent lights OFF. X-axis is local time and Y-axis is sleep per 30 min. Days −3 to −1 are baseline cycles and Days 1–3 are cycles after the shift; Day 0 (dashed line), the cycle during which the shift occurred, was not analyzed. Error bars in line plots are ±SEM across 4 populations. A linear mixed-effects model with selection as a fixed factor, day as a repeated-measures factor and block as a random factor, followed by Tukey’s HSD post hoc test, was used for statistical comparisons. Asterisks between lines on a given day indicate significant differences among selection lines on that day, shown where the selection × day interaction was significant. Asterisks above the plots indicate significant differences between days, shown where only the effect of day was significant. \**p* < 0.05, \*\**p* < 0.01, \*\*\**p* < 0.001. Full model results are given in Table S8 and S9. ALT TEXT: Graphs showing sleep in fly chronotype populations before and after a 9h phase advance (A, B) and a 9h phase delay (C, D) of the LD cycle. Panels A and C show sleep profiles across the three baseline and three post-shift days. Panels B and D show line plots of the change from baseline in daytime and nighttime sleep duration and latency, where *early* flies show the largest increase in daytime sleep on the first day after the advance.

We then defined behavioural night under DDTC based on mid-rise time of second prolonged sleep period of the day for each chronotype and measured sleep parameters accordingly. Greater sleep fragmentation, i.e., increase in bout number (trend) and decrease in median bout duration was still observed in *early* flies with comparable magnitudes as seen under only LD cycles (Figure 4B, Table S8). We also estimated sleep parameters during 12h of thermophase and cryophase and found no difference in any sleep parameters except cryophase sleep latency being longer in *late* than *early* (Figure S5A, Table S8). Additionally, we characterized sleep state profiles under DDTC (Figure S5B) which have not been reported previously. During thermophase, a stark reduction of long sleep was observed as compared to cryophase, suggesting that temperature may influence long sleep state. Interestingly, in contrast to LD cycles, *early* flies now tended towards earlier onset of long sleep at night as compared to others (Figure S5B, Table S8). Taken together, these results suggest that light and temperature cycles modulate sleep timing of chronotype populations in distinct ways, with temperature cycles producing an exaggerated chronotype divergence.

### Chronotype specific sleep changes occur after advance jetlag

Human chronotypes are known to exhibit differences in recovery of sleep timing post jetlag [54]. In the contemporary world, larger fractions of the workforce are subjected to jetlagged conditions, which includes long distance air-travel and shift work to match global economies in addition to local needs. Jetlag profoundly disrupts circadian rhythms and sleep architecture [71]. A variety of confounding factors including travel schedules and personal obligations may obfuscate inferences regarding how chronotypes achieve sleep recovery post jetlag [54]. Thus, the question remains as to how advance, and delay shifts may differentially alter sleep architecture of chronotypes. We imposed identical phase shifts on our artificially selected chronotype populations with matched genetic background and environment using light/dark cycles and characterized sleep architecture in detail.

Under 9h phase-advance jetlag conditions, flies experienced an abrupt lights-ON transition during their subjective night (Figure 4A black arrowhead), resulting in immediate awakening and disruption of their night sleep. Subsequently flies returned to sleep during the advanced daytime. For all three chronotypes, daytime sleep directly transitioned into night sleep, lacking the characteristic evening decline in sleep associated with high evening activity of flies. By Day 2 post-shift, sleep profiles of all populations resembled baseline conditions. We quantified changes in sleep parameters upon jetlag for three days and whenever a sleep parameter returned to its baseline level, we considered it as recovered (see methods). We found that regardless of chronotype, flies show an increase in sleep during daytime on Day 1 post shift (Figure 4B; Table S10). This increase in sleep is possibly a result of sleep disruption during the night of transition. The chronotypes differed in the magnitude of this response, with *early* flies showing almost twice as large an increase in sleep duration as compared to the others (Figure 4B, top left). Chronotype-specific response is observed for daytime sleep latency as well with *early* flies showing greatest reduction in day sleep latency, post shift (Figure 4B, top right). Daytime sleep latency results together with sleep duration suggest that *early* flies show higher magnitude of compensation to recover to baseline level as a response to advance jetlag. We also observed that day sleep latency is not significantly altered post shift as compared to baseline days for *control* and *late,* although qualitatively we see an increase in latency for *late* flies suggesting an opposite effect of chronotype on sleep latency (Table S10). During night, a chronotype specific response is observed for sleep duration (Figure 4B, bottom left). Advance shift reduces night sleep in *late* flies while *early* and *control* are unchanged as compared to baseline (Table S10B). In contrast, all chronotypes show a similar reduction in night sleep latency post advance shift which does not recover to baseline even till Day 3 post shift.

In contrast to 9h phase advance paradigm, 9h phase delay shift indicated by black arrowhead in Figure 4C, caused prolonged dark phase which caused the flies to remain asleep for longer with slight attenuation during subjective day transition as per previous unshifted LD cycle (shown by asterisk, Figure 4C). Consequently, flies remain active for longer, post lights-ON (shown by double arrowhead, Figure 4C), increasing daytime latency and falling asleep during nighttime as per previous unshifted LD cycle. Flies transitioned directly from delayed daytime sleep into subsequent night sleep without exhibiting the characteristic evening dip in sleep similar to advance jetlag regime. Day 2 onwards post shift, sleep profiles resembled baseline conditions. Upon quantification, we found that post delay shift, day sleep duration decreases and day sleep latency increases (Figure 4D, Table S11). As expected, these changes are in opposite direction as compared to advance phase shift. An increase in night sleep duration is observed post delay shift which does not return to baseline levels till Day 3, whereas night sleep latency decreases on Day 1 and returns to baseline by Day 2. Interestingly, unlike advance, all chronotypes are affected similarly by the delay phase shift.

To characterise the sleep quality further, we measured change in bout architecture due to phase shift. An overall increase in bout number and decrease in median bout duration during daytime is observed post advance phase shift which does not quickly return to baseline levels suggesting sleep fragmentation (Figure S6A, top). During night also we see an overall increase in bout number and decrease in median bout duration but only on Day 1 post shift. On Day 2 post shift, the opposite effect is observed i.e., decrease in bout number and increase in median bout duration suggesting that sleep is fragmented on Day 1 post jetlag but consolidates on Day 2.

Although *early* flies qualitatively do not show the same alterations as *control* and *late*, no significant effect of interaction was detected. Similar to advance shift, sleep fragmentation during daytime is observed post delay shift also, which does not return to baseline levels (Figure S6B). However, no significant change in bout parameters is observed during nighttime. Together, these results demonstrate that sleep responses to circadian misalignment depend on the direction of phase shift, with advance shifts eliciting chronotype-specific changes in sleep duration and latency, whereas delay shifts produce largely similar responses across chronotypes. Both shifts additionally induce persistent daytime sleep fragmentation, suggesting that recovery of sleep timing and duration does not necessarily coincide with recovery of sleep architecture.

### Upon sleep deprivation, *late* chronotype flies show less sleep rebound at night

Sleep loss increases sleep pressure, inducing the sleep homeostat to restore sleep-wake balance [72,73]. Lost sleep is compensated by recovery sleep or rebound which predominantly comprises of deep consolidated sleep [48,57,74,75]. We hypothesised that our chronotype populations may differ in the extent and quality of rebound sleep, and that it would indicate that sleep homeostasis has co-evolved in response to selection for divergent eclosion chronotypes. To assess this, we sleep deprived flies for 12h during night and recorded sleep for subsequent recovery period, alongside undisturbed flies (Figure 5A and 5B).

**Figure 5.**
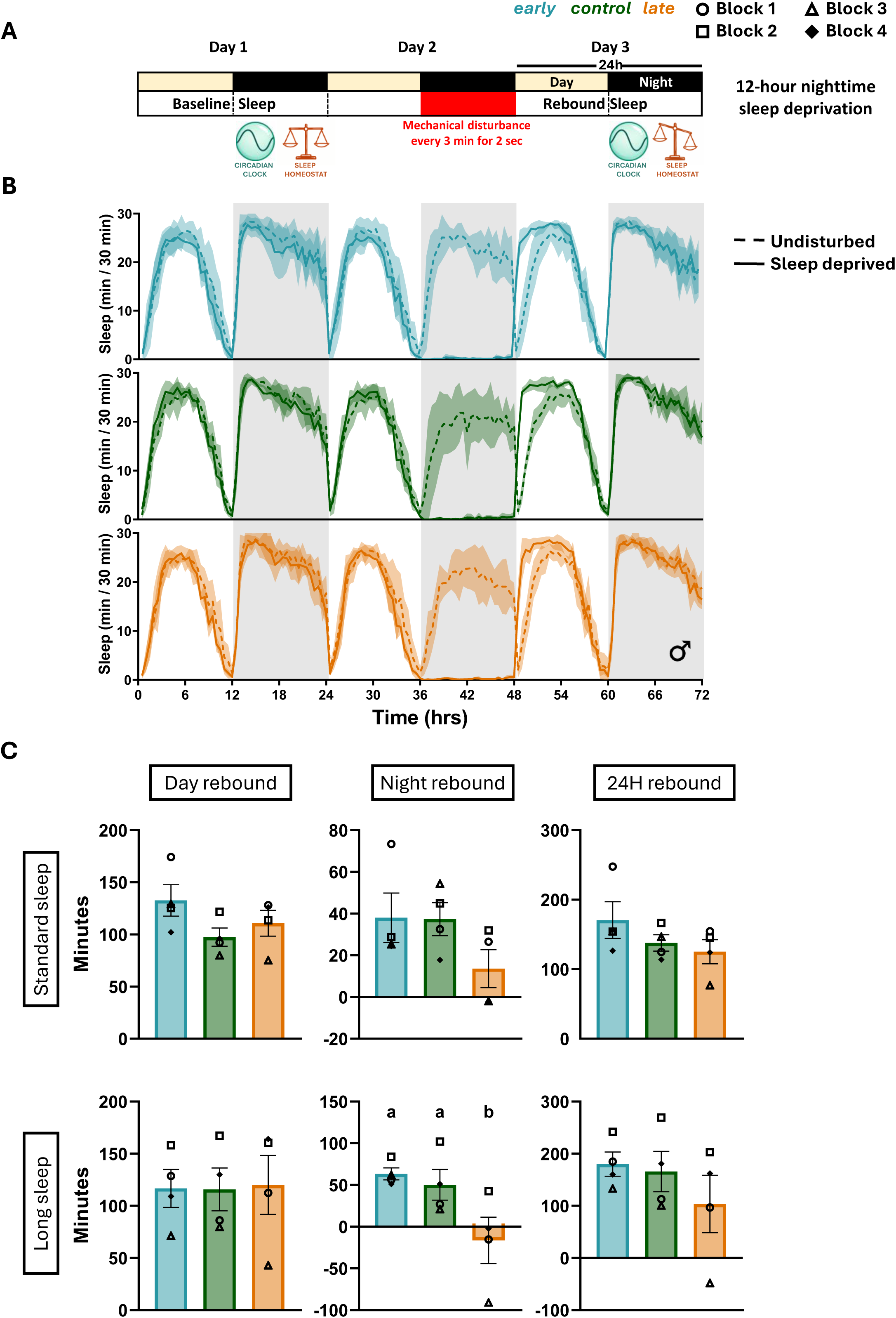
Sleep rebound after night sleep deprivation in fly chronotype populations. (A) Schematics demonstrating sleep deprivation regime. Yellow and black rectangles represent light and dark phase of LD cycle respectively. Red rectangle represents 12h of mechanical deprivation of sleep (2s disturbance every 3 min) during the night of Day 2; Day 1 served as baseline day and Day 3 as the rebound day post deprivation. (B) Sleep profiles of male *early* (top), *control* (middle) and *late* (bottom) flies across 3 days; time 0 represents lights ON on Day 1. Dashed lines represent undisturbed flies, and solid lines represent sleep-deprived flies. Profiles are averaged across 4 replicate populations, error bands are ±SEM across 4 populations and grey shaded regions represent lights OFF. (C) Sleep rebound quantification using standard sleep definition (top) - day (*F_(2,6)_* = 2.36, *p* = 0.174), night (*F_(2,6)_* = 2.47, *p* =0.164) and 24h rebound (*F_(2,6)_* = 1.69, *p* = 0.261) and the long sleep definition (bottom) - day (*F_(2,6)_* = 0.03, *p* = 0.965), night (*F_(2,6)_* = 10.41, *p* = 0.011) and 24h (*F_(2,6)_* = 3.24, *p* = 0.111). Day, night and 24h rebound represent rebound during 0-12h window, 12-24h window and 0-24h windows post deprivation, respectively. Bar plots show means ± SEM across 4 populations, with symbols representing individual blocks. A two-way randomized block ANOVA (selection as a fixed factor and block as a random factor) followed by Tukey’s HSD post hoc test was used for statistical comparisons. Different letters on top of bar plots represent statistically significant differences (*p* < 0.05); no letters are shown where the ANOVA was not significant. ALT TEXT: Schematics and graphs showing response of male chronotype flies to 12h of sleep deprivation during night. Panel A shows deprivation schedule, panel B shows 3-day sleep profiles of deprived and undisturbed flies and panel C shows bar plots of day, night and 24h sleep rebound using standard and long sleep definitions. Standard sleep rebound does not differ across chronotype populations, but *late* flies show reduced long sleep rebound during night (ZT12-24) after deprivation than other chronotypes.

Sleep rebound was detected and quantified for different time windows - day (0 – 12 h), night (12 - 24h) and 24h (0 - 24h) post deprivation (see methods for details). Using the standard definition of sleep, the magnitude of rebound did not differ significantly among chronotypes, indicating comparable homeostatic responses (Figure 5C; Table S12). We also quantified the long sleep rebound as it is considered a better reflection of sleep rebound (Figure S7A) [48]. We found that upon estimation of rebound using the window 0 – 12 h post deprivation, all chronotypes showed similar degree of long sleep rebound. However, during the subsequent night (12-24h) post deprivation, *late* showed significantly less long sleep rebound than *early* and *control* during night (12-24h) post deprivation (Figure 5C; Table S12). We also quantified rebound for other sleep states and found that *early* flies showed negative rebound for intermediate sleep during night period. (Figure S7B). This suggests *early* flies show increase in long, homeostatic sleep during rebound at the expense of intermediate sleep, whereas *control* flies do not show this trade off and simply increase long sleep. No difference in any other time window or short sleep across chronotypes was observed (Table S12). Thus, upon sleep deprivation using mechanical stimuli, *late* flies show lower long sleep recovery, and the observed altered sleep homeostasis is dependent on time-of-day. Additionally, we observed that *early* flies continued to show more number of sleep bouts and shorter median bout durations on the night following deprivation (Figure S7B, Table S13), indicating that their fragmentation phenotype persisted under elevated sleep pressure.

## Discussion

### Artificial selection of chronotype flies leads to correlated evolution of sleep architecture

Human studies have reported variable associations between chronotype and sleep, making it difficult to establish whether differences in sleep architecture are intrinsically associated with chronotype or arise from differences in genetic background, environmental exposure and lifestyle [16–21,23,24]. The experimentally evolved *Drosophila melanogaster* chronotype populations provided an opportunity to examine this relationship under controlled conditions. Here we present evidence that artificial selection on the circadian clock driven timing of adult emergence resulted in correlated evolution of sleep architecture. The *early* and *late* chronotype flies exhibited diverged sleep timing at night mimicking human sleep chronotypes. Interestingly, the most consistent and robust differences in sleep of chronotype populations was in how their sleep was organised. *early* flies exhibited more fragmented nighttime sleep whereas *late* flies showed weaker and less consistent changes in the same direction as *early*. These findings suggest that chronotypes are indeed correlated with differences in sleep architecture and this correlation is detectable even in the absence of the social and environmental confounds that complicate interpretation of human chronotype-sleep associations. Additionally, using this long-term laboratory selection approach we demonstrated that chronotypes and sleep are evolutionarily linked potentially through shared genetic mechanisms.

### Effect of chronotypes on sleep timing

Our study mainly focused on three features of sleep – latency or sleep onset timing, sleep fragmentation and sleep duration and asked how these parameters vary across chronotypes as well as how distinct environmental factors contribute to their manifestation. In humans, chronotypes are typically associated with timing of sleep, and our fly chronotype model successfully recapitulated this feature, i.e., *early* fly chronotypes had shorter sleep latency than *late.* This divergence was enhanced further under temperature cycles when masking effects of light was not present. In humans, a few studies have looked at effect of temperature cycles or fluctuations on sleep, as thermoregulation plays an important role in sleep timing [76,77]. Zhang et al. (2025) reported an association between thermoregulatory changes and shorter sleep latency in humans, although chronotype-dependent responses were not investigated [76]. Interestingly, we observed a sleep-state specific difference in sleep latency in fly chronotype populations which was dependent on environmental conditions. During the night, under LD, *early* took longer time to initiate long sleep, which is considered a better reflection of homeostatic sleep, instead, *early* flies initiated sleep sooner with short sleep bouts. Under TC however, this relationship was almost reversed such that *early* flies now exhibited long sleep sooner than others. This suggests that light and temperature cycles modulate long sleep timing of chronotype populations in distinct ways, potentially reflecting differences not only in circadian clock and homeostat interactions but also how these processes are regulated by distinct environmental inputs across chronotypes.

### Effect of chronotypes on sleep fragmentation and duration

While differences in sleep timing were expected given our divergent chronotypes, it is not intuitive that it could also cause changes in sleep architecture beyond timing. In our study, across replicate populations, we find that *early* flies exhibited increased night sleep fragmentation. As sleep maintenance is typically a function of sleep homeostasis, one possible explanation for more fragmented sleep in *early* flies is lower sleep pressure. Studies have reported that faster circadian clocks are associated with faster decay of sleep pressure [16,78]. While this is a possibility, we observed that even under elevated sleep pressure post sleep deprivation, night sleep fragmentation of *early* flies persisted, suggesting that the fragmentation is not a direct indication of low sleep pressure. Moreover, *late* flies which have slower circadian clocks also show changes in similar direction as *early*, albeit weaker and less consistent. This suggests that altered sleep architecture may reflect mechanisms beyond sleep-pressure dynamics alone.

Sleep quality is also known to be influenced by the circadian clock. Studies in rodents and flies have demonstrated that animals with mutation in clock genes exhibit greater sleep fragmentation [79–81]. Tabuchi 2018 proposed that a subset of circadian pacemaker neurons in flies show regular firing patterns at night that reduces firing rate of downstream arousal-promoting neurons which improves sleep consolidation and sleep quality at night relative to daytime when the firing pattern is irregular [82]. Electrophysiological characterisation of these populations could further establish their utility as a model for investigating the evolution of circadian regulation of sleep architecture.

Furthermore, we have observed that upon disrupting the clock using constant light (which makes flies arrhythmic), the previously observed fragmentation differences between chronotype flies were abolished (data not shown). This suggests that altered circadian clock of these diverged chronotype populations might contribute to greater sleep fragmentation during night. However, we remain cautious about this inference considering that constant light causes overall increase in fragmentation across populations, potentially producing a ceiling effect that may obscure chronotype differences.

While differences in sleep duration were not apparent using standard sleep definition (Inactivity of 5 min or more), sleep state specific analyses revealed that *early* flies exhibit less long sleep and more short sleep. Although these inactivity-based states are not directly shown to correspond to a particular sleep depth, studies in *Drosophila* have reported that increase in inactivity duration correlates with increase in sleep depth [48,56,60,74]. Hence, short and long sleep can potentially correspond to light and deep sleep respectively. To assess the extent to which these changes in sleep architecture parallel those reported in human chronotypes, we compared our findings with studies examining sleep architecture across human chronotypes. Mongrain et al. (2005) found morning chronotype men exhibit lower sleep efficiency and spend longer time in lighter sleep stages using polysomnography (PSG) approach [16]. Similarly, Kerkhof, (1991) reported decline in deep sleep stages in morning types using electroencephalography (EEG) [23]. More recently, Colelli et. al., 2023 showed that morning types were significantly associated with increase in rapid eye movement (REM) sleep using PSG, [24] while Merikanto et. al., 2017 showed association of lower sleep spindle amplitude and intensity in morning types using EEG [83]. Despite differences in specific sleep parameters affected, our findings share parallels with human studies reporting altered sleep architecture in morning chronotypes. Notably, our *late* flies also departed from *control,* though less strongly, in same direction as *early*, which suggests that any divergence from *control* in either direction can lead to correlated changes in sleep architecture. However, the fitness consequences of sleep differences remain unexplored in the current study. Previous studies on these populations (at ∼ 250 generations of divergence) have shown no differences in lifespan or fecundity among the chronotypes [84], however other physiological markers such as immune responses, reactive oxygen species (ROS) levels, or cognitive functions such as memory consolidation which are directly linked with sleep should be explored.

While most of our reported changes have been for nighttime sleep, *late* flies also showed a consistent trend of reduction in day sleep (Table S3). This distinguishes our findings from those reported for *period* gene mutants, where altered clock speed produces shifts in the evening activity peak, with short period mutant *per^S^*advancing it leading to less daytime sleep while long period mutant *per^L^* delaying it into the night leading to increase in daytime sleep [85,86]. No such phasing differences are apparent in our selected chronotype populations under LD12:12 (Figure S2). Notably, unlike *per^L^*, *late* flies exhibited a trend of reduced daytime sleep consistently, suggesting that sleep phenotype has evolved independently of activity. The reduction in day sleep of *late* flies could be a consequence of their high levels of Pigment Dispersing Factor (PDF) neuropeptide as reported previously in Nikhil et. al., 2016 [42]. PDF is a major synchronising factor in the fly circadian circuit and studies have shown that *pdf* null flies have increased daytime sleep as PDF release suppresses sleep and promotes wakefulness during day [87,88].

Additionally, we observed similar differences in sleep architecture across chronotypes in both males and females, although effects were more consistent across replicate experiments and numerically larger in males (Supplementary Table S3). Human studies have reported mixed findings regarding influence of sex on associations between chronotypes and sleep architecture. Mongrain et. al., 2005 has shown that men exhibit greater chronotype-dependent differences in sleep architecture using PSG, [16] whereas Raman and Coogan, 2019 using subjective questionnaires reported that sex does not moderate association between sleep quality and chronotype [89]. Together, our findings suggest that selection for divergent chronotypes can lead to correlated evolution of sleep architecture, extending beyond differences in sleep timing, to its maintenance and organisation. The broadly similar patterns observed across sexes further support the robustness of these evolved differences.

### Chronotype-dependent responses to sleep perturbations and circadian misalignment

We examined if sleep homeostasis itself has co-evolved in our chronotype populations using sleep rebound post sleep deprivation, a commonly used marker of sleep homeostasis. A recent study has shown that long sleep is a better reflection of homeostatic sleep [48]. We found that while *early* flies had to reduce intermediate sleep to show similar increase in long sleep as *control* during recovery period post deprivation, *late* flies showed a significant reduction in long, homeostatic sleep rebound in a time-of-the-day dependent manner. Reduction in rebound sleep could be due to – (a) lower homeostatic sleep need or (b) inability to recover required sleep need. Since the differences are observed only during night, it is possible that they stem from interactions between the circadian clock and sleep homeostat. Studies in humans have also suggested association between chronotypes and differential sleep homeostasis. Using EEGs, Taillard et al., 2003 reported that evening types are associated with slower increase in sleep pressure during extended wakefulness [25], whereas others using similar methods reported that morning types show higher homeostatic response to sleep disruption [90,91]. Thus, while differences in sleep homeostasis are associated with chronotypes, direction and nature of differences observed for human chronotypes vary across studies. Our findings extend this association to an evolutionary context, suggesting that chronotypes, which are typically thought of as a trait of clock alone are not only correlated with differential sleep homeostasis, but that the two co-evolve. The co-evolution of these traits raises the possibility of shared genetic mechanisms underlying their regulation.

Beyond responses to sleep loss, chronotype may also influence how sleep is regulated under circadian misalignment [27]. While social jetlag is a major confounding factor in humans, the same does not exist in our animal models. Our simulated jetlag using light alone as a cue showed that sleep responses varied across chronotypes when subjected to advance phase shift. Our *early* and *late* chronotypes exhibited opposing changes in sleep duration during day and night, respectively. Following phase advances, *early* flies exhibited a greater magnitude of change in daytime sleep duration than the other chronotypes. This parallels findings in humans, where morning chronotypes showed greater changes in wake time following eastward (advance time zone) travel [54]. Although the sleep parameters measured is different, the greater response of one chronotype to circadian misalignment is conceptually consistent across systems. In contrast, *late* flies exhibited a reduction in night sleep following phase advance. The advanced night window corresponds to the evening activity period under the original LD cycle, suggesting that *late* flies may retain some influence of the previous light regime. Additionally, night sleep latency after advance shift and day sleep fragmentation following either shift does not recover to baseline even till last day of recording (Day 3). A similar dissociation has been reported in humans, where sleep duration recovers rapidly following jetlag due to homeostatic mechanisms, while sleep architecture and timing take longer to readjust [54]. Together, these findings highlight the influence of chronotype on sleep responses to circadian misalignment and allow us to extend our inferences beyond this model system.

### Genetic basis of sleep differences across fly chronotype populations

A previous whole-genome characterisation of these chronotype populations identified several sleep-related genes harbouring SNPs, particularly in the *early* populations, including *kirre, Egfr, Fbxl-4, Tβh* and *Nlg4* [44]*. Egfr* is a particularly interesting candidate given its established role in sleep regulation [92]. Downregulation of *Egfr* or downstream molecules in *Egfr* signalling pathways have been reported to cause nighttime sleep fragmentation [93], resembling the phenotype that we observed in our *early* populations. It should also be noted that selection acted directly on timing of adult emergence, not on sleep, so it is possible that sleep phenotypes may arise due to pleiotropic effect of genes under direct selection or due to selective sweep of sleep specific genes. The molecular mechanisms behind observed sleep differences in fly chronotype populations therefore remains an open question. As most of the SNPs were found in regulatory region and had only modifier impact [44], expression level analysis of these candidate genes across chronotype populations needs to be examined, and transcriptome-wide profiling in these populations is a focus of ongoing work.

## Conclusion

In summary, our study demonstrates that artificial selection for divergent chronotype of eclosion rhythm in *Drosophila melanogaster* led to correlated evolution of sleep architecture, even though sleep was never the direct target of selection. The parallels between human chronotypes-associated differences in sleep timing, architecture, homeostatic regulation and phase advanced jetlag response, suggest some conserved features across systems. Our findings in this study, using a model system with common ancestry, population-level replicates and controlled environmental conditions, provides strong evidence for clear relationships between chronotypes and sleep organization. These findings present novel phenotypic characterizations that offer a basis for future mechanistic investigation. Moreover, these selection lines offer an evolutionary perspective on how multiple traits can coevolve in response to selection on timing of a behaviour, as expected in natural populations.

## Acknowledgements

We thank Abhilash Lakshman, Arijit Ghosh, Rutvij Kulkarni, Pragya Sharma and Meet Joshi for valuable discussions and suggestions during the course of this work. We thank Bhavi Kesharwani for helping with making LED panels for light intensity experiments and Asima Mishra for helping with some sleep deprivation experiments. Claude (Anthropic) was used to assist with drafting some portions of the manuscript and for writing R codes for analysis. We also thank Samuel, Muniraju and Rajanna for their technical assistance. We would like to acknowledge financial support through intramural funding from the Jawaharlal Nehru Centre for Advanced Scientific Research (JNCASR) and the Council of Scientific & Industrial Research (CSIR), New Delhi, India, for a junior research fellowship and a senior research fellowship to MR.

## Data availability

All data are available upon request.

## Supplementary figure legends

**Figure S1. Selection of *early* and *late* eclosion chronotypes.** (A) Schematic of artificial selection for *early* and *late* chronotypes from ancestral *control* populations; subscripts 1–4 denote the four replicate populations of each. Grey shaded region represents Lights OFF. Flies emerging in morning window (ZT21 - ZT01, blue rectangle) were collected to establish *early* populations and those emerging in evening window (ZT09 – ZT13, orange rectangle) to establish *late* populations. (B) Percentage emergence of flies in morning (left) and evening (right) windows across generations of selection; Error bars are ±SEM across 4 replicate populations. *early* flies emerge more in morning window and less in evening window whereas *late* flies emerge less in morning and more in evening window across generations.

ALT TEXT: Schematic representation of artificial selection for *early* and *late* eclosion chronotypes of *Drosophila melanogaster* from ancestral *control* populations and percentage emergence of flies of each chronotype in morning and evening windows across 412 generations of selection, where *early* flies emerge increasingly in the morning and *late* flies in the evening.

**Figure S2. Sleep and activity measures in fly chronotype populations.** (A) Daytime sleep parameters of virgin male (top) and female (bottom) chronotype flies under LD 12:12-latency, number of bouts and median bout duration. (B) Raster plots of sleep during one representative night (ZT12-24) in male (left) and female (right) chronotype flies from one replicate population (Block 1); each row represents one fly and each blue bar a sleep bout. (C) Activity profiles (left) and total activity counts during daytime and nighttime (right) of male (top) and female (bottom) flies. Profiles are averaged across 4 replicate populations, error bands are ±SEM across 4 populations and grey shaded regions represent lights OFF. Bar plots show means ± SEM across 4 populations, with symbols representing individual blocks. A two-way randomized block ANOVA (selection as a fixed factor and block as a random factor) was used for statistical comparisons; no significant differences among selection lines were found for any parameter (all *p* > 0.05; Table S2 and S4).

ALT TEXT: Graphs showing sleep and activity measures in virgin male and female chronotype flies. Panel A shows bar plots of daytime sleep latency, number of bouts and median bout duration, and panel C shows activity profiles and day and night activity counts, none of which differ among chronotypes. Panel B shows raster plots of individual sleep bouts during one representative night, where *early* flies have more fragmented sleep than *control*.

**Figure S3. Nighttime sleep-state architecture in fly chronotype populations.** (A) Number of nighttime bouts of each sleep state – short, intermediate and long – in virgin male (left) and female (right) chronotype flies under LD12:12. Male: short (F_(2,6)_ = 4.57, *p* = 0.062), intermediate (F_(2,6)_ = 5.54, *p* = 0.043), long (F_(2,6)_ = 0.98, *p* = 0.426); female: short (F_(2,6)_ = 4.16, *p* = 0.073), intermediate (F_(2,6)_ = 20.54, *p* = 0.002), long (F_(2,6)_ = 4.86, *p* = 0.056). Bar plots show means ± SEM across 4 replicate populations, with symbols representing individual blocks. A two-way randomized block ANOVA (selection as a fixed factor and block as a random factor) followed by Tukey’s HSD post hoc test was performed separately for each sleep state. Different letters on top of bar plots represent statistically significant differences (*p* < 0.05); no letters are shown where the ANOVA was not significant. (B) Probability of the first sleep bout of the night being short, intermediate or long in male (left) and female (right) flies; error bars are ±SEM of each state’s probability across 4 replicate populations. Data in B are descriptive and were not tested statistically.

ALT TEXT: Graphs showing the architecture of nighttime sleep states in virgin male and female chronotype flies. Panel A shows bar plots of the number of short, intermediate and long sleep bouts at night, where *early* flies have more intermediate bouts than *control* flies in both sexes. Panel B shows stacked bars of the probability that the first sleep bout of the night is short, intermediate or long, with *early* flies tending to begin the night with short sleep more often.

**Figure S4. Effect of light intensity modulation on sleep of chronotype populations.** Sleep parameters of male chronotype flies under different light intensities - 1 lux, 10 lux, 100 lux and 1000 lux. Top row – daytime and bottom row – nighttime sleep duration, latency, number of bouts and median bout duration. Points are means ±SEM across 4 populations. A randomised block ANOVA (selection and light intensity as fixed factors and block as random factor) followed by Tukey’s HSD post hoc test was used for statistical comparisons. Different letters above the points represent significant differences among light intensities (effect of light intensity, *p <* 0.05); the selection × light intensity interaction was not significant for any parameter. Full statistical results are given in Table S6.

ALT TEXT: Graphs showing day and night sleep parameters of virgin male chronotype flies under different light intensities-1, 10, 100 and 1000 lux. Daytime sleep and night sleep latency increase with light intensity. The response to light intensity is similar across chronotypes.

**Figure S5. Sleep in male chronotype flies across 12h thermophase and cryophase under DDTC** (A) Sleep parameters of virgin male chronotype flies under DDTC (temperature cycles in constant darkness; 28 °C thermophase, 18 °C cryophase) - sleep duration during thermophase (*F_(2,6)_* = 3.36, *p* = 0.105) and cryophase (*F_(2,6)_* = 0.33, *p* = 0.733); latency during thermophase (*F_(2,6)_* = 0.13, *p* = 0.877) and cryophase (*F_(2,6)_* = 5.83, *p* = 0.039), number of bouts during thermophase (*F_(2,6)_* = 2.17, *p* = 0.195) and cryophase (*F_(2,6)_* = 0.99, *p* = 0.422) and median bout duration during thermophase (*F_(2,6)_* = 1.89, *p* = 0.23) and cryophase (*F_(2,6)_* = 0.10, *p* = 0.903).

(B) Short, intermediate and long sleep profiles averaged across 4 replicate populations and 3 days under DDTC regime and midpoint of rise in long sleep (*F_(2,6)_* = 5.1, *p* = 0.051). Error bands in sleep profiles are ±SEM across 4 populations. Grey rectangle on top of sleep profiles represents lights off. Red and blue rectangles and shaded regions in sleep profiles represent thermo- and cryo-phase respectively. All bar plots show means ± SEM across 4 populations, with symbols representing individual blocks. A two-way randomized block ANOVA (selection as a fixed factor and block as a random factor) followed by Tukey’s HSD post hoc test was used for statistical comparisons. Different letters on top of bar plots represent statistically significant differences (*p* < 0.05); no letters are shown where the ANOVA was not significant.

ALT TEXT: Graphs showing sleep in male chronotype flies under DDTC cycles. Panel A shows bar plots of sleep duration, latency, number of bouts and median bout duration during thermophase and cryophase, where *late* flies take longer to fall asleep in cryophase than *early* flies. Panel B shows sleep profiles of each sleep state (short, intermediate and long) across temperature cycle and a bar plot for midpoint of rise in long sleep, which occurs earliest in *early* and latest in *late* chronotype flies.

**Figure S6. Change in sleep bout number and duration of chronotype populations after jetlag** (A) Change from baseline in daytime (top) and nighttime (bottom) number of bouts (left) and median sleep bout duration (right) of male chronotype flies after a 9h phase advance of the LD cycle. (B) Same as A for the 9h phase delay. Days −3 to −1 are baseline cycles and Days 1–3 are cycles after the shift; Day 0 (dashed line), the cycle during which the shift occurred, was not analysed. Points are means ± SEM across 4 replicate populations. A linear mixed-effects model with selection as a fixed factor, day as a repeated-measures factor and population (block) as a random factor, followed by Tukey’s HSD post hoc test, was used for statistical comparisons. Asterisks indicate significant differences between days (effect of day); the selection × day interaction was not significant for any parameter. \**p* < 0.05, \*\**p* < 0.01, \*\*\**p* < 0.001. Full model results are given in Table S11.

ALT TEXT: Graphs showing change from baseline in daytime and nighttime sleep bout number and median bout duration of male chronotype flies after a 9h phase advance (Panel A) and delay (Panel B). Bout number and bout duration change over the days after both shifts, but the change is similar across chronotypes.

**Figure S7. Sleep-state specific rebound and sleep architecture after night sleep deprivation in chronotype populations** (A) Long sleep profiles of male *early* (top), *control* (middle) and *late* (bottom) flies across 3 days of the sleep deprivation regime described in Figure 5; dashed lines represent undisturbed flies, and solid lines represent sleep-deprived flies. Time 0 represents lights ON on Day 1. Profiles are averaged across 4 replicate populations, error bands are ±SEM across 4 populations and grey shaded regions represent lights OFF. (B) Intermediate sleep rebound during the night window (12–24 h post deprivation) (F_(2,6)_ = 9.06, *p* = 0.015). (C) Number of bouts (selection: F(2,6) = 4.80, p = 0.057; day: F(1,9) = 13.29, p = 0.005; selection × day: F(2,9) = 14.32, p = 0.002) and median bout duration (selection: F(2,15) = 13.14, p < 0.001; day: F(1,15) = 0.26, p = 0.616; selection × day: F(2,15) = 0.66, p = 0.530) of sleep-deprived flies during the night before (Day 1) and after (Day 3) deprivation for standard sleep. Bar plots show means ± SEM across 4 populations, with symbols representing individual blocks. A two-way randomized block ANOVA (selection as a fixed factor and block as a random factor) was used for statistical comparisons in B and a linear mixed-effects model (selection as a fixed factor, day as a repeated-measures factor and block as a random factor) was used in C, followed by Tukey’s HSD post hoc test. Different letters on top of bar plots represent statistically significant differences (*p* < 0.05); no letters are shown where the ANOVA was not significant.

ALT TEXT: Graphs showing sleep rebound of male chronotype flies to 12h night sleep deprivation. Panel A shows long sleep profile of sleep deprived and undisturbed flies across 3-days. Panel B shows that *early* flies have less intermediate sleep rebound at night than other chronotypes. Panel C shows the number and median duration of standard sleep bouts at night before and after deprivation.

## Supplementary Tables Legends

In all statistical test tables, statistically significant terms (*p* <0.05) are highlighted in red. K, N in Tukey’s table represents number of means compared and number of replicates respectively.

**Table S1. Generations of selection:** Number of generations of selection of the *early, control* and *late* populations at the time of each experiment.

**Table S2. Sleep under LD12:12:** Two-way randomized block ANOVA (selection as a fixed factor and block as a random factor) for daytime and nighttime sleep parameters of virgin male (A) and female (B) chronotype flies under LD12:12. (C) Tukey’s HSD post hoc comparisons for nighttime sleep parameters with a significant effect of selection.

**Table S3. Effect sizes of selection on sleep:** Effect sizes (Cohen’s *d* ± 95% CI) for daytime and nighttime sleep parameters of *early* vs *control* (A, C) and *late* vs *control* (B, D) flies, for males (A, B) and females (C, D), shown for each experiment and replicate population (block).

**Table S4. Locomotor activity under LD12:12:** Two-way randomized block ANOVA for daytime and nighttime activity counts of virgin male (A) and female (B) chronotype flies under LD12:12.

**Table S5. Nighttime sleep states:** Two-way randomized block ANOVA for duration, number of bouts and latency of short (A), intermediate (int) (B) and long (C) sleep at night in virgin male and female chronotype flies under LD12:12, with Tukey’s HSD post hoc comparisons for parameters with a significant effect of selection.

**Table S6. Daytime sleep states:** Two-way randomized block ANOVA for duration, number of bouts and latency of short (A), intermediate (int) (B) and long (C) sleep at day in virgin male and female chronotype flies under LD12:12

**Table S7. Sleep across light intensities:** Randomized block ANOVA (selection and light intensity as fixed factors and block as a random factor) for daytime (A) and nighttime (B) sleep parameters of virgin male chronotype flies under 1, 10, 100 and 1000 lux. (C) Tukey’s HSD post hoc comparisons for the main effect of light intensity.

**Table S8. Sleep under DDTC:** Two-way randomized block ANOVA for sleep parameters of virgin male chronotype flies under DDTC: midpoint of sleep rise and behavioural night parameters (A), sleep parameters during the 12 h thermophase (B) and cryophase (C), and midpoint of rise in long sleep (D). (E) Tukey’s HSD post hoc comparisons for parameters with a significant effect of selection.

**Table S9. Midpoint of rise in sleep under LD12:12:** Midpoint of rise in sleep of virgin male chronotype flies under LD12:12. (A) Mean ± SEM midpoint (Zeitgeber time, h) and number of flies for each replicate population (block). (B) Two-way randomized block ANOVA (selection as a fixed factor and block as a random factor).

**Table S10. Phase advance:** Linear mixed-effects model (selection as a fixed factor, day as a repeated-measures factor and block as a random factor) for change from baseline in sleep parameters of male chronotype flies after a 9 h phase advance. (A) Type III ANOVA with Satterthwaite df. (B) Tukey’s HSD post hoc comparisons for significant terms.

**Table S11. Phase delay:** Same as Table S10 for the 9 h phase delay.

**Table S12. Sleep rebound after sleep deprivation:** Two-way randomized block ANOVA for day (0–12 h), night (12–24 h) and 24 h sleep rebound after 12 h of night sleep deprivation in male chronotype flies, using standard (std) (A), long (B), intermediate (int) (C) and short (D) sleep definitions, with Tukey’s HSD post hoc comparisons for night rebound of long and intermediate sleep.

**Table S13. Sleep fragmentation before and after sleep deprivation:** Linear mixed-effects model (selection as a fixed factor, day as a repeated-measures factor and block as a random factor) for nighttime number of bouts and median bout duration of sleep-deprived male flies on Day 1 (pre-deprivation) and Day 3 (post-deprivation). (A) Type III ANOVA with Satterthwaite df. (B) Tukey’s HSD post hoc comparisons for significant terms.

## References

1. Sharma VK. Adaptive Significance of Circadian Clocks. Chronobiology International. 2003;20(6):901–919. doi:10.1081/CBI-120026099

2. Roenneberg T. What is chronotype?: Preface. Sleep and Biological Rhythms. 2012;10(2):75–76. doi:10.1111/j.1479-8425.2012.00541.x

3. Rittenhouse JL, Robart AR, Watts HE. Variation in chronotype is associated with migratory timing in a songbird. Biol Lett. 2019;15(8):20190453. doi:10.1098/rsbl.2019.0453

4. Labyak SE, Lee TM, Goel N. Rhythm chronotypes in a diurnal rodent, Octodon degus. American Journal of Physiology-Regulatory, Integrative and Comparative Physiology. 1997;273(3):R1058–R1066. doi:10.1152/ajpregu.1997.273.3.R1058

5. Martorell-Barceló M, Abecasis D, Akaarir M, et al. Revealing Chronotypes Across Aquatic Species Using Acoustic Telemetry. doi:10.1111/faf.70022

6. Roenneberg T, Wirz-Justice A, Merrow M. Life between Clocks: Daily Temporal Patterns of Human Chronotypes. J Biol Rhythms. 2003;18(1):80–90. doi:10.1177/0748730402239679

7. Li R, Rodríguez-Muñoz R, Tregenza T. Evidence for chronotypes and associations with fitness-related traits in a wild insect.

8. Horne JA, Ostberg O. A self-assessment questionnaire to determine morningness-eveningness in human circadian rhythms. Int J Chronobiol. 1976;4(2):97–110.

9. Duffy JF, Dijk DJ, Hall EF, Czeisler CA. Relationship of endogenous circadian melatonin and temperature rhythms to self-reported preference for morning or evening activity in young and older people. J Investig Med. 1999;47(3):141–150.

10. Pagani L, Semenova EA, Moriggi E, et al. The Physiological Period Length of the Human Circadian Clock In Vivo Is Directly Proportional to Period in Human Fibroblasts. PLOS ONE. 2010;5(10):e13376. doi:10.1371/journal.pone.0013376

11. Hida A, Kitamura S, Ohsawa Y, et al. In vitro circadian period is associated with circadian/sleep preference. Scientific Reports. 2013;3(1):2074. doi:10.1038/srep02074

12. Ferrante A, Gellerman D, Ay A, et al. Diurnal Preference Predicts Phase Differences in Expression of Human Peripheral Circadian Clock Genes. J Circadian Rhythms. 13:4. doi:10.5334/jcr.ae

13. Chan ZM, Patel PP, Levitsky GM, McClung CA, Nouraie SM, Kelly NJ. Heart rate phase is an indicator of chronotype-relevant circadian shifts associated with human disease: an *All of Us* Research Program analysis. doi:10.1113/JP290337

14. Chauhan S, Norbury R, Faßbender KC, Ettinger U, Kumari V. Beyond sleep: A multidimensional model of chronotype. Neuroscience & Biobehavioral Reviews. 2023;148:105114. doi:10.1016/j.neubiorev.2023.105114

15. Merikanto I, Lahti T, Puolijoki H, et al. Associations of Chronotype and Sleep With Cardiovascular Diseases and Type 2 Diabetes. Chronobiology International. 2013;30(4):470–477. doi:10.3109/07420528.2012.741171

16. Mongrain V, Carrier J, Dumont M. Chronotype and Sex Effects on Sleep Architecture and Quantitative Sleep EEG in Healthy Young Adults. Sleep. 2005;28(7):819–827. doi:10.1093/sleep/28.7.819

17. McMahon DM, Burch JB, Youngstedt SD, et al. Relationships between chronotype, social jetlag, sleep, obesity and blood pressure in healthy young adults. Chronobiology International. 2019;36(4):493–509. doi:10.1080/07420528.2018.1563094

18. Zhai. Full article: Associations of Sleep Insufficiency and Chronotype with Inflammatory Cytokines in College Students. 2022. Accessed August 14, 2024. https://www.tandfonline.com/doi/full/10.2147/NSS.S329894

19. Suh S, Yang HC, Kim N, et al. Chronotype Differences in Health Behaviors and Health-Related Quality of Life: A Population-Based Study Among Aged and Older Adults. Behavioral Sleep Medicine. 2017;15(5):361–376. doi:10.1080/15402002.2016.1141768

20. Partonen T. Chronotype and Health Outcomes: An Update. Curr Sleep Medicine Rep. 2026;12(1):10. doi:10.1007/s40675-026-00366-y

21. Gulec M, Selvi Y, Boysan M, Aydin A, Oral E, Aydin EF. Chronotype effects on general well-being and psychopathology levels in healthy young adults. Biological Rhythm Research. 2013;44(3):457–468. doi:10.1080/09291016.2012.704795

22. Yuri G, Chamorro R, Tobar N, Cifuentes M. The human chronotype: A multidimensional construct at the crossroad of physiology and behavior, with important health implications. Sleep Medicine Reviews. 2025;83:102142. doi:10.1016/j.smrv.2025.102142

23. Kerkhof GA, Lancel M. EEG Slow Wave Activity, REM Sleep, and Rectal Temperature During Night and Day Sleep in Morning-Type and Evening-Type Subjects. Psychophysiology. 1991;28(6):678–688. doi:10.1111/j.1469-8986.1991.tb01014.x

24. Colelli DR, Dela Cruz GR, Kendzerska T, Murray BJ, Boulos MI. Impact of sleep chronotype on in-laboratory polysomnography parameters. Journal of Sleep Research. 2023;32(5):e13922. doi:10.1111/jsr.13922

25. Taillard J, Philip P, Coste O, Sagaspe P, Bioulac B. The circadian and homeostatic modulation of sleep pressure during wakefulness differs between morning and evening chronotypes. Journal of Sleep Research. 2003;12(4):275–282. doi:10.1046/j.0962-1105.2003.00369.x

26. Putilov AA, Donskaya OG. What Can Make the Difference Between Chronotypes in Sleep Duration? Testing the Similarity of Their Homeostatic Processes. Front Neurosci. 2022;16. doi:10.3389/fnins.2022.832807

27. Roenneberg T, Pilz LK, Zerbini G, Winnebeck EC. Chronotype and Social Jetlag: A (Self-) Critical Review. Biology. 2019;8(3):54. doi:10.3390/biology8030054

28. Levandovski R, Sasso E, Hidalgo MP. Chronotype: a review of the advances, limits and applicability of the main instruments used in the literature to assess human phenotype. Trends Psychiatry Psychother. 2013;35:3–11. 10.1590/S2237-60892013000100002

29. Juda M, Vetter C, Roenneberg T. Chronotype Modulates Sleep Duration, Sleep Quality, and Social Jet Lag in Shift-Workers. J Biol Rhythms. 2013;28(2):141–151. doi:10.1177/0748730412475042

30. Aili K, Åström-Paulsson S, Stoetzer U, Svartengren M, Hillert L. Reliability of Actigraphy and Subjective Sleep Measurements in Adults: The Design of Sleep Assessments. J Clin Sleep Med. 2017;13(1):39–47. doi:10.5664/jcsm.6384

31. Allebrandt KV, Teder-Laving M, Kantermann T, et al. Chronotype and sleep duration: the influence of season of assessment. Chronobiol Int. 2014;31(5):731–740. doi:10.3109/07420528.2014.901347

32. Roepke SE, Duffy JF. Differential impact of chronotype on weekday and weekend sleep timing and duration. Nat Sci Sleep. 2010;2010(2):213–220. doi:10.2147/nss.s12572

33. Wright KP, McHill AW, Birks BR, Griffin BR, Rusterholz T, Chinoy ED. Entrainment of the human circadian clock to the natural light-dark cycle. Curr Biol. 2013;23(16):1554–1558. doi:10.1016/j.cub.2013.06.039

34. Shang Y, Griffith LC, Rosbash M. Light-arousal and circadian photoreception circuits intersect at the large PDF cells of the Drosophila brain. Proceedings of the National Academy of Sciences. 2008;105(50):19587–19594. doi:10.1073/pnas.0809577105

35. Cajochen C, Stefani O, Schöllhorn I, Lang D, Chellappa S. Influence of evening light exposure on polysomnographically assessed night-time sleep: A systematic review with meta-analysis. Lighting Research & Technology. 2022;54(6):609–624. doi:10.1177/14771535221078765

36. Mazzotta G, Damulewicz M, Cusumano P. Better Sleep at Night: How Light Influences Sleep in Drosophila. Frontiers in Physiology. 2020;11:997. doi:10.3389/fphys.2020.00997

37. Wams EJ, Woelders T, Marring I, et al. Linking Light Exposure and Subsequent Sleep: A Field Polysomnography Study in Humans. Sleep. 2017;40(12). doi:10.1093/sleep/zsx165

38. Porcheret K, Wald L, Fritschi L, et al. Chronotype and environmental light exposure in a student population. Chronobiology International. 2018;35(10):1365–1374. doi:10.1080/07420528.2018.1482556

39. van der Zwet G, Bor Z, Bos R, et al. Sex and age modulate the relationship between melanopsin-dependent light sensitivity and chronotype. Sleep. 2026;49(6):zsag057. doi:10.1093/sleep/zsag057

40. Shawa N, Rae DE, Roden LC. Impact of seasons on an individual’s chronotype: current perspectives. Nature and Science of Sleep. 2018;10:345–354. doi:10.2147/NSS.S158596

41. Kumar S, Kumar D, Paranjpe DA, R AC, Sharma VK. Selection on the timing of adult emergence results in altered circadian clocks in fruit flies Drosophila melanogaster. J Exp Biol. 2007;210(Pt 5):906–918. doi:10.1242/jeb.001354

42. Nikhil KL, Abhilash L, Sharma VK. Molecular Correlates of Circadian Clocks in Fruit Fly *Drosophila melanogaster* Populations Exhibiting *early* and *late* Emergence Chronotypes. J Biol Rhythms. 2016;31(2):125–141. doi:10.1177/0748730415627933

43. Abhilash L, Sharma VK. On the relevance of using laboratory selection to study the adaptive value of circadian clocks. Physiological Entomology. 2016;41(4):293–306. doi:10.1111/phen.12158

44. Ghosh A, Abhilash L, Sharma VK, Nikhil KL, Sheeba V. Pooled sequencing reveals patterns of underlying genomic divergence in Drosophila chronotypes. bioRxiv. Preprint posted online November 16, 2026:2026.11.14.683300. doi:10.64898/2026.11.14.683300

45. Jones SE, Lane JM, Wood AR, et al. Genome-wide association analyses of chronotype in 697,828 individuals provides insights into circadian rhythms. Nat Commun. 2019;10(1):343. doi:10.1038/s41467-018-08259-7

46. Shaw PJ, Cirelli C, Greenspan RJ, Tononi G. Correlates of sleep and waking in Drosophila melanogaster. Science. 2000;287(5459):1834–1837. doi:10.1126/science.287.5459.1834

47. Hendricks JC, Finn SM, Panckeri KA, et al. Rest in Drosophila is a sleep-like state. Neuron. 2000;25(1):129–138. doi:10.1016/s0896-6273(00)80877-6

48. Abhilash L, Evans R, Shafer OT. Recognition of distinct sleep states in Drosophila uncovers previously obscured homeostatic and circadian control of sleep. Current Biology. 2026;36(4):968–978.e4. doi:10.1016/j.cub.2026.01.015

49. Ghosh A, Sharma P, Dansana S, Sheeba V. Evidence for Co-Evolution of Masking With Circadian Phase in Drosophila Melanogaster. J Biol Rhythms. 2021;36(3):254–270. doi:10.1177/0748730421997262

50. Vaze KM, KL N, L A, Sharma VK. Early- and Late-Emerging Drosophila melanogaster Fruit Flies Differ in Their Sensitivity to Light During Morning and Evening. Chronobiology International. 2012;29(6):674–682. doi:10.3109/07420528.2012.680557

51. Bonduriansky R, Day T. Nongenetic Inheritance and Its Evolutionary Implications. Annual Review of Ecology, Evolution, and Systematics. 2009;40(1):103–125. doi:10.1146/annurev.ecolsys.39.110707.173441

52. Persons JL, Abhilash L, Lopatkin AJ, et al. PHASE: An Open-Source Program for the Analysis of DrosophilaPhase, Activity, and Sleep Under Entrainment. J Biol Rhythms. 2022;37(4):455–467. doi:10.1177/07487304221093114

53. Savitzky Abraham, Golay MJE. Smoothing and Differentiation of Data by Simplified Least Squares Procedures. Anal Chem. 1964;36(8):1627–1639. doi:10.1021/ac60214a047

54. Willoughby AR, Vallat R, Ong JL, Chee MWL. Insights about travel-related sleep disruption from 1.5 million nights of data. Sleep. 2025;48(7):zsaf077. doi:10.1093/sleep/zsaf077

55. Borenstein M, Hedges LV, Higgins JPT, Rothstein HR. Generality of the Basic Inverse-Variance Method. In: Introduction to Meta-Analysis. 2009:311–319. doi:10.1002/9780470743386.ch34

56. Faville R, Kottler B, Goodhill GJ, Shaw PJ, van Swinderen B. How deeply does your mutant sleep? Probing arousal to better understand sleep defects in Drosophila. Sci Rep. 2015;5:8454. doi:10.1038/srep08454

57. Huber R, Hill SL, Holladay C, Biesiadecki M, Tononi G, Cirelli C. Sleep homeostasis in Drosophila melanogaster. Sleep. 2004;27(4):628–639. doi:10.1093/sleep/27.4.628

58. Ghosh A, Harbison S. 0038 Inferring the Genetic Basis of Sleep States in Drosophila with Hidden Markov Models (HMMs). Sleep. 2026;49(Supplement_1):Page A17. 10.1093/sleep/zsag091.0038

59. Wiggin TD, Goodwin PR, Donelson NC, et al. Covert sleep-related biological processes are revealed by probabilistic analysis in Drosophila. Proceedings of the National Academy of Sciences. 2020;117(18):10024–10034. doi:10.1073/pnas.1917573117

60. Alphen B van, Yap MHW, Kirszenblat L, Kottler B, Swinderen B van. A Dynamic Deep Sleep Stage in Drosophila. J Neurosci. 2013;33(16):6917–6927. doi:10.1523/JNEUROSCI.0061-13.2013

61. Stahl BA, Slocumb ME, Chaitin H, DiAngelo JR, Keene AC. Sleep-Dependent Modulation of Metabolic Rate in Drosophila. Sleep. 2017;40(8):zsx084. doi:10.1093/sleep/zsx084

62. Tainton-Heap LAL, Kirszenblat LC, Notaras ET, et al. A Paradoxical Kind of Sleep in Drosophila melanogaster. Current Biology. 2021;31(3):578–590.e6. doi:10.1016/j.cub.2020.10.081

63. Schlichting M, Menegazzi P, Rosbash M, Helfrich-Förster C. A Distinct Visual Pathway Mediates High-Intensity Light Adaptation of the Circadian Clock in Drosophila. J Neurosci. 2019;39(9):1621–1630. doi:10.1523/JNEUROSCI.1497-18.2018

64. Parisky KM, Agosto Rivera JL, Donelson NC, Kotecha S, Griffith LC. Reorganization of Sleep by Temperature in Drosophila Requires Light, the Homeostat, and the Circadian Clock. Curr Biol. 2016;26(7):882–892. doi:10.1016/j.cub.2016.02.011

65. Yadlapalli S, Jiang C, Bahle A, Reddy P, Meyhofer E, Shafer OT. Circadian clock neurons constantly monitor environmental temperature to set sleep timing. Nature. 2018;555(7694):98–102. doi:10.1038/nature25740

66. Ma H, Liu G, Zhang S, et al. Effects of thermal environment on sleep quality and comfort: Neurophysiological evidence from sleep microstructure. Building and Environment. 2026;296:114502. doi:10.1016/j.buildenv.2026.114502

67. Nikhil KL, Goirik G, Ratna K, Sharma VK. Role of Temperature in Mediating Morning and Evening Emergence Chronotypes in Fruit Flies Drosophila melanogaster. J Biol Rhythms. 2014;29(6):427–441. doi:10.1177/0748730414553797

68. Vaze KM, Kannan NN, Abhilash L, Sharma VK. Chronotype differences in Drosophila are enhanced by semi-natural conditions. Naturwissenschaften. 2012;99(11):967–971. doi:10.1007/s00114-012-0978-1

69. Abhilash L, Kalliyil A, Sheeba V. Responses of activity rhythms to temperature cues evolve in Drosophila populations selected for divergent timing of eclosion. J Exp Biol. 2020;223(Pt 11):jeb222414. doi:10.1242/jeb.222414

70. Abhilash L, Ghosh A, Sheeba V. Selection for Timing of Eclosion Results in Co-evolution of Temperature Responsiveness in Drosophila melanogaster. J Biol Rhythms. 2019;34(6):596–609. doi:10.1177/0748730419877315

71. Avidan A, Colwell C. Jet lag syndrome: circadian organization, pathophysiology, and management strategies. NSS. Published online August 2010:187. doi:10.2147/NSS.S6683

72. Borbély AA. A two process model of sleep regulation. Human neurobiology. 1982;1 3:195–204.

73. Borbély A. The two-process model of sleep regulation: Beginnings and outlook. Journal of Sleep Research. 2022;31(4):e13598. doi:10.1111/jsr.13598

74. Chowdhury B, Abhilash L, Ortega A, Liu S, Shafer O. Homeostatic control of deep sleep and molecular correlates of sleep pressure in Drosophila. Elife. 2023;12:e91355. doi:10.7554/eLife.91355

75. Vyazovskiy VV, Achermann P, Tobler I. Sleep homeostasis in the rat in the light and dark period. Brain Research Bulletin. 2007;74(1):37–44. doi:10.1016/j.brainresbull.2007.05.001

76. Zhang N, Zhu Y, Cao B. Effects of environmental temperature mimicking circadian rhythms on thermal comfort and sleep quality. Building and Environment. 2025;281:113196. doi:10.1016/j.buildenv.2025.113196

77. Kondo M, Tokura H, Wakamura T, et al. Physiological Significance of Cyclic Changes in Room Temperature around Dusk and Dawn for Circadian Rhythms of Core and Skin Temperature, Urinary 6-hydroxymelatonin Sulfate, and Waking Sensation just after Rising. Journal of PHYSIOLOGICAL ANTHROPOLOGY. 2007;26(4):429–436. doi:10.2114/jpa2.26.429

78. Abhilash L, Shafer OT. A two-process model of Drosophila sleep reveals an inter-dependence between circadian clock speed and the rate of sleep pressure decay. Sleep. 2023;47(2):zsad277. doi:10.1093/sleep/zsad277

79. Fuller PM, Gooley JJ, Saper CB. Neurobiology of the Sleep-Wake Cycle: Sleep Architecture, Circadian Regulation, and Regulatory Feedback. J Biol Rhythms. 2006;21(6):482–493. doi:10.1177/0748730406294627

80. Tabuchi M. Dynamic neuronal instability generates synaptic plasticity and behavior: Insights from *Drosophila* sleep. Neuroscience Research. 2024;198:1–7. doi:10.1016/j.neures.2023.06.009

81. Liu S, Lamaze A, Liu Q, et al. WIDE AWAKE Mediates the Circadian Timing of Sleep Onset. Neuron. 2014;82(1):151–166. doi:10.1016/j.neuron.2014.01.040

82. Tabuchi M, Monaco JD, Duan G, et al. Clock-Generated Temporal Codes Determine Synaptic Plasticity to Control Sleep. Cell. 2018;175(5):1213–1227.e18. doi:10.1016/j.cell.2018.09.016

83. Merikanto I, Kuula L, Makkonen T, et al. Morningness is associated with lower sleep spindle amplitudes and intensities during adolescence. Sleep Medicine. 2017;40:e221. doi:10.1016/j.sleep.2017.11.645

84. Nikhil KL, Ratna K, Sharma VK. Life-history traits of Drosophila melanogaster populations exhibiting early and late eclosion chronotypes. BMC Evol Biol. 2016;16:46. doi:10.1186/s12862-016-0622-3

85. Fropf R, Zhou H, Yin JCP. The clock gene *period* differentially regulates sleep and memory in *Drosophila*. Neurobiology of Learning and Memory. 2018;153:2–12. doi:10.1016/j.nlm.2018.02.016

86. Horn M, Mitesser O, Hovestadt T, Yoshii T, Rieger D, Helfrich-Förster C. The Circadian Clock Improves Fitness in the Fruit Fly, Drosophila melanogaster. Front Physiol. 2019;10. doi:10.3389/fphys.2019.01374

87. Chung BY, Kilman VL, Keath JR, Pitman JL, Allada R. The GABAA Receptor RDL Acts in Peptidergic PDF Neurons to Promote Sleep in Drosophila. Current Biology. 2009;19(5):386–390. doi:10.1016/j.cub.2009.01.040

88. Parisky KM, Agosto J, Pulver SR, et al. PDF Cells Are a GABA-Responsive Wake-Promoting Component of the Drosophila Sleep Circuit. Neuron. 2008;60(4):672–682. doi:10.1016/j.neuron.2008.10.042

89. Raman S, Coogan AN. A Cross-Sectional Study of the Associations between Chronotype, Social Jetlag and Subjective Sleep Quality in Healthy Adults. Clocks & Sleep. 2020;2(1):1–6. doi:10.3390/clockssleep2010001

90. Mongrain V, Dumont M. Increased Homeostatic Response to Behavioral Sleep Fragmentation in Morning Types Compared to Evening Types. Sleep. 2007;30(6):773–780. doi:10.1093/sleep/30.6.773

91. Mongrain V, Carrier J, Dumont M. Circadian and homeostatic sleep regulation in morningness–eveningness. Journal of Sleep Research. 2006;15(2):162–166. doi:10.1111/j.1365-2869.2006.00532.x

92. Harbison ST, Negron YLS, Hansen NF, Lobell AS. Selection for long and short sleep duration in Drosophila melanogaster reveals the complex genetic network underlying natural variation in sleep. PLOS Genetics. 2017;13(12):e1007098. doi:10.1371/journal.pgen.1007098

93. Foltenyi K, Greenspan RJ, Newport JW. Activation of EGFR and ERK by rhomboid signaling regulates the consolidation and maintenance of sleep in Drosophila. Nat Neurosci. 2007;10(9):1160–1167. doi:10.1038/nn1957

