## Supplementary for "Evidence for correlated evolution of sleep architecture in chronotype populations of *Drosophila melanogaster* created by artificial selection"

#### Figure S1

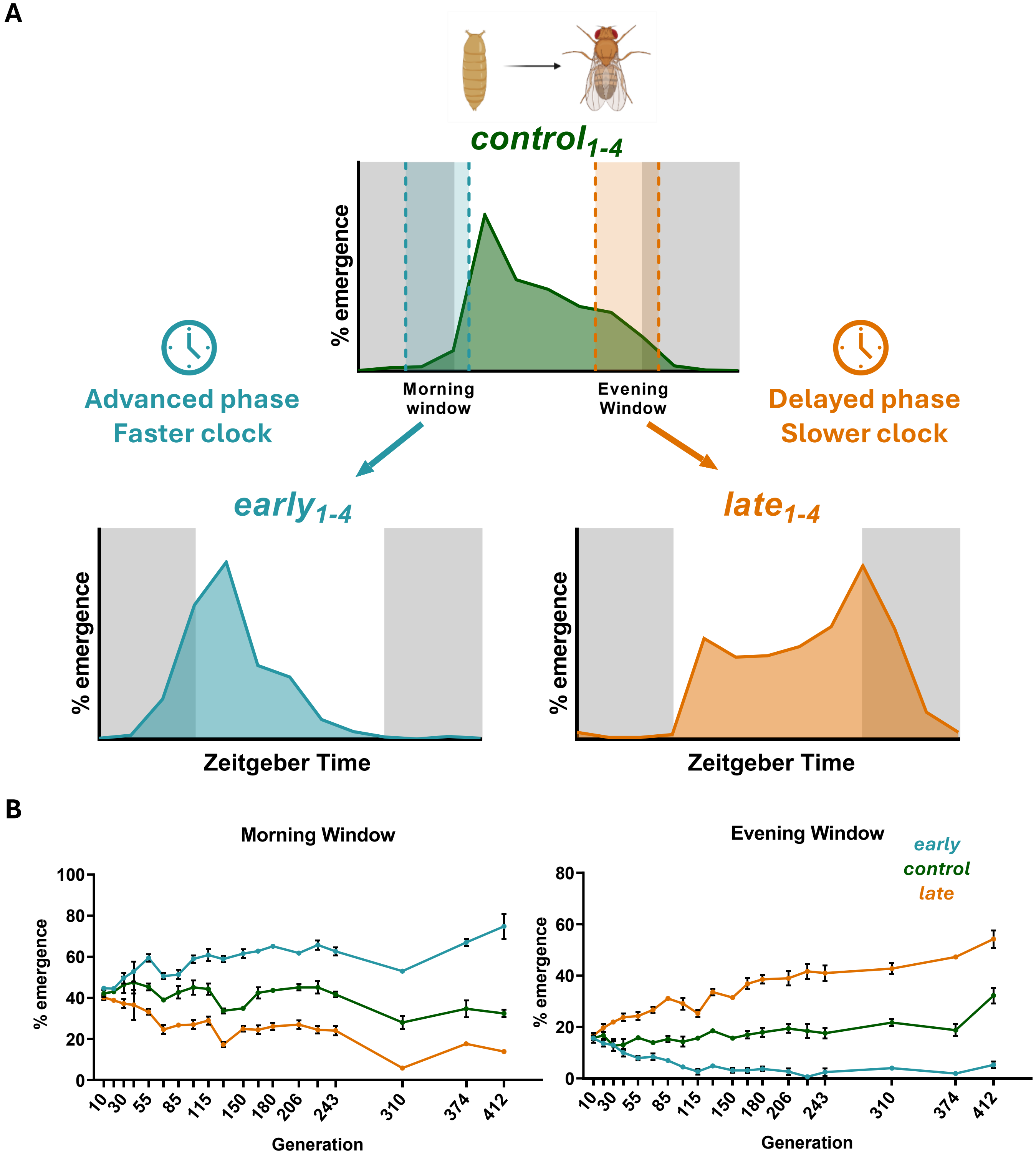
A
control1-4
% emergence
Morning window
Evening Window
Advanced phase
Faster clock
Delayed phase
Slower clock
early1-4
late1-4
% emergence
Zeitgeber Time
% emergence
Zeitgeber Time
B
Morning Window
Evening Window
early
control
late
% emergence
100
80
60
40
20
0
10
30
55
85
115
150
180
206
243
310
374
412
Generation
% emergence
80
60
40
20
0
Generation

### Figure S2

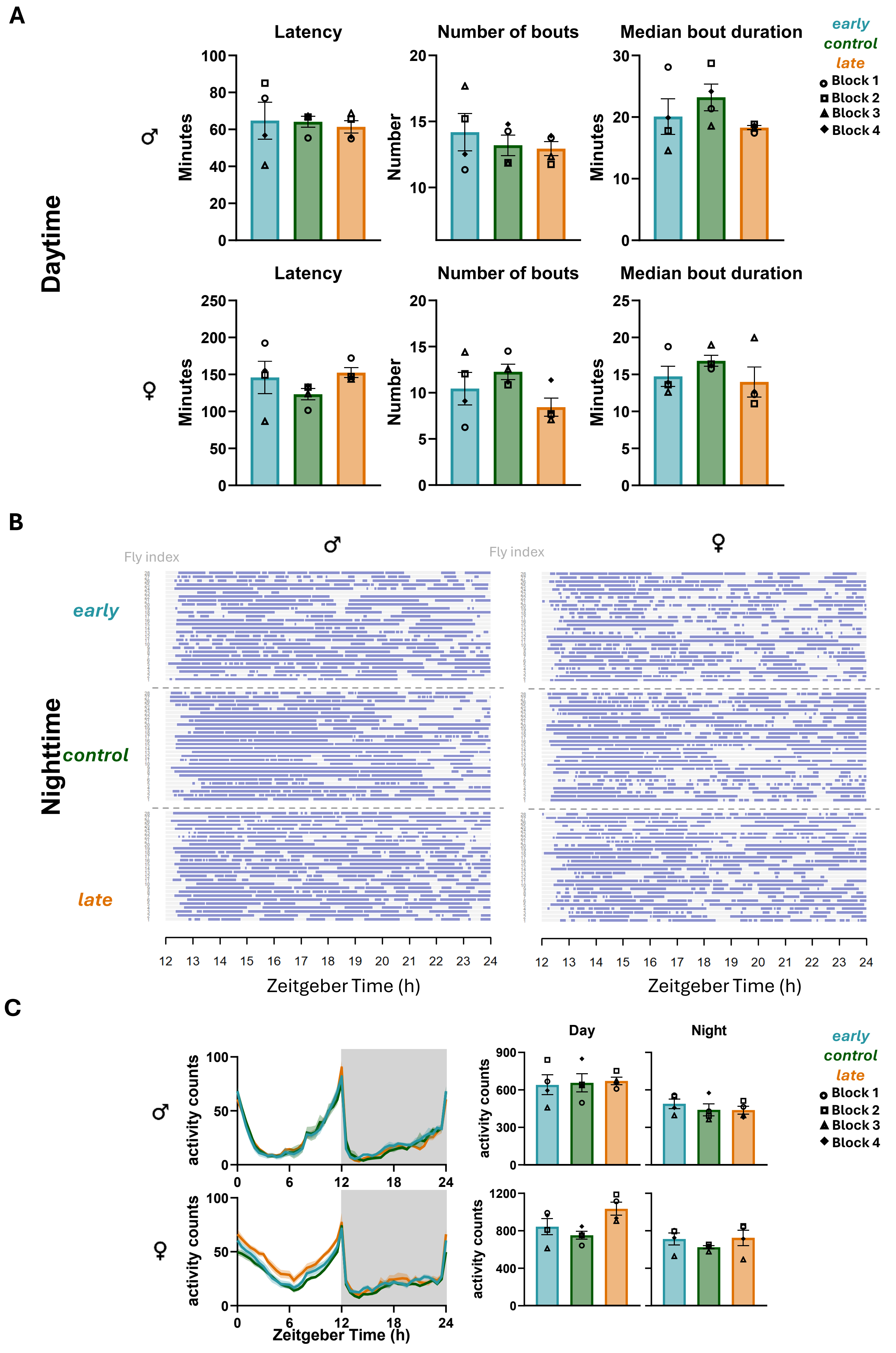
A
Daytime
Latency
Number of bouts
Median bout duration
Minutes
Number
early
control
late
Block 1
Block 2
Block 3
Block 4
B
Nighttime
Fly index
early
control
late
Zeitgeber Time (h)
C
Day
Night
activity counts
Zeitgeber Time (h)

### Figure S3

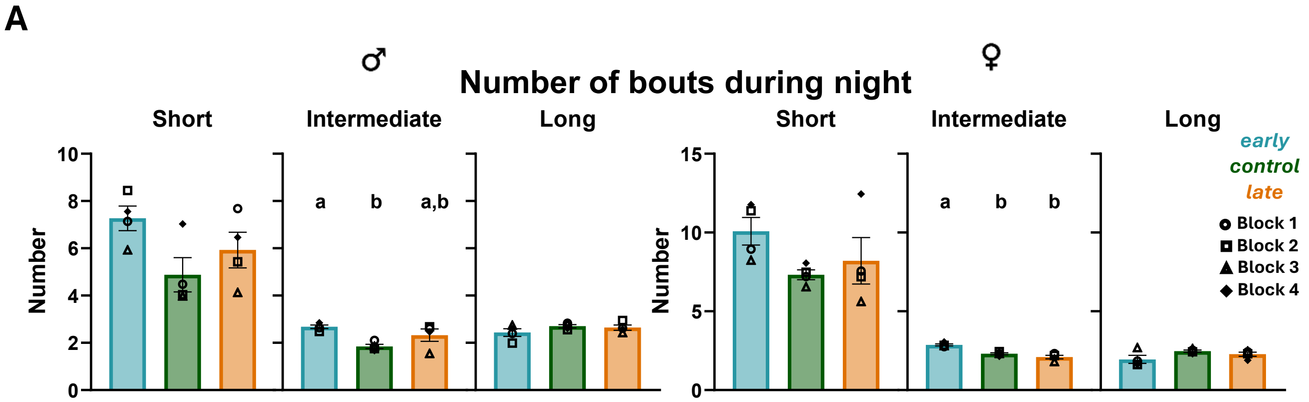
A
♂
Number of bouts during night
♀
Short
Intermediate
Long
Short
Intermediate
Long
Number
a
b
a,b
a
b
b
early
control
late
Block 1
Block 2
Block 3
Block 4

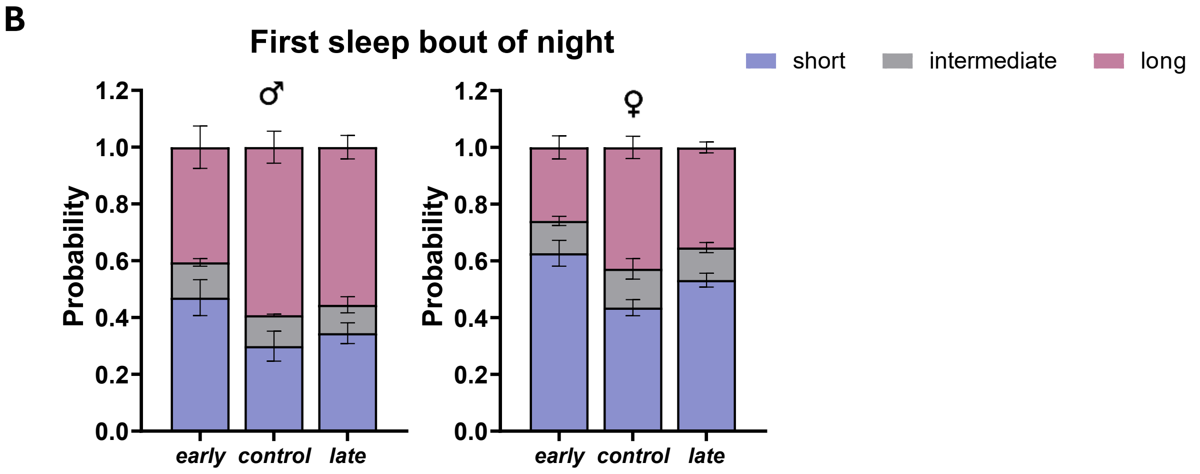
B
First sleep bout of night
short
intermediate
long
♂
♀
Probability
early
control
late

### Figure S4

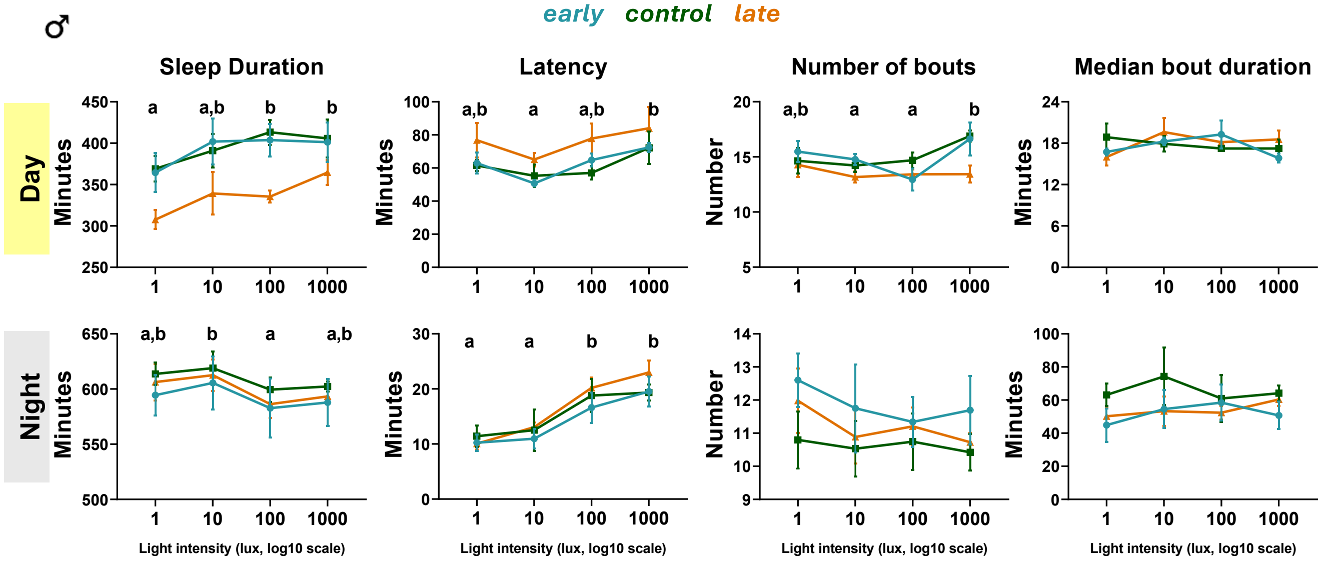
♂
early control late
Sleep Duration
Latency
Number of bouts
Median bout duration
Day
Night
Minutes
Number
a
a,b
b
Light intensity (lux, log10 scale)
1
10
100
1000

### Figure S5

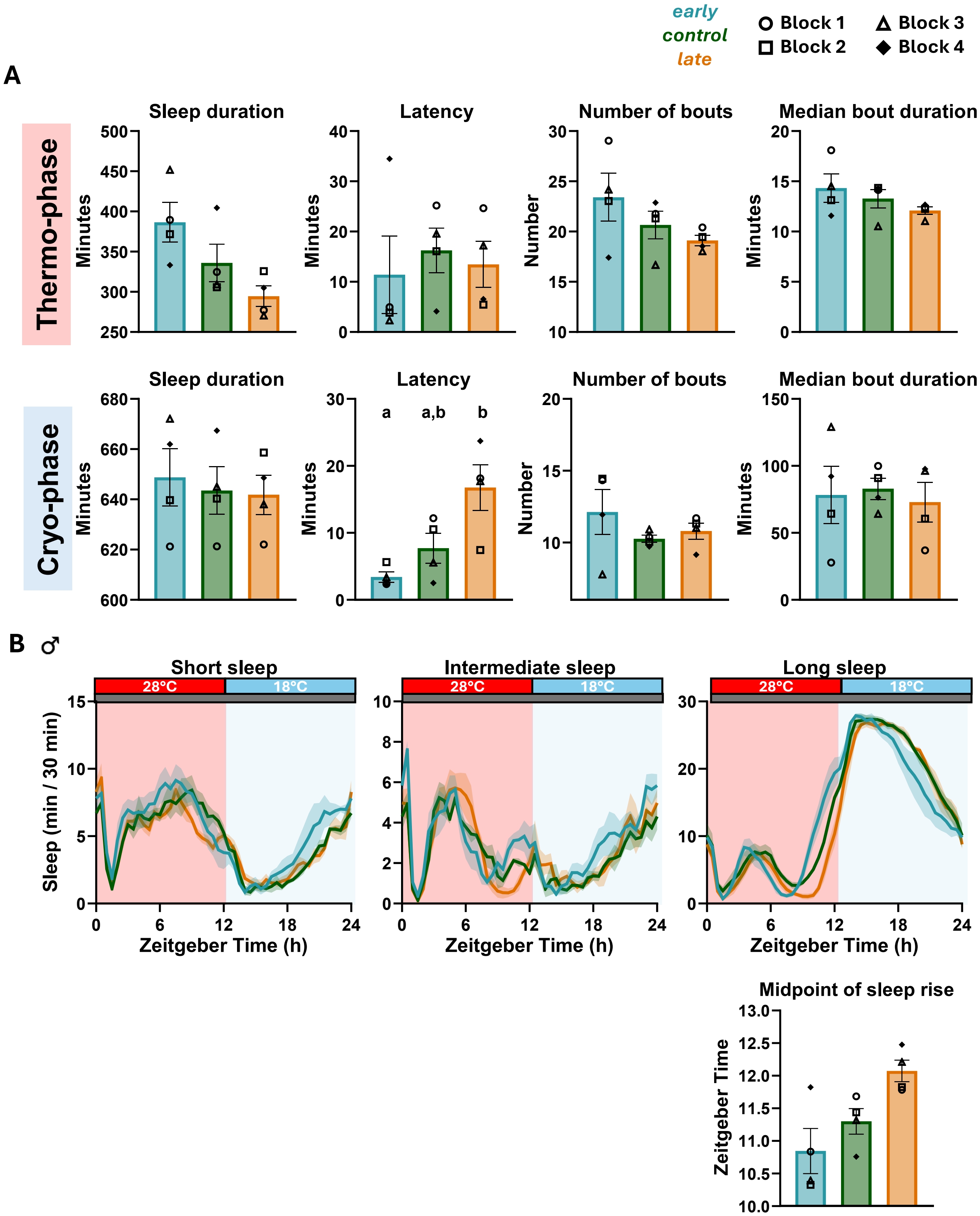
early
control
late
Block 1
Block 2
Block 3
Block 4
A
Thermo-phase
Sleep duration
Minutes
Latency
Minutes
Number of bouts
Number
Median bout duration
Minutes
Cryo-phase
Sleep duration
Minutes
Latency
a
a,b
b
Minutes
Number of bouts
Number
Median bout duration
Minutes
B ♂
Short sleep
Intermediate sleep
Long sleep
28°C
18°C
Sleep (min / 30 min)
Zeitgeber Time (h)
Midpoint of sleep rise
Zeitgeber Time

### Figure S6

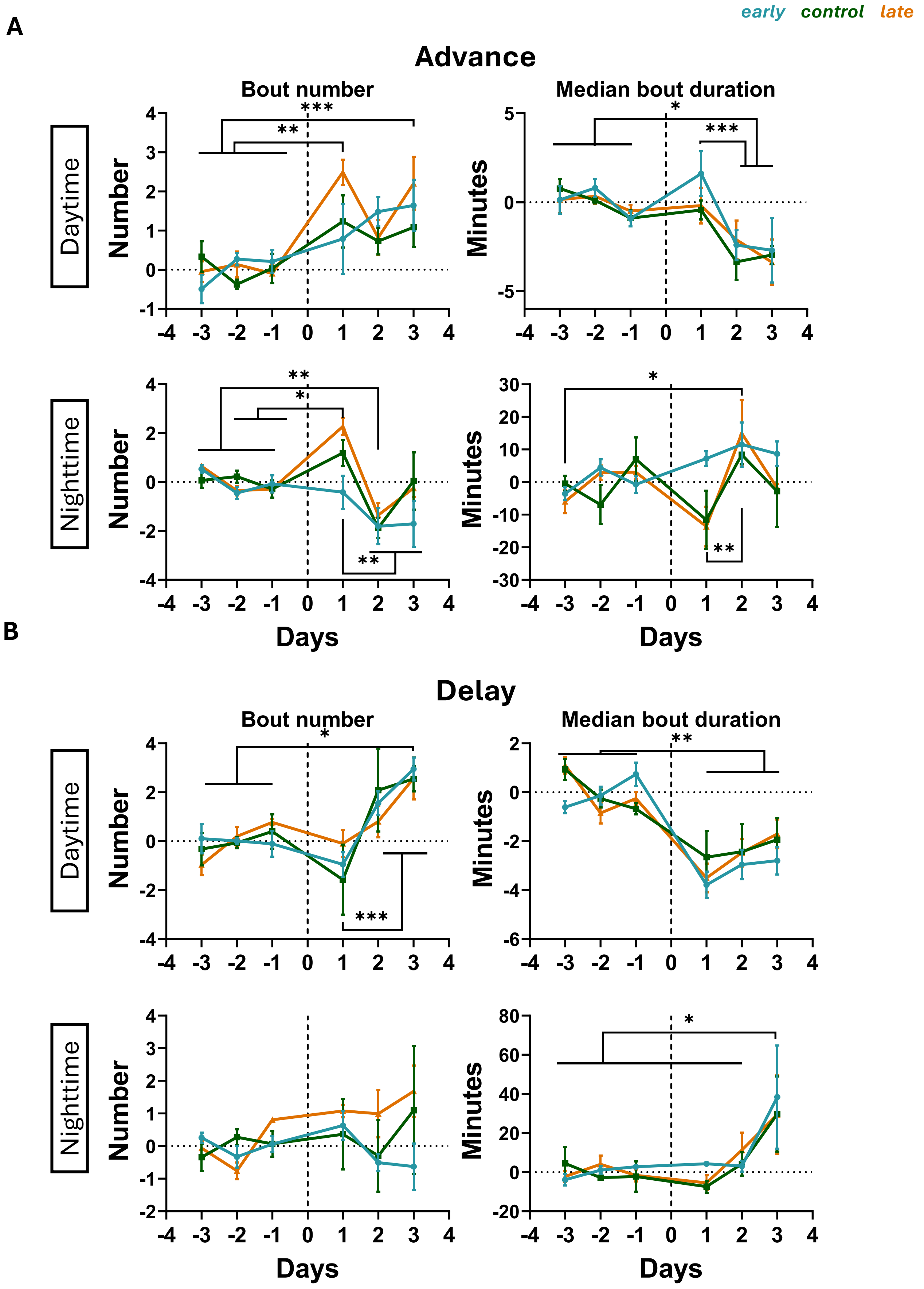

early control late
A
Advance
Bout number
Median bout duration
Daytime
Nighttime
Number
Minutes
Days
B
Delay
Bout number
Median bout duration
Daytime
Nighttime
Number
Minutes
Days

### Figure S7

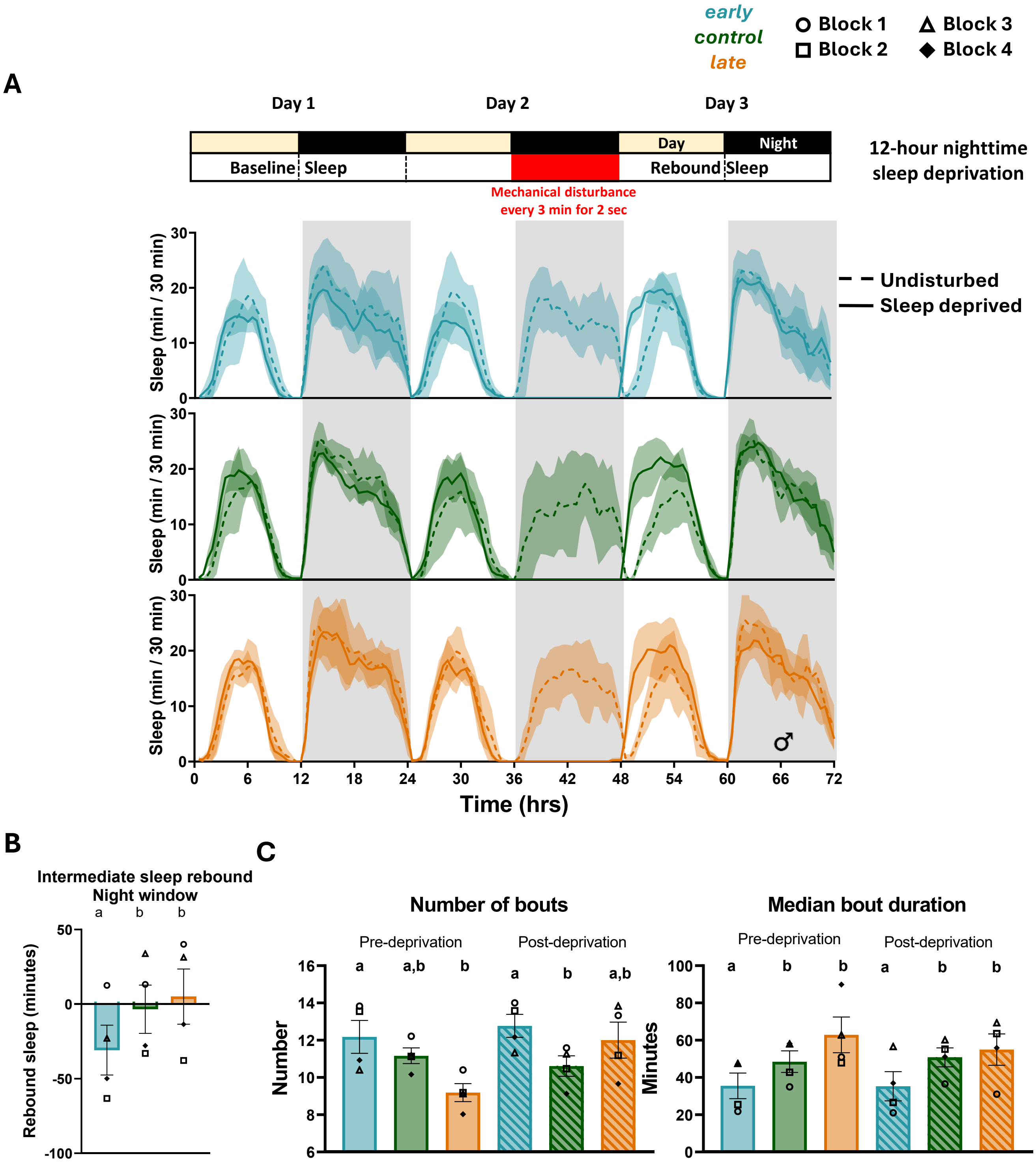
early
control
late
Block 1
Block 2
Block 3
Block 4
A
Day 1
Day 2
Day 3
Day
Night
Baseline
Sleep
Rebound
Sleep
12-hour nighttime sleep deprivation
Mechanical disturbance every 3 min for 2 sec
Undisturbed
Sleep deprived
Sleep (min / 30 min)
Time (hrs)
0
6
12
18
24
30
36
42
48
54
60
66
72
B
Intermediate sleep rebound
Night window
a
b
b
Rebound sleep (minutes)
C
Number of bouts
Median bout duration
Pre-deprivation
Post-deprivation
a
a,b
b
a
b
a,b
Number
a
b
b
a
b
b
Minutes

**Table S1. Generations of selection:** Number of generations of selection of the *early*, *control* and *late* populations at the time of each. experiment

| Experiment | Number of generations of selection |
|---|---|
| LD 12:12 Exp. 1 ♂ ♀ | ~363 |
| LD 12:12 Exp. 2 ♂ ♀ | ~364 |
| LD 12:12 Exp. 3 ♂ ♀ | ~364 |
| LD 12:12 Exp. 4 ♂ | ~365 |
| LD 12:12 Exp. 5 ♂ | ~365 |
| Light intensity Block1 | 390 |
| Light intensity Block2 | 392 |
| Light intensity Block3 | 392 |
| Light intensity Block4 | 392 |
| DDTC Exp. 1 | 410 |
| DDTC Exp. 2 | 420 |
| DDTC Exp. 3 | 420 |
| Jetlag | 415 |
| Sleep deprivation Block1 | 390 |
| Sleep deprivation Block2 | 380 |
| Sleep deprivation Block3 | 390 |
| Sleep deprivation Block4 | 380 |

**Table S2. Sleep under LD12:12:** Two-way randomized block ANOVA (selection as a fixed factor and block as a random factor) for daytime and nighttime sleep parameters of virgin male (A) and female (B) chronotype flies under LD12:12. (C) Tukey's HSD post hoc comparisons for nighttime sleep parameters with a significant effect of selection.

A

| **Day sleep duration ♂** | **Effect** | **SS** | **$DF_n$** | **MS** | **$DF_d$** | **MS error** | **F** | **p** |
|---|---|---|---|---|---|---|---|---|
| selection | Fixed | 1477 | 2 | 738 | 6 | 1370.827 | 0.539 | 0.609 |
| block | Random | 1632 | 3 | 544 | 6 | 1370.827 | 0.397 | 0.760 |
| selection*block | Random | 8225 | 6 | 1371 | 0 | 0 | | |
| Error | | | 0 | | | | | |
| **Day latency ♂** | **Effect** | **SS** | **$DF_n$** | **MS** | **$DF_d$** | **MS error** | **F** | **p** |
| selection | Fixed | 26.71 | 2 | 13.36 | 6 | 183.5746 | 0.073 | 0.931 |
| block | Random | 342.27 | 3 | 114.09 | 6 | 183.5746 | 0.622 | 0.626 |
| selection*block | Random | 1101.45 | 6 | 183.57 | 0 | 0 | | |
| Error | | | 0 | | | | | |
| **Day bout numbers ♂** | **Effect** | **SS** | **$DF_n$** | **MS** | **$DF_d$** | **MS error** | **F** | **p** |
| selection | Fixed | 3.457 | 2 | 1.729 | 6 | 5.439866 | 0.318 | 0.739 |
| block | Random | 2.147 | 3 | 0.716 | 6 | 5.439866 | 0.132 | 0.938 |
| selection*block | Random | 32.639 | 6 | 5.44 | 0 | 0 | | |
| Error | | | 0 | | | | | |
| **Day med. Bout duration ♂** | **Effect** | **SS** | **$DF_n$** | **MS** | **$DF_d$** | **MS error** | **F** | **p** |
| selection | Fixed | 49.524 | 2 | 24.762 | 6 | 18.02719 | 1.374 | 0.323 |
| block | Random | 49.465 | 3 | 16.488 | 6 | 18.02719 | 0.915 | 0.488 |
| selection*block | Random | 108.163 | 6 | 18.027 | 0 | 0 | | |
| Error | | | 0 | | | | | |

| **Night sleep duration ♂** | **Effect** | **SS** | **$DF_n$** | **MS** | **$DF_d$** | **MS error** | **F** | **p** |
|---|---|---|---|---|---|---|---|---|
| selection | Fixed | 2072 | 2 | 1036 | 6 | 507.987 | 2.039 | 0.211 |
| block | Random | 4675 | 3 | 1558 | 6 | 507.987 | 3.068 | 0.113 |
| selection*block | Random | 3048 | 6 | 508 | 0 | 0 | | |
| Error | | | 0 | | | | | |
| **Night latency ♂** | **Effect** | **SS** | **$DF_n$** | **MS** | **$DF_d$** | **MS error** | **F** | **p** |
| selection | Fixed | 42.055 | 2 | 21.027 | 6 | 2.61324 | 8.046 | 0.020 |
| block | Random | 82.619 | 3 | 27.54 | 6 | 2.61324 | 10.539 | 0.008 |
| selection*block | Random | 15.679 | 6 | 2.613 | 0 | 0 | | |
| Error | | | 0 | | | | | |

| Night bout numbers ♂ | Effect | SS | $DF_n$ | MS | $DF_d$ | MS error | F | p |
|---|---|---|---|---|---|---|---|---|
| selection | Fixed | 18.039 | 2 | 9.02 | 6 | 1.339166 | 6.735 | 0.029 |
| block | Random | 11.256 | 3 | 3.752 | 6 | 1.339166 | 2.802 | 0.131 |
| selection*block | Random | 8.035 | 6 | 1.339 | 0 | 0 | | |
| Error | | | 0 | | | | | |
| **Night med. bout duration ♂** | **Effect** | **SS** | **$DF_n$** | **MS** | **$DF_d$** | **MS error** | **F** | **p** |
| selection | Fixed | 954.84 | 2 | 477.42 | 6 | 118.271 | 4.037 | **0.077** |
| block | Random | 353.53 | 3 | 117.84 | 6 | 118.271 | 0.996 | 0.456 |
| selection*block | Random | 709.63 | 6 | 118.27 | 0 | 0 | | |
| Error | | | 0 | | | | | |

B

| Day sleep duration ♀ | Effect | SS | $DF_n$ | MS | $DF_d$ | MS error | F | p |
|---|---|---|---|---|---|---|---|---|
| selection | Fixed | 9543.1 | 2 | 4771.6 | 6 | 2415.5 | 1.975 | 0.219 |
| block | Random | 9289.2 | 3 | 3096.4 | 6 | 2415.5 | 1.282 | 0.363 |
| selection*block | Random | 14493 | 6 | 2415.5 | 0 | 0 | | |
| Error | | | 0 | | | | | |
| **Day latency ♀** | **Effect** | **SS** | **$DF_n$** | **MS** | **$DF_d$** | **MS error** | **F** | **p** |
| selection | Fixed | 1881.7 | 2 | 940.8 | 6 | 786.1873 | 1.197 | 0.365 |
| block | Random | 2257.8 | 3 | 752.6 | 6 | 786.1873 | 0.957 | 0.471 |
| selection*block | Random | 4717.1 | 6 | 786.2 | 0 | 0 | | |
| Error | | | 0 | | | | | |
| **Day bout numbers ♀** | **Effect** | **SS** | **$DF_n$** | **MS** | **$DF_d$** | **MS error** | **F** | **p** |
| selection | Fixed | 29.487 | 2 | 14.743 | 6 | 8.674181 | 1.700 | 0.260 |
| block | Random | 5.632 | 3 | 1.877 | 6 | 8.674181 | 0.216 | 0.882 |
| selection*block | Random | 52.045 | 6 | 8.674 | 0 | 0 | | |
| Error | | | 0 | | | | | |
| **Day med. Bout duration ♀** | **Effect** | **SS** | **$DF_n$** | **MS** | **$DF_d$** | **MS error** | **F** | **p** |
| selection | Fixed | 17.382 | 2 | 8.691 | 6 | 9.345373 | 0.930 | 0.445 |
| block | Random | 21.872 | 3 | 7.291 | 6 | 9.345373 | 0.780 | 0.547 |
| selection*block | Random | 56.072 | 6 | 9.345 | 0 | 0 | | |
| Error | | | 0 | | | | | |

| Night sleep duration ♀ | Effect | SS | $DF_n$ | MS | $DF_d$ | MS error | F | p |
|---|---|---|---|---|---|---|---|---|
| selection | Fixed | 3308 | 2 | 1654 | 5 | 230.536 | 7.174 | 0.034 |
| block | Random | 4959 | 3 | 1653 | 5 | 230.536 | 7.171 | 0.029 |
| selection*block | Random | 1153 | 5 | 231 | 0 | 0 | | |
| Error | | | 0 | | | | | |
| **Night latency ♀** | **Effect** | **SS** | **$DF_n$** | **MS** | **$DF_d$** | **MS error** | **F** | **p** |
| selection | Fixed | 450.7 | 2 | 225.35 | 6 | 44.08687 | 5.112 | **0.051** |
| block | Random | 62.42 | 3 | 20.81 | 6 | 44.08687 | 0.472 | 0.713 |
| selection*block | Random | 264.52 | 6 | 44.09 | 0 | 0 | | |
| Error | | | 0 | | | | | |
| **Night bout numbers ♀** | **Effect** | **SS** | **$DF_n$** | **MS** | **$DF_d$** | **MS error** | **F** | **p** |
| selection | Fixed | 18.341 | 2 | 9.17 | 6 | 2415.5 | 4.917 | **0.054** |
| block | Random | 18.495 | 3 | 6.165 | 6 | 2415.5 | 3.306 | 0.099 |
| selection*block | Random | 11.19 | 6 | 1.865 | 0 | 0 | | |
| Error | | | 0 | | | | | |
| **Night med. bout duration ♀** | **Effect** | **SS** | **$DF_n$** | **MS** | **$DF_d$** | **MS error** | **F** | **p** |
| selection | Fixed | 633.11 | 2 | 316.55 | 6 | 15.2367 | 20.776 | 0.002 |
| block | Random | 397.12 | 3 | 132.37 | 6 | 15.2367 | 8.688 | 0.013 |
| selection*block | Random | 91.42 | 6 | 15.24 | 0 | 0 | | |
| Error | | | 0 | | | | | |

C

| Tukey's HSD test- night sleep parameters | | | | |
|---|---|---|---|---|
| Mean difference | Latency ♂ | Bout numbers ♂ | Sleep duration ♀ | Median bout duration ♀ |
| ***early* vs *control*** | 2.6383 | -3.0029 | -40.5240 | -17.0016 |
| ***early* vs *late*** | 4.5672 | -1.5429 | -13.3529 | -13.0421 |
| ***control* vs *late*** | 1.9289 | 1.4600 | 27.1710 | 3.9596 |
| Minimum significant difference | 3.5071 | 2.5106 | 32.9404 | 8.4685 |
| q critical | 4.339 | 4.339 | 4.339 | 4.339 |
| K, N | 3, 4 | 3, 4 | 3, 4 | 3, 4 |

**Table S3. Effect sizes of selection on sleep:** Effect sizes (Cohen's d ± 95% CI) for daytime and nighttime sleep parameters of *early vs control* (A, C) and *late vs control* (B, D) flies, for males (A, B) and females (C, D), shown for each experiment and replicate population (block).

**A**

| ***Cohen's d ± 95% CI: early vs control ♂*** | | | | | | | |
|---|---|---|---|---|---|---|---|
| **Experiment** | **Block** | **n (E, C)** | **Day** | **Night** | | | |
| | | | **Sleep duration** | **Sleep duration** | **Latency** | **Bout numbers** | **Med. bout duration** |
| **Exp. 1** | 1 | 29, 30 | -0.43 ± 0.52 | -0.55 ± 0.52 | -0.26 ± 0.51 | 0.69 ± 0.53 | -0.47 ± 0.52 |
| | 2 | 28, 31 | -0.92 ± 0.54 | -0.89 ± 0.54 | -0.21 ± 0.51 | 1.16 ± 0.55 | -1.05 ± 0.55 |
| | 3 | 30, 30 | 0.72 ± 0.52 | -0.07 ± 0.51 | -0.25 ± 0.51 | 0.76 ± 0.52 | -0.71 ± 0.52 |
| | 4 | 31, 29 | 0.62 ± 0.52 | 0.10 ± 0.51 | 0.03 ± 0.51 | 0.32 ± 0.51 | -0.13 ± 0.51 |
| **Exp. 2** | 1 | 25, 27 | -1.18 ± 0.59 | -1.76 ± 0.64 | 0.69 ± 0.56 | 0.66 ± 0.56 | -0.75 ± 0.56 |
| | 2 | 28, 32 | 0.19 ± 0.51 | 0.10 ± 0.51 | -0.68 ± 0.52 | 0.92 ± 0.53 | -0.48 ± 0.51 |
| | 3 | 32, 31 | -0.39 ± 0.50 | 0.24 ± 0.50 | -0.03 ± 0.49 | -0.12 ± 0.49 | 0.14 ± 0.49 |
| | 4 | 30, 31 | 0.16 ± 0.50 | -0.39 ± 0.51 | 0.21 ± 0.50 | 0.87 ± 0.53 | -1.03 ± 0.53 |
| **Exp. 3** | 1 | 24, 31 | -0.45 ± 0.54 | -1.44 ± 0.60 | 0.63 ± 0.55 | 0.63 ± 0.55 | -0.73 ± 0.55 |
| | 2 | 29, 31 | -0.31 ± 0.51 | -0.25 ± 0.51 | -0.43 ± 0.51 | 0.61 ± 0.52 | -0.70 ± 0.52 |
| | 3 | 31, 28 | 0.63 ± 0.52 | 0.45 ± 0.52 | -0.40 ± 0.52 | -0.25 ± 0.51 | 0.42 ± 0.52 |
| | 4 | 31, 23 | -0.80 ± 0.56 | -0.94 ± 0.57 | 0.70 ± 0.56 | 1.18 ± 0.58 | -0.82 ± 0.56 |
| **Exp. 4** | 1 | 32, 30 | 0.15 ± 0.50 | -0.29 ± 0.50 | 0.18 ± 0.50 | 0.10 ± 0.50 | 0.05 ± 0.50 |
| | 2 | 29, 32 | -0.43 ± 0.51 | -0.26 ± 0.50 | -0.46 ± 0.51 | 0.81 ± 0.52 | -0.45 ± 0.51 |
| | 3 | 32, 32 | 0.81 ± 0.51 | 0.63 ± 0.50 | -0.28 ± 0.49 | -0.77 ± 0.51 | 0.72 ± 0.51 |
| | 4 | 31, 32 | -0.15 ± 0.49 | -0.06 ± 0.49 | 0.07 ± 0.49 | 0.25 ± 0.50 | -0.21 ± 0.50 |
| **Exp. 5** | 1 | 32, 30 | -1.18 ± 0.54 | -1.31 ± 0.55 | 0.48 ± 0.51 | 0.82 ± 0.52 | -0.59 ± 0.51 |
| | 2 | 28, 32 | -0.49 ± 0.51 | -0.15 ± 0.51 | -0.91 ± 0.53 | 0.97 ± 0.54 | -0.63 ± 0.52 |
| | 3 | 32, 30 | 0.47 ± 0.50 | 0.43 ± 0.50 | -0.29 ± 0.50 | 0.07 ± 0.50 | 0.17 ± 0.50 |
| | 4 | 31, 29 | -0.58 ± 0.52 | -0.66 ± 0.52 | -0.01 ± 0.51 | 0.91 ± 0.53 | -0.56 ± 0.52 |

**B**

| ***Cohen's d ± 95% CI: late vs control ♂*** | | | | | | | |
|---|---|---|---|---|---|---|---|
| **Experiment** | **Block** | **n (L, C)** | **Day** | **Night** | | | |
| | | | **Sleep duration** | **Sleep duration** | **Latency** | **Bout numbers** | **Med. bout duration** |
| **Exp. 1** | 1 | 30, 30 | -0.35 ± 0.51 | -0.16 ± 0.51 | 0.15 ± 0.51 | 0.71 ± 0.52 | -0.39 ± 0.51 |
| | 2 | 29, 31 | -0.74 ± 0.52 | -0.56 ± 0.52 | 0.20 ± 0.51 | 0.85 ± 0.53 | -0.72 ± 0.52 |
| | 3 | 30, 30 | -0.32 ± 0.51 | -0.18 ± 0.51 | 0.02 ± 0.51 | -0.14 ± 0.51 | 0.13 ± 0.51 |
| | 4 | 26, 29 | 0.19 ± 0.53 | 0.38 ± 0.53 | 0.07 ± 0.53 | 0.03 ± 0.53 | -0.13 ± 0.53 |
| **Exp. 2** | 1 | 30, 27 | -0.86 ± 0.54 | -0.85 ± 0.54 | 0.69 ± 0.53 | 0.60 ± 0.53 | -0.65 ± 0.53 |
| | 2 | 28, 32 | 0.10 ± 0.51 | -0.24 ± 0.51 | 0.13 ± 0.51 | 0.43 ± 0.51 | -0.58 ± 0.52 |
| | 3 | 32, 31 | -0.76 ± 0.51 | -0.14 ± 0.49 | 0.22 ± 0.50 | -0.12 ± 0.49 | 0.22 ± 0.50 |
| | 4 | 32, 31 | 0.18 ± 0.49 | 0.03 ± 0.49 | 0.47 ± 0.50 | 0.25 ± 0.50 | -0.29 ± 0.50 |
| **Exp. 3** | 1 | 32, 31 | 0.47 ± 0.50 | -0.23 ± 0.50 | 0.57 ± 0.50 | 0.05 ± 0.49 | -0.47 ± 0.50 |
| | 2 | 30, 31 | -0.84 ± 0.52 | -1.23 ± 0.55 | 0.23 ± 0.50 | 0.97 ± 0.53 | -0.96 ± 0.53 |
| | 3 | 32, 28 | -0.50 ± 0.51 | -0.03 ± 0.51 | 0.30 ± 0.51 | -0.28 ± 0.51 | 0.54 ± 0.52 |
| | 4 | 32, 23 | -0.82 ± 0.56 | -0.76 ± 0.55 | 0.98 ± 0.57 | 0.66 ± 0.55 | -0.11 ± 0.54 |
| **Exp. 4** | 1 | 32, 30 | -0.03 ± 0.50 | -0.37 ± 0.50 | 0.25 ± 0.50 | 0.48 ± 0.51 | -0.28 ± 0.50 |
| | 2 | 28, 32 | -0.48 ± 0.51 | -0.02 ± 0.51 | -0.22 ± 0.51 | 0.32 ± 0.51 | -0.36 ± 0.51 |
| | 3 | 32, 32 | -1.04 ± 0.52 | 0.23 ± 0.49 | -0.43 ± 0.50 | -0.91 ± 0.51 | 0.60 ± 0.50 |
| | 4 | 32, 32 | -0.43 ± 0.50 | 0.15 ± 0.49 | 0.26 ± 0.49 | -0.20 ± 0.49 | 0.17 ± 0.49 |

| | | | | | | | |
|---|---|---|---|---|---|---|---|
| Exp. 5 | 1 | 32, 30 | -0.71 ± 0.51 | -0.88 ± 0.52 | 0.66 ± 0.51 | 0.11 ± 0.50 | -0.04 ± 0.50 |
| | 2 | 31, 32 | 0.04 ± 0.49 | 1.25 ± 0.54 | -0.74 ± 0.51 | -0.18 ± 0.49 | 0.19 ± 0.50 |
| | 3 | 32, 30 | -0.88 ± 0.52 | 0.11 ± 0.50 | -0.38 ± 0.50 | -0.34 ± 0.50 | 0.64 ± 0.51 |
| | 4 | 31, 29 | -1.08 ± 0.54 | -0.69 ± 0.52 | 0.73 ± 0.52 | 0.06 ± 0.51 | -0.09 ± 0.51 |

**C**

| ***Cohen's d ± 95% CI: early vs control ♀*** | | | | | | | |
|---|---|---|---|---|---|---|---|
| **Experiment** | **Block** | **n (E, C)** | **Day** | **Night** | | | |
| | | | **Sleep duration** | **Sleep duration** | **Latency** | **Bout numbers** | **Med. bout duration** |
| **Exp. 1** | 1 | 29, 31 | -0.84 ± 0.53 | -0.53 ± 0.52 | 0.19 ± 0.51 | 0.32 ± 0.51 | -0.35 ± 0.51 |
| | 2 | 31, 32 | -0.13 ± 0.49 | -0.52 ± 0.50 | 0.07 ± 0.49 | 0.72 ± 0.51 | -0.63 ± 0.51 |
| | 3 | 32, 32 | 0.40 ± 0.49 | -0.13 ± 0.49 | -0.85 ± 0.51 | 0.42 ± 0.50 | -0.70 ± 0.50 |
| | 4 | 31, 32 | -0.06 ± 0.49 | -0.36 ± 0.50 | 0.22 ± 0.50 | 0.72 ± 0.51 | -0.45 ± 0.50 |
| **Exp. 2** | 1 | 32, 30 | -0.76 ± 0.52 | -0.60 ± 0.51 | 0.35 ± 0.50 | -0.46 ± 0.50 | 0.25 ± 0.50 |
| | 2 | 32, 27 | 0.08 ± 0.51 | 0.28 ± 0.51 | -0.55 ± 0.52 | 0.01 ± 0.51 | 0.22 ± 0.51 |
| | 3 | 29, 30 | 0.75 ± 0.53 | 0.03 ± 0.51 | -0.15 ± 0.51 | 0.65 ± 0.52 | -0.67 ± 0.52 |
| | 4 | 30, 30 | 0.30 ± 0.51 | -0.11 ± 0.51 | 0.02 ± 0.51 | 0.39 ± 0.51 | -0.14 ± 0.51 |
| **Exp. 3** | 1 | 31, 32 | -1.18 ± 0.54 | -0.90 ± 0.52 | 0.29 ± 0.50 | 0.12 ± 0.49 | 0.09 ± 0.49 |
| | 2 | 31, 30 | 0.09 ± 0.50 | -0.26 ± 0.50 | 0.09 ± 0.50 | 0.04 ± 0.50 | -0.61 ± 0.51 |
| | 3 | 31, 31 | 0.88 ± 0.52 | 1.02 ± 0.53 | -1.42 ± 0.56 | -0.58 ± 0.51 | 0.27 ± 0.50 |
| | 4 | 32, 31 | -0.35 ± 0.50 | -1.15 ± 0.53 | 0.04 ± 0.49 | 1.63 ± 0.57 | -0.91 ± 0.52 |

**D**

| ***Cohen's d ± 95% CI: late vs control ♀*** | | | | | | | |
|---|---|---|---|---|---|---|---|
| **Experiment** | **Block** | **n (L, C)** | **Day** | **Night** | | | |
| | | | **Sleep duration** | **Sleep duration** | **Latency** | **Bout numbers** | **Med. bout duration** |
| **Exp. 1** | 1 | 32, 31 | -0.89 ± 0.52 | -0.15 ± 0.49 | 0.51 ± 0.50 | 0.04 ± 0.49 | -0.10 ± 0.49 |
| | 2 | 31, 32 | -1.19 ± 0.54 | -0.17 ± 0.49 | 0.16 ± 0.49 | -0.13 ± 0.49 | -0.11 ± 0.49 |
| | 3 | 30, 32 | -0.67 ± 0.51 | -0.26 ± 0.50 | 0.30 ± 0.50 | -0.32 ± 0.50 | 0.03 ± 0.50 |
| | 4 | 32, 32 | 0.27 ± 0.49 | -0.28 ± 0.49 | 0.33 ± 0.49 | 0.62 ± 0.50 | -0.24 ± 0.49 |
| **Exp. 2** | 1 | 30, 30 | -1.00 ± 0.54 | -0.16 ± 0.51 | 0.35 ± 0.51 | -0.07 ± 0.51 | -0.13 ± 0.51 |
| | 2 | 31, 27 | -0.50 ± 0.52 | 0.53 ± 0.52 | -0.40 ± 0.52 | -0.46 ± 0.52 | 0.34 ± 0.52 |
| | 3 | 28, 30 | -0.50 ± 0.52 | -0.34 ± 0.52 | 0.26 ± 0.52 | 0.19 ± 0.52 | -0.53 ± 0.52 |
| | 4 | 31, 30 | 0.29 ± 0.50 | -0.21 ± 0.50 | 0.71 ± 0.52 | 0.87 ± 0.52 | -0.28 ± 0.50 |
| **Exp. 3** | 1 | 32, 32 | -1.11 ± 0.53 | -0.42 ± 0.50 | 0.12 ± 0.49 | -0.05 ± 0.49 | 0.22 ± 0.49 |
| | 2 | 32, 30 | 0.10 ± 0.50 | 0.38 ± 0.50 | -0.03 ± 0.50 | -0.82 ± 0.52 | 0.61 ± 0.51 |
| | 3 | 32, 31 | 0.77 ± 0.51 | 0.53 ± 0.50 | -0.47 ± 0.50 | -0.57 ± 0.50 | 0.32 ± 0.50 |
| | 4 | 32, 31 | -0.82 ± 0.51 | -1.08 ± 0.53 | 0.79 ± 0.51 | 1.23 ± 0.54 | -0.69 ± 0.51 |

*Cohen's d = (mean of selected line − mean of control) / pooled SD, computed separately for each block within each experiment; flies were compared with the control population of the same block.*

*Positive d = higher value in the selected line than in its matched control. 95% CI = d ± 1.96 × SE(d), with* $SE(d) = \sqrt{[(n1 + n2)/(n1 \cdot n2) + d^2/(2(n1 + n2))]}$.

**Table S4. Locomotor activity under LD12:12:** Two-way randomized block ANOVA for daytime and nighttime activity counts of virgin male (A) and female (B) chronotype flies under LD12:12.

A

| **day activity counts♂** | **Effect** | **SS** | $DF_n$ | **MS** | $DF_d$ | **MS error** | **F** | **p** |
|---|---|---|---|---|---|---|---|---|
| SELECTION | Fixed | 1975 | 2 | 987 | 6 | 15858.17 | 0.0623 | 0.940236 |
| BLOCK | Random | 55212 | 3 | 18404 | 6 | 15858.17 | 1.1605 | 0.39922 |
| SELECTION*BLOCK | Random | 95149 | 6 | 15858 | 0 | 0 | | |
| Error | | | 0 | | | | | |
| **night activity counts ♂** | **Effect** | **SS** | $DF_n$ | **MS** | $DF_d$ | **MS error** | **F** | **p** |
| SELECTION | Fixed | 6554 | 2 | 3277 | 6 | 5885.173 | 0.5568 | 0.600057 |
| BLOCK | Random | 20137 | 3 | 6712 | 6 | 5885.173 | 1.1405 | 0.405664 |
| SELECTION*BLOCK | Random | 35311 | 6 | 5885 | 0 | 0 | | |
| Error | | | 0 | | | | | |

B

| day activity counts ♀ | Effect | SS | $DF_n$ | MS | $DF_d$ | MS error | F | p |
|---|---|---|---|---|---|---|---|---|
| SELECTION | Fixed | 166830 | 2 | 83415 | 6 | 19115.47 | 4.3638 | 0.067619 |
| BLOCK | Random | 53698 | 3 | 17899 | 6 | 19115.47 | 0.9364 | 0.479303 |
| SELECTION*BLOCK | Random | 114693 | 6 | 19115 | 0 | 0 | | |
| Error | | | 0 | | | | | |
| **night activity counts ♀** | **Effect** | **SS** | **$DF_n$** | **MS** | **$DF_d$** | **MS error** | **F** | **p** |
| SELECTION | Fixed | 13282 | 2 | 6641 | 6 | 2781.74 | 2.3873 | 0.172683 |
| BLOCK | Random | 58574 | 3 | 19525 | 6 | 2781.74 | 7.0188 | 0.021774 |
| SELECTION*BLOCK | Random | 16690 | 6 | 2782 | 0 | 0 | | |
| Error | | | 0 | | | | | |

**Table S5. Nighttime sleep states:** Two-way randomized block ANOVA for duration, number of bouts and latency of short (A), intermediate (int) (B) and long (C) sleep at night in virgin male and female chronotype flies under LD12:12, with Tukey's HSD post hoc comparisons for parameters with a significant effect of selection.

A

| short-duration ♂ | Effect | SS | $DF_n$ | MS | $DF_d$ | MS error | F | p |
|---|---|---|---|---|---|---|---|---|
| Intercept | Fixed | 74599.43 | 1 | 74599.43 | 3 | 398.3371 | 187.277 | 0.001 |
| selection | Fixed | 2594.18 | 2 | 1297.09 | 6 | 149.6297 | 8.669 | 0.017 |
| block | Random | 1195.01 | 3 | 398.34 | 6 | 149.6297 | 2.662 | 0.142 |
| selection*block | Random | 897.78 | 6 | 149.63 | 0 | 0 | | |
| Error | | | 0 | | | | | |
| **short-bout numbers ♂** | **Effect** | **SS** | **$DF_n$** | **MS** | **$DF_d$** | **MS error** | **F** | **p** |
| Intercept | Fixed | 435.6023 | 1 | 435.6023 | 3 | 2.945625 | 147.881 | 0.001 |
| selection | Fixed | 11.4591 | 2 | 5.7296 | 6 | 1.252589 | 4.574 | 0.062 |
| block | Random | 8.8369 | 3 | 2.9456 | 6 | 1.252589 | 2.352 | 0.172 |
| selection*block | Random | 7.5155 | 6 | 1.2526 | 0 | 0 | | |
| Error | | | 0 | | | | | |
| **short-latency ♂** | **Effect** | **SS** | **$DF_n$** | **MS** | **$DF_d$** | **MS error** | **F** | **p** |
| Intercept | Fixed | 729575.4 | 1 | 729575.4 | 3 | 6393.438 | 114.113 | 0.002 |
| selection | Fixed | 16876.1 | 2 | 8438 | 6 | 973.264 | 8.670 | 0.017 |
| block | Random | 19180.3 | 3 | 6393.4 | 6 | 973.264 | 6.569 | 0.025 |
| selection*block | Random | 5839.6 | 6 | 973.3 | 0 | 0 | | |
| Error | | | 0 | | | | | |

| short-duration ♀ | Effect | SS | $DF_n$ | MS | $DF_d$ | MS error | F | p |
|---|---|---|---|---|---|---|---|---|
| Intercept | Fixed | 141009.4 | 1 | 141009.4 | 3 | 1063.382 | 132.605 | 0.001 |
| selection | Fixed | 3438.3 | 2 | 1719.2 | 6 | 248.075 | 6.930 | 0.028 |
| block | Random | 3190.1 | 3 | 1063.4 | 6 | 248.075 | 4.287 | 0.061 |
| selection*block | Random | 1488.4 | 6 | 248.1 | 0 | 0 | | |
| Error | | | 0 | | | | | |
| **short-bout numbers ♀** | **Effect** | **SS** | **$DF_n$** | **MS** | **$DF_d$** | **MS error** | **F** | **p** |
| Intercept | Fixed | 874.2452 | 1 | 874.2452 | 3 | 8.345869 | 104.752 | 0.002 |
| selection | Fixed | 15.8787 | 2 | 7.9394 | 6 | 1.907435 | 4.162 | 0.073 |
| block | Random | 25.0376 | 3 | 8.3459 | 6 | 1.907435 | 4.375 | 0.059 |
| selection*block | Random | 11.4446 | 6 | 1.9074 | 0 | 0 | | |
| Error | | | 0 | | | | | |
| **short-latency ♀** | **Effect** | **SS** | **$DF_n$** | **MS** | **$DF_d$** | **MS error** | **F** | **p** |
| Intercept | Fixed | 297654.1 | 1 | 297654.1 | 3 | 2746.145 | 108.390 | 0.002 |
| selection | Fixed | 10662.1 | 2 | 5331.1 | 6 | 735.863 | 7.245 | 0.025 |
| block | Random | 8238.4 | 3 | 2746.1 | 6 | 735.863 | 3.732 | 0.080 |
| selection*block | Random | 4415.2 | 6 | 735.9 | 0 | 0 | | |
| Error | | | 0 | | | | | |

B

| int-duration ♂ | Effect | SS | $DF_n$ | MS | $DF_d$ | MS error | F | p |
|---|---|---|---|---|---|---|---|---|
| Intercept | Fixed | 115967.7 | 1 | 115967.7 | 3 | 156.886 | 739.184 | 0.000 |
| selection | Fixed | 2421.1 | 2 | 1210.6 | 6 | 249.6128 | 4.850 | 0.056 |
| block | Random | 470.7 | 3 | 156.9 | 6 | 249.6128 | 0.628 | 0.623 |
| selection*block | Random | 1497.7 | 6 | 249.6 | 0 | 0 | | |
| Error | | | 0 | | | | | |
| **int-bout numbers ♂** | **Effect** | **SS** | **$DF_n$** | **MS** | **$DF_d$** | **MS error** | **F** | **p** |
| Intercept | Fixed | 62.51722 | 1 | 62.51722 | 3 | 0.085573 | 730.575 | 0.000 |
| selection | Fixed | 1.40134 | 2 | 0.70067 | 6 | 0.12643 | 5.542 | 0.043 |
| block | Random | 0.25672 | 3 | 0.08557 | 6 | 0.12643 | 0.677 | 0.597 |
| selection*block | Random | 0.75858 | 6 | 0.12643 | 0 | 0 | | |
| Error | | | 0 | | | | | |
| **int-latency ♂** | **Effect** | **SS** | **$DF_n$** | **MS** | **$DF_d$** | **MS error** | **F** | **p** |
| Intercept | Fixed | 1025944 | 1 | 1025944 | 3 | 3670.317 | 279.525 | 0.000 |
| selection | Fixed | 9805 | 2 | 4903 | 6 | 1644.144 | 2.982 | 0.126 |
| block | Random | 11011 | 3 | 3670 | 6 | 1644.144 | 2.232 | 0.185 |
| selection*block | Random | 9865 | 6 | 1644 | 0 | 0 | | |
| Error | | | 0 | | | | | |

| int-duration ♀ | Effect | SS | $DF_n$ | MS | $DF_d$ | MS error | F | p |
|---|---|---|---|---|---|---|---|---|
| Intercept | Fixed | 127773.7 | 1 | 127773.7 | 3 | 28.59831 | 4467.876 | 0.000 |
| selection | Fixed | 1883 | 2 | 941.5 | 6 | 43.19527 | 21.796 | 0.002 |
| block | Random | 85.8 | 3 | 28.6 | 6 | 43.19527 | 0.662 | 0.605 |
| selection*block | Random | 259.2 | 6 | 43.2 | 0 | 0 | | |
| Error | | | 0 | | | | | |
| **int-bout numbers ♀** | **Effect** | **SS** | **$DF_n$** | **MS** | **$DF_d$** | **MS error** | **F** | **p** |
| Intercept | Fixed | 70.42881 | 1 | 70.42881 | 3 | 0.019588 | 3595.528 | 0.000 |
| selection | Fixed | 1.28578 | 2 | 0.64289 | 6 | 0.031289 | 20.547 | 0.002 |
| block | Random | 0.05876 | 3 | 0.01959 | 6 | 0.031289 | 0.626 | 0.624 |
| selection*block | Random | 0.18773 | 6 | 0.03129 | 0 | 0 | | |
| Error | | | 0 | | | | | |
| **int-latency ♀** | **Effect** | **SS** | **$DF_n$** | **MS** | **$DF_d$** | **MS error** | **F** | **p** |
| Intercept | Fixed | 671098.9 | 1 | 671098.9 | 3 | 1216.905 | 551.480 | 0.000 |
| selection | Fixed | 4546.6 | 2 | 2273.3 | 6 | 1155.118 | 1.968 | 0.220 |
| block | Random | 3650.7 | 3 | 1216.9 | 6 | 1155.118 | 1.054 | 0.435 |
| selection*block | Random | 6930.7 | 6 | 1155.1 | 0 | 0 | | |
| Error | | | 0 | | | | | |

C

| long-duration ♂ | Effect | SS | $DF_n$ | MS | $DF_d$ | MS error | F | p |
|---|---|---|---|---|---|---|---|---|
| Intercept | Fixed | 1660853 | 1 | 1660853 | 3 | 5055.559 | 328.520 | 0.000 |
| selection | Fixed | 21398 | 2 | 10699 | 6 | 790.058 | 13.542 | 0.006 |
| block | Random | 15167 | 3 | 5056 | 6 | 790.058 | 6.399 | 0.027 |
| selection*block | Random | 4740 | 6 | 790 | 0 | 0 | | |
| Error | | | 0 | | | | | |
| **long-bout numbers ♂** | **Effect** | **SS** | **$DF_n$** | **MS** | **$DF_d$** | **MS error** | **F** | **p** |
| Intercept | Fixed | 80.6588 | 1 | 80.6588 | 3 | 0.013711 | 5882.912 | 0.000 |
| selection | Fixed | 0.15921 | 2 | 0.07961 | 6 | 0.080649 | 0.987 | 0.426 |
| block | Random | 0.04113 | 3 | 0.01371 | 6 | 0.080649 | 0.170 | 0.913 |
| selection*block | Random | 0.4839 | 6 | 0.08065 | 0 | 0 | | |
| Error | | | 0 | | | | | |
| **long-latency ♂** | **Effect** | **SS** | **$DF_n$** | **MS** | **$DF_d$** | **MS error** | **F** | **p** |
| Intercept | Fixed | 102857.7 | 1 | 102857.7 | 3 | 2120.258 | 48.512 | 0.006 |
| selection | Fixed | 3242 | 2 | 1621 | 6 | 209.996 | 7.719 | 0.022 |
| block | Random | 6360.8 | 3 | 2120.3 | 6 | 209.996 | 10.097 | 0.009 |
| selection*block | Random | 1260 | 6 | 210 | 0 | 0 | | |

| Error | | | 0 |
|---|---|---|---|

| long-duration ♀ | Effect | SS | $DF_n$ | MS | $DF_d$ | MS error | F | p |
|---|---|---|---|---|---|---|---|---|
| Intercept | Fixed | 935533.7 | 1 | 935533.7 | 3 | 5345.601 | 175.010 | 0.001 |
| selection | Fixed | 22126.3 | 2 | 11063.1 | 6 | 644.56 | 17.164 | 0.003 |
| block | Random | 16036.8 | 3 | 5345.6 | 6 | 644.56 | 8.293 | 0.015 |
| selection*block | Random | 3867.4 | 6 | 644.6 | 0 | 0 | | |
| Error | | | 0 | | | | | |
| **long-bout numbers ♀** | **Effect** | **SS** | **$DF_n$** | **MS** | **$DF_d$** | **MS error** | **F** | **p** |
| Intercept | Fixed | 59.75147 | 1 | 59.75147 | 3 | 0.232513 | 256.981 | 0.001 |
| selection | Fixed | 0.55956 | 2 | 0.27978 | 6 | 0.057521 | 4.864 | 0.056 |
| block | Random | 0.69754 | 3 | 0.23251 | 6 | 0.057521 | 4.042 | 0.069 |
| selection*block | Random | 0.34513 | 6 | 0.05752 | 0 | 0 | | |
| Error | | | 0 | | | | | |
| **long-latency ♀** | **Effect** | **SS** | **$DF_n$** | **MS** | **$DF_d$** | **MS error** | **F** | **p** |
| Intercept | Fixed | 253830.8 | 1 | 253830.8 | 3 | 570.1878 | 445.171 | 0.000 |
| selection | Fixed | 9444.5 | 2 | 4722.3 | 6 | 842.1858 | 5.607 | 0.042 |
| block | Random | 1710.6 | 3 | 570.2 | 6 | 842.1858 | 0.677 | 0.597 |
| selection*block | Random | 5053.1 | 6 | 842.2 | 0 | 0 | | |
| Error | | | 0 | | | | | |

| Tukey's HSD test | | | | | | | | | | | |
|---|---|---|---|---|---|---|---|---|---|---|---|
| | Males | | | | | Females | | | | | |
| | Short | | Intermediate | Long | | Short | | Intermediate | | Long | |
| Mean difference | Duration | Latency | Bout numbers | Duration | Latency | Duration | Latency | Duration | Bout numbers | Duration | Latency |
| *early vs control* | 35.9276 | -86.5232 | 0.8341 | -103.3234 | 35.5551 | 38.7372 | -63.1144 | 20.6873 | 0.5615 | -100.5930 | 52.6112 |
| *early vs late* | 15.7913 | -69.9793 | 0.3561 | -55.8466 | 34.1365 | 32.1724 | -63.3491 | 29.9685 | 0.7764 | -76.9073 | 64.5906 |
| *control vs late* | -20.1363 | 16.5439 | -0.4780 | 47.4768 | -1.4186 | -6.5647 | -0.2347 | 9.2812 | 0.2150 | 23.6857 | 11.9794 |
| Minimum significant difference | 26.5380 | 67.6823 | 0.7714 | 60.9802 | 31.4387 | 34.1705 | 58.8516 | 14.2586 | 0.3838 | 55.0797 | 62.9598 |
| q critical | 4.339 | 4.339 | 4.339 | 4.339 | 4.339 | 4.339 | 4.339 | 4.339 | 4.339 | 4.339 | 4.339 |
| K, N | 3, 4 | 3, 4 | 3, 4 | 3, 4 | 3, 4 | 3, 4 | 3, 4 | 3, 4 | 3, 4 | 3, 4 | 3, 4 |

**Table S6. Daytime sleep states:** Two-way randomized block ANOVA for duration, number of bouts and latency of short (A), intermediate (int) (B) and long (C) sleep at day in virgin male and female chronotype flies under LD12:12.

A

| short-duration ♂ | Effect | SS | $DF_n$ | MS | $DF_d$ | MS error | F | p |
|---|---|---|---|---|---|---|---|---|
| Intercept | Fixed | 159950.7 | 1 | 159950.7 | 3 | 5.1513 | 31050.38 | 0 |
| Selection | Fixed | 1028.8 | 2 | 514.4 | 6 | 528.1167 | 0.97 | 0.43019 |
| Block | Random | 15.5 | 3 | 5.2 | 6 | 528.1167 | 0.01 | 0.998529 |
| Selection*Block | Random | 3168.7 | 6 | 528.1 | 0 | 0 | | |
| Error | | | 0 | | | | | |
| **short-bout numbers ♂** | **Effect** | **SS** | **$DF_n$** | **MS** | **$DF_d$** | **MS error** | **F** | **p** |
| Intercept | Fixed | 1172.487 | 1 | 1172.487 | 3 | 0.670542 | 1748.566 | 0.00003 |
| Selection | Fixed | 3.319 | 2 | 1.659 | 6 | 6.150317 | 0.27 | 0.772322 |
| Block | Random | 2.012 | 3 | 0.671 | 6 | 6.150317 | 0.109 | 0.951709 |
| Selection*Block | Random | 36.902 | 6 | 6.15 | 0 | 0 | | |
| Error | | | 0 | | | | | |
| **short-latency ♂** | **Effect** | **SS** | **$DF_n$** | **MS** | **$DF_d$** | **MS error** | **F** | **p** |
| Intercept | Fixed | 125626.4 | 1 | 125626.4 | 3 | 65.6439 | 1913.755 | 0.000026 |
| Selection | Fixed | 340.7 | 2 | 170.4 | 6 | 655.8809 | 0.26 | 0.779494 |
| Block | Random | 196.9 | 3 | 65.6 | 6 | 655.8809 | 0.1 | 0.957041 |
| Selection*Block | Random | 3935.3 | 6 | 655.9 | 0 | 0 | | |
| Error | | | 0 | | | | | |

| short-duration ♀ | Effect | SS | $DF_n$ | MS | $DF_d$ | MS error | F | p |
|---|---|---|---|---|---|---|---|---|

| | | | | | | | | |
|---|---|---|---|---|---|---|---|---|
| Intercept | Fixed | 92416.98 | 1 | 92416.98 | 3 | 179.3753 | 515.2159 | 0.000187 |
| Selection | Fixed | 2504.1 | 2 | 1252.05 | 6 | 811.5483 | 1.5428 | 0.288002 |
| Block | Random | 538.13 | 3 | 179.38 | 6 | 811.5483 | 0.221 | 0.878472 |
| Selection*Block | Random | 4869.29 | 6 | 811.55 | 0 | 0 | | |
| Error | | | 0 | | | | | |
| **short-bout numbers ♀** | **Effect** | **SS** | **$DF_n$** | **MS** | **$DF_d$** | **MS error** | **F** | **p** |
| Intercept | Fixed | 901.1507 | 1 | 901.1507 | 3 | 0.795422 | 1132.921 | 0.000058 |
| Selection | Fixed | 23.5317 | 2 | 11.7658 | 6 | 6.574418 | 1.79 | 0.245728 |
| Block | Random | 2.3863 | 3 | 0.7954 | 6 | 6.574418 | 0.121 | 0.944392 |
| Selection*Block | Random | 39.4465 | 6 | 6.5744 | 0 | 0 | | |
| Error | | | 0 | | | | | |
| **short-latency ♀** | **Effect** | **SS** | **$DF_n$** | **MS** | **$DF_d$** | **MS error** | **F** | **p** |
| Intercept | Fixed | 414473.5 | 1 | 414473.5 | 3 | 507.348 | 816.9407 | 0.000094 |
| Selection | Fixed | 2425.5 | 2 | 1212.8 | 6 | 1731.446 | 0.7004 | 0.532852 |
| Block | Random | 1522 | 3 | 507.3 | 6 | 1731.446 | 0.293 | 0.829394 |
| Selection*Block | Random | 10388.7 | 6 | 1731.4 | 0 | 0 | | |
| Error | | | 0 | | | | | |

B

| int-duration ♂ | Effect | SS | $DF_n$ | MS | $DF_d$ | MS error | F | p |
|---|---|---|---|---|---|---|---|---|
| Intercept | Fixed | 83024.29 | 1 | 83024.29 | 3 | 42.14859 | 1969.8 | 0.000025 |
| Selection | Fixed | 115.72 | 2 | 57.86 | 6 | 61.73541 | 0.937 | 0.44237 |
| Block | Random | 126.45 | 3 | 42.15 | 6 | 61.73541 | 0.683 | 0.594263 |
| Selection*Block | Random | 370.41 | 6 | 61.74 | 0 | 0 | | |
| Error | | | 0 | | | | | |
| **int-bout numbers ♂** | **Effect** | **SS** | **$DF_n$** | **MS** | **$DF_d$** | **MS error** | **F** | **p** |
| Intercept | Fixed | 50.11012 | 1 | 50.11012 | 3 | 0.018626 | 2690.329 | 0.000016 |
| Selection | Fixed | 0.0576 | 2 | 0.0288 | 6 | 0.04542 | 0.634 | 0.562597 |
| Block | Random | 0.05588 | 3 | 0.01863 | 6 | 0.04542 | 0.41 | 0.751819 |
| Selection*Block | Random | 0.27252 | 6 | 0.04542 | 0 | 0 | | |
| Error | | | 0 | | | | | |
| **int-latency ♂** | **Effect** | **SS** | **$DF_n$** | **MS** | **$DF_d$** | **MS error** | **F** | **p** |
| Intercept | Fixed | 508170.4 | 1 | 508170.4 | 3 | 362.5604 | 1401.616 | 0.000042 |
| Selection | Fixed | 52.9 | 2 | 26.5 | 6 | 893.7524 | 0.03 | 0.970975 |
| Block | Random | 1087.7 | 3 | 362.6 | 6 | 893.7524 | 0.406 | 0.754661 |
| Selection*Block | Random | 5362.5 | 6 | 893.8 | 0 | 0 | | |
| Error | | | 0 | | | | | |

| int-duration ♀ | Effect | SS | $DF_n$ | MS | $DF_d$ | MS error | F | p |
|---|---|---|---|---|---|---|---|---|
| Intercept | Fixed | 17442.53 | 1 | 17442.53 | 3 | 117.7284 | 148.1591 | 0.001194 |
| Selection | Fixed | 170.51 | 2 | 85.25 | 6 | 278.5296 | 0.3061 | 0.747171 |
| Block | Random | 353.19 | 3 | 117.73 | 6 | 278.5296 | 0.4227 | 0.743767 |
| Selection*Block | Random | 1671.18 | 6 | 278.53 | 0 | 0 | | |
| Error | | | 0 | | | | | |
| **int-bout numbers ♀** | **Effect** | **SS** | **$DF_n$** | **MS** | **$DF_d$** | **MS error** | **F** | **p** |
| Intercept | Fixed | 11.29533 | 1 | 11.29533 | 3 | 0.065454 | 172.5688 | 0.000953 |
| Selection | Fixed | 0.1128 | 2 | 0.0564 | 6 | 0.158244 | 0.3564 | 0.714075 |
| Block | Random | 0.19636 | 3 | 0.06545 | 6 | 0.158244 | 0.4136 | 0.749546 |
| Selection*Block | Random | 0.94946 | 6 | 0.15824 | 0 | 0 | | |
| Error | | | 0 | | | | | |
| **int-latency ♀** | **Effect** | **SS** | **$DF_n$** | **MS** | **$DF_d$** | **MS error** | **F** | **p** |
| Intercept | Fixed | 1041983 | 1 | 1041983 | 3 | 382.7667 | 2722.239 | 0.000016 |
| Selection | Fixed | 245 | 2 | 122 | 6 | 569.9708 | 0.215 | 0.812867 |
| Block | Random | 1148 | 3 | 383 | 6 | 569.9708 | 0.672 | 0.600007 |
| Selection*Block | Random | 3420 | 6 | 570 | 0 | 0 | | |
| Error | | | 0 | | | | | |

C

| **long-duration ♂** | **Effect** | **SS** | **$DF_n$** | **MS** | **$DF_d$** | **MS error** | **F** | **p** |
|---|---|---|---|---|---|---|---|---|
| Intercept | Fixed | 327644.8 | 1 | 327644.8 | 3 | 682.988 | 479.7225 | 0.000208 |
| Selection | Fixed | 2877.5 | 2 | 1438.7 | 6 | 1645.083 | 0.8746 | 0.464186 |
| Block | Random | 2049 | 3 | 683 | 6 | 1645.083 | 0.4152 | 0.748559 |
| Selection*Block | Random | 9870.5 | 6 | 1645.1 | 0 | 0 | | |
| Error | | | 0 | | | | | |
| **long-bout numbers ♂** | **Effect** | **SS** | **$DF_n$** | **MS** | **$DF_d$** | **MS error** | **F** | **p** |
| Intercept | Fixed | 22.61716 | 1 | 22.61716 | 3 | 0.023932 | 945.0554 | 0.000076 |
| Selection | Fixed | 0.12352 | 2 | 0.06176 | 6 | 0.097824 | 0.6313 | 0.563852 |
| Block | Random | 0.0718 | 3 | 0.02393 | 6 | 0.097824 | 0.2446 | 0.862367 |
| Selection*Block | Random | 0.58694 | 6 | 0.09782 | 0 | 0 | | |
| Error | | | 0 | | | | | |
| **long-latency ♂** | **Effect** | **SS** | **$DF_n$** | **MS** | **$DF_d$** | **MS error** | **F** | **p** |
| Intercept | Fixed | 432048 | 1 | 432048 | 3 | 1099.798 | 392.8429 | 0.000281 |
| Selection | Fixed | 512.5 | 2 | 256.2 | 6 | 237.161 | 1.0804 | 0.397426 |
| Block | Random | 3299.4 | 3 | 1099.8 | 6 | 237.161 | 4.6374 | 0.052617 |
| Selection*Block | Random | 1423 | 6 | 237.2 | 0 | 0 | | |
| Error | | | 0 | | | | | |

| **long-duration ♀** | **Effect** | **SS** | **$DF_n$** | **MS** | **$DF_d$** | **MS error** | **F** | **p** |
|---|---|---|---|---|---|---|---|---|
| Intercept | Fixed | 48515.08 | 1 | 48515.08 | 3 | 1654.484 | 29.32339 | 0.012352 |
| Selection | Fixed | 1285.84 | 2 | 642.92 | 6 | 280.303 | 2.29366 | 0.18201 |
| Block | Random | 4963.45 | 3 | 1654.48 | 6 | 280.303 | 5.90249 | 0.031904 |
| Selection*Block | Random | 1681.82 | 6 | 280.3 | 0 | 0 | | |
| Error | | | 0 | | | | | |
| **long-bout numbers ♀** | **Effect** | **SS** | **$DF_n$** | **MS** | **$DF_d$** | **MS error** | **F** | **p** |
| Intercept | Fixed | 5.361838 | 1 | 5.361838 | 3 | 0.116656 | 45.96295 | 0.006559 |
| Selection | Fixed | 0.064132 | 2 | 0.032066 | 6 | 0.018887 | 1.69782 | 0.260421 |
| Block | Random | 0.349967 | 3 | 0.116656 | 6 | 0.018887 | 6.17664 | 0.028915 |
| Selection*Block | Random | 0.11332 | 6 | 0.018887 | 0 | 0 | | |
| Error | | | 0 | | | | | |
| **long-latency ♀** | **Effect** | **SS** | **$DF_n$** | **MS** | **$DF_d$** | **MS error** | **F** | **p** |
| Intercept | Fixed | 567947.9 | 1 | 567947.9 | 3 | 167.324 | 3394.291 | 0.000011 |
| Selection | Fixed | 4021.3 | 2 | 2010.6 | 6 | 2596.499 | 0.774 | 0.502147 |
| Block | Random | 502 | 3 | 167.3 | 6 | 2596.499 | 0.064 | 0.976765 |
| Selection*Block | Random | 15579 | 6 | 2596.5 | 0 | 0 | | |
| Error | | | 0 | | | | | |

**Table S7. Sleep across light intensities:** Randomized block ANOVA (selection and light intensity as fixed factors and block as a random factor) for daytime (A) and nighttime (B) sleep parameters of virgin male chronotype flies under 1, 10, 100 and 1000 lux. (C) Tukey's HSD post hoc comparisons for the main effect of light intensity.

A

| **Day duration** | Effect | SS | $DF_n$ | MS | $DF_d$ | MS error | F | p |
|---|---|---|---|---|---|---|---|---|
| Intercept | Fixed | 6741623 | 1 | 6741623 | 3 | 8884.617 | 758.7973 | 0.000105 |
| SELECTION | Fixed | 34770 | 2 | 17385 | 6 | 3420.539 | 5.0826 | **0.051135** |
| INTENSITY | Fixed | 13348 | 3 | 4449 | 9 | 741.633 | 5.9993 | 0.015721 |
| BLOCK | Random | 26654 | 3 | 8885 | 7.87416 | 3981.239 | 2.2316 | 0.163088 |
| SELECTION*INTENSITY | Fixed | 2051 | 6 | 342 | 18 | 180.934 | 1.8891 | 0.138066 |
| SELECTION*BLOCK | Random | 20523 | 6 | 3421 | 18 | 180.934 | 18.9049 | 0.000001 |
| INTENSITY*BLOCK | Random | 6675 | 9 | 742 | 18 | 180.934 | 4.0989 | 0.005267 |
| SELECTION*INTENSITY*BLOCK | Random | 3257 | 18 | 181 | 0 | 0 | | |
| Error | | | 0 | | | | | |
| **Day latency** | Effect | SS | $DF_n$ | MS | $DF_d$ | MS error | F | p |
| Intercept | Fixed | 213665.9 | 1 | 213665.9 | 3 | 519.1837 | 411.5421 | 0.000262 |
| SELECTION | Fixed | 2042.9 | 2 | 1021.5 | 6 | 565.3548 | 1.8068 | 0.24311 |

| | | | | | | | | |
|---|---|---|---|---|---|---|---|---|
| INTENSITY | Fixed | 2204.5 | 3 | 734.8 | 9 | 134.1547 | 5.4776 | 0.020327 |
| BLOCK | Random | 1557.6 | 3 | 519.2 | 7.24972 | 634.3558 | 0.8184 | 0.522337 |
| SELECTION*INTENSITY | Fixed | 202.5 | 6 | 33.7 | 18 | 65.1538 | 0.518 | 0.787145 |
| SELECTION*BLOCK | Random | 3392.1 | 6 | 565.4 | 18 | 65.1538 | 8.6772 | 0.000159 |
| INTENSITY*BLOCK | Random | 1207.4 | 9 | 134.2 | 18 | 65.1538 | 2.059 | 0.09187 |
| SELECTION*INTENSITY*BLOCK | Random | 1172.8 | 18 | 65.2 | 0 | 0 | | |
| Error | | | 0 | | | | | |
| **Day bout numbers** | Effect | SS | $DF_n$ | MS | $DF_d$ | MS error | F | p |
| Intercept | Fixed | 10159.84 | 1 | 10159.84 | 3 | 5.194117 | 1956.028 | 0.000025 |
| SELECTION | Fixed | 22.89 | 2 | 11.45 | 6 | 7.545743 | 1.517 | 0.292991 |
| INTENSITY | Fixed | 27.22 | 3 | 9.07 | 9 | 0.781447 | 11.61 | 0.001901 |
| BLOCK | Random | 15.58 | 3 | 5.19 | 3.70413 | 6.039792 | 0.86 | 0.534937 |
| SELECTION*INTENSITY | Fixed | 21.59 | 6 | 3.6 | 18 | 2.287397 | 1.573 | 0.211933 |
| SELECTION*BLOCK | Random | 45.27 | 6 | 7.55 | 18 | 2.287397 | 3.299 | 0.022771 |
| INTENSITY*BLOCK | Random | 7.03 | 9 | 0.78 | 18 | 2.287397 | 0.342 | 0.948319 |
| SELECTION*INTENSITY*BLOCK | Random | 41.17 | 18 | 2.29 | 0 | 0 | | |
| Error | | | 0 | | | | | |
| **Day median bout duration** | Effect | SS | $DF_n$ | MS | $DF_d$ | MS error | F | p |
| Intercept | Fixed | 15218.91 | 1 | 15218.91 | 3 | 4.39307 | 3464.302 | 0.000011 |
| selection | Fixed | 2.44 | 2 | 1.22 | 6 | 21.04648 | 0.058 | 0.94413 |
| intensity | Fixed | 18.25 | 3 | 6.08 | 9 | 3.10949 | 1.957 | 0.191129 |
| block | Random | 13.18 | 3 | 4.39 | 5.27998 | 20.01265 | 0.22 | 0.879166 |
| selection*intensity | Fixed | 45.08 | 6 | 7.51 | 18 | 4.14331 | 1.813 | 0.152961 |
| selection*block | Random | 126.28 | 6 | 21.05 | 18 | 4.14331 | 5.08 | 0.003279 |
| intensity*block | Random | 27.99 | 9 | 3.11 | 18 | 4.14331 | 0.75 | 0.660737 |
| selection*intensity*block | Random | 74.58 | 18 | 4.14 | 0 | 0 | | |
| Error | | | 0 | | | | | |

B

| **Night duration** | Effect | SS | $DF_n$ | MS | $DF_d$ | MS error | F | p |
|---|---|---|---|---|---|---|---|---|
| Intercept | Fixed | 17296173 | 1 | 17296173 | 3 | 7377.759 | 2344.367 | 0.000019 |
| SELECTION | Fixed | 2051 | 2 | 1026 | 6 | 1992.66 | 0.515 | 0.621859 |
| INTENSITY | Fixed | 3796 | 3 | 1265 | 9 | 204.365 | 6.192 | 0.014347 |
| BLOCK | Random | 22133 | 3 | 7378 | 6.24284 | 2041.748 | 3.613 | 0.081537 |
| SELECTION*INTENSITY | Fixed | 114 | 6 | 19 | 18 | 155.278 | 0.122 | 0.992221 |
| SELECTION*BLOCK | Random | 11956 | 6 | 1993 | 18 | 155.278 | 12.833 | 0.000012 |
| INTENSITY*BLOCK | Random | 1839 | 9 | 204 | 18 | 155.278 | 1.316 | 0.295311 |
| SELECTION*INTENSITY*BLOCK | Random | 2795 | 18 | 155 | 0 | 0 | | |
| Error | | | 0 | | | | | |
| **Night latency** | Effect | SS | $DF_n$ | MS | $DF_d$ | MS error | F | p |
| Intercept | Fixed | 11490.33 | 1 | 11490.33 | 3 | 107.3308 | 107.0553 | 0.001926 |
| SELECTION | Fixed | 40.14 | 2 | 20.07 | 6 | 36.7661 | 0.5459 | 0.605612 |
| INTENSITY | Fixed | 848.94 | 3 | 282.98 | 9 | 9.8363 | 28.7692 | 0.000061 |
| BLOCK | Random | 321.99 | 3 | 107.33 | 7.40763 | 41.9227 | 2.5602 | 0.13358 |
| SELECTION*INTENSITY | Fixed | 33.36 | 6 | 5.56 | 18 | 4.6796 | 1.188 | 0.356296 |
| SELECTION*BLOCK | Random | 220.6 | 6 | 36.77 | 18 | 4.6796 | 7.8567 | 0.000292 |
| INTENSITY*BLOCK | Random | 88.53 | 9 | 9.84 | 18 | 4.6796 | 2.1019 | 0.085945 |
| SELECTION*INTENSITY*BLOCK | Random | 84.23 | 18 | 4.68 | 0 | 0 | | |
| Error | | | 0 | | | | | |
| **Night bout number** | Effect | SS | $DF_n$ | MS | $DF_d$ | MS error | F | p |
| Intercept | Fixed | 6046.338 | 1 | 6046.338 | 3 | 11.72856 | 515.5225 | 0.000187 |
| SELECTION | Fixed | 11.975 | 2 | 5.988 | 6 | 6.59504 | 0.9079 | 0.452415 |
| INTENSITY | Fixed | 5.327 | 3 | 1.776 | 9 | 1.71768 | 1.0337 | 0.423058 |
| BLOCK | Random | 35.186 | 3 | 11.729 | 7.6881 | 7.64478 | 1.5342 | 0.281499 |
| SELECTION*INTENSITY | Fixed | 2.213 | 6 | 0.369 | 18 | 0.66794 | 0.5523 | 0.762044 |
| SELECTION*BLOCK | Random | 39.57 | 6 | 6.595 | 18 | 0.66794 | 9.8738 | 0.00007 |
| INTENSITY*BLOCK | Random | 15.459 | 9 | 1.718 | 18 | 0.66794 | 2.5716 | 0.042083 |

| | | | | | | | | |
|---|---|---|---|---|---|---|---|---|
| SELECTION*INTENSITY*BLOCK | Random | 12.023 | 18 | 0.668 | 0 | 0 | | |
| Error | | | 0 | | | | | |
| **Night median bout duration** | Effect | SS | $DF_n$ | MS | $DF_d$ | MS error | F | p |
| Intercept | Fixed | 157458.7 | 1 | 157458.7 | 3 | 2161.493 | 72.84718 | 0.003379 |
| selection | Fixed | 1697.9 | 2 | 848.9 | 6 | 697.637 | 1.21687 | 0.360076 |
| intensity | Fixed | 403.5 | 3 | 134.5 | 9 | 132.29 | 1.01662 | 0.429717 |
| block | Random | 6484.5 | 3 | 2161.5 | 6.42439 | 732.791 | 2.94967 | 0.114564 |
| selection*intensity | Fixed | 653.7 | 6 | 109 | 18 | 97.136 | 1.12163 | 0.388877 |
| selection*block | Random | 4185.8 | 6 | 697.6 | 18 | 97.136 | 7.18203 | 0.000498 |
| intensity*block | Random | 1190.6 | 9 | 132.3 | 18 | 97.136 | 1.3619 | 0.275044 |
| selection*intensity*block | Random | 1748.5 | 18 | 97.1 | 0 | 0 | | |
| Error | | | 0 | | | | | |

C

| Tukey's HSD test - main effect of intensity | | | | | |
|---|---|---|---|---|---|
| | Daytime | | | Nighttime | |
| **Mean difference** | **Duration** | **Latency** | **Bout number** | **Duration** | **Latency** |
| **1 lux-10 lux** | -30.3412 | 10.0324 | 0.7528 | -7.5263 | -1.6076 |
| **1 lux-100 lux** | -37.0436 | 0.5564 | 1.1185 | 15.3521 | -7.9582 |
| **1 lux-1000 lux** | -43.5350 | -9.1262 | -0.8449 | 10.3068 | -10.0520 |
| **10 lux-100 lux** | -6.7024 | -9.4760 | 0.3657 | 22.8784 | -6.3506 |
| **10 lux-1000 lux** | -13.1938 | -19.1586 | -1.5977 | 17.8331 | -8.4444 |
| **100 lux-1000 lux** | -6.4914 | -9.6826 | -1.9634 | -5.0453 | -2.0938 |
| Minimum significant difference | 34.7080 | 14.762 | 1.1270 | 18.219 | 3.9970 |
| q critical | 4.4149 | 4.4149 | 4.4149 | 4.4149 | 4.4149 |
| K, N | 4,12 | 4,12 | 4,12 | 4,12 | 4,12 |

**Table S8. Sleep under DDTC:** Two-way randomized block ANOVA for sleep parameters of virgin male chronotype flies under DDTC: midpoint of sleep rise and behavioural night parameters (A), sleep parameters during the 12 h thermophase (B) and cryophase (C) and midpoint of rise in long sleep (D). (E) Tukey's HSD post hoc comparisons for parameters with a significant effect of selection.

A

| **Midpoint of sleep rise** | **Effect** | **SS** | **$DF_n$** | **MS** | **$DF_d$** | **MS error** | **F** | **p** |
|---|---|---|---|---|---|---|---|---|
| Intercept | Fixed | 1297.897 | 1 | 1297.897 | 3 | 0.030586 | 42433.820 | 0.000 |
| selection | Fixed | 5.864 | 2 | 2.932 | 6 | 0.102879 | 28.500 | 0.001 |
| block | Random | 0.092 | 3 | 0.031 | 6 | 0.102879 | 0.300 | 0.826 |
| selection*block | Random | 0.617 | 6 | 0.103 | 0 | 0 | | |
| Error | | | 0 | | | | | |
| **Night bout numbers** | **Effect** | **SS** | **$DF_n$** | **MS** | **$DF_d$** | **MS error** | **F** | **p** |
| Intercept | Fixed | 2401.677 | 1 | 2401.677 | 3 | 3.158463 | 760.394 | 0.000 |
| selection | Fixed | 32.286 | 2 | 16.143 | 6 | 5.119106 | 3.154 | 0.116 |
| block | Random | 9.475 | 3 | 3.158 | 6 | 5.119106 | 0.617 | 0.629 |
| selection*block | Random | 30.715 | 6 | 5.119 | 0 | 0 | | |
| Error | | | 0 | | | | | |
| **Night median bout duration** | **Effect** | **SS** | **$DF_n$** | **MS** | **$DF_d$** | **MS error** | **F** | **p** |
| Intercept | Fixed | 25395.35 | 1 | 25395.35 | 3 | 498.5224 | 50.941 | 0.006 |
| selection | Fixed | 1319.62 | 2 | 659.81 | 6 | 123.5285 | 5.341 | 0.047 |
| block | Random | 1495.57 | 3 | 498.52 | 6 | 123.5285 | 4.036 | 0.069 |
| selection*block | Random | 741.17 | 6 | 123.53 | 0 | 0 | | |
| Error | | | 0 | | | | | |

B

| **Thermo- Duration** | **Effect** | **SS** | **$DF_n$** | **MS** | **$DF_d$** | **MS error** | **F** | **p** |
|---|---|---|---|---|---|---|---|---|
| Intercept | Fixed | 1379506 | 1 | 1379506 | 3 | 192.284 | 7174.308 | 0.000 |
| selection | Fixed | 16989 | 2 | 8494 | 6 | 2528.986 | 3.359 | 0.105 |
| block | Random | 577 | 3 | 192 | 6 | 2528.986 | 0.076 | 0.971 |
| selection*block | Random | 15174 | 6 | 2529 | 0 | 0 | | |
| Error | | | 0 | | | | | |

| Thermo- Latency | Effect | SS | $DF_n$ | MS | $DF_d$ | MS error | F | p |
|---|---|---|---|---|---|---|---|---|
| Intercept | Fixed | 2253.258 | 1 | 2253.258 | 3 | 50.5302 | 44.592 | 0.007 |
| selection | Fixed | 47.093 | 2 | 23.546 | 6 | 175.2705 | 0.134 | 0.877 |
| block | Random | 151.591 | 3 | 50.53 | 6 | 175.2705 | 0.288 | 0.833 |
| selection*block | Random | 1051.623 | 6 | 175.271 | 0 | 0 | | |
| Error | | | 0 | | | | | |
| **Thermo- Bout number** | **Effect** | **SS** | **$DF_n$** | **MS** | **$DF_d$** | **MS error** | **F** | **p** |
| Intercept | Fixed | 3551.851 | 1 | 3551.851 | 3 | 8.062866 | 440.520 | 0.000 |
| selection | Fixed | 6.754 | 2 | 3.377 | 6 | 1.556352 | 2.170 | 0.195 |
| block | Random | 24.189 | 3 | 8.063 | 6 | 1.556352 | 5.181 | 0.042 |
| selection*block | Random | 9.338 | 6 | 1.556 | 0 | 0 | | |
| Error | | | 0 | | | | | |
| **Thermo- median bout duration** | **Effect** | **SS** | **$DF_n$** | **MS** | **$DF_d$** | **MS error** | **F** | **p** |
| Intercept | Fixed | 5321.7 | 1 | 5321.7 | 3 | 11.33047 | 469.680 | 0.000 |
| selection | Fixed | 38.003 | 2 | 19.002 | 6 | 10.01352 | 1.898 | 0.230 |
| block | Random | 33.991 | 3 | 11.33 | 6 | 10.01352 | 1.131 | 0.409 |
| selection*block | Random | 60.081 | 6 | 10.014 | 0 | 0 | | |
| Error | | | 0 | | | | | |

C

| Cryo- Duration | Effect | SS | $DF_n$ | MS | $DF_d$ | MS error | F | p |
|---|---|---|---|---|---|---|---|---|
| Intercept | Fixed | 4987538 | 1 | 4987538 | 3 | 801.6408 | 6221.662 | 0.000 |
| selection | Fixed | 105 | 2 | 52 | 6 | 160.2181 | 0.327 | 0.733 |
| block | Random | 2405 | 3 | 802 | 6 | 160.2181 | 5.003 | 0.045 |
| selection*block | Random | 961 | 6 | 160 | 0 | 0 | | |
| Error | | | 0 | | | | | |
| **Cryo- latency** | **Effect** | **SS** | **$DF_n$** | **MS** | **$DF_d$** | **MS error** | **F** | **p** |
| Intercept | Fixed | 1035.204 | 1 | 1035.204 | 3 | 4.803 | 215.533 | 0.001 |
| selection | Fixed | 371.128 | 2 | 185.564 | 6 | 31.80599 | 5.834 | 0.039 |
| block | Random | 14.409 | 3 | 4.803 | 6 | 31.80599 | 0.151 | 0.925 |
| selection*block | Random | 190.836 | 6 | 31.806 | 0 | 0 | | |
| Error | | | 0 | | | | | |
| **Cryo- bout number** | **Effect** | **SS** | **$DF_n$** | **MS** | **$DF_d$** | **MS error** | **F** | **p** |
| Intercept | Fixed | 1466.59 | 1 | 1466.59 | 3 | 3.900196 | 376.030 | 0.000 |
| selection | Fixed | 7.383 | 2 | 3.691 | 6 | 3.6975 | 0.998 | 0.422 |
| block | Random | 11.701 | 3 | 3.9 | 6 | 3.6975 | 1.055 | 0.435 |
| selection*block | Random | 22.185 | 6 | 3.697 | 0 | 0 | | |
| Error | | | 0 | | | | | |
| **Cryo- median bout duration** | **Effect** | **SS** | **$DF_n$** | **MS** | **$DF_d$** | **MS error** | **F** | **p** |
| Intercept | Fixed | 73068.25 | 1 | 73068.25 | 3 | 1029.763 | 70.956 | 0.004 |
| selection | Fixed | 200.03 | 2 | 100.02 | 6 | 965.017 | 0.104 | 0.903 |
| block | Random | 3089.29 | 3 | 1029.76 | 6 | 965.017 | 1.067 | 0.430 |
| selection*block | Random | 5790.1 | 6 | 965.02 | 0 | 0 | | |
| Error | | | 0 | | | | | |

D

| Midpoint of sleep rise (long) | Effect | SS | $DF_n$ | MS | $DF_d$ | MS error | F | p |
|---|---|---|---|---|---|---|---|---|
| Intercept | Fixed | 1561.343 | 1 | 1561.343 | 3 | 0.132432 | 11789.78 | 0.000 |
| selection | Fixed | 3.091 | 2 | 1.545 | 6 | 0.302752 | 5.1 | 0.051 |
| block | Random | 0.397 | 3 | 0.132 | 6 | 0.302752 | 0.44 | 0.734 |
| selection*block | Random | 1.817 | 6 | 0.303 | 0 | 0 | | |
| Error | | | 0 | | | | | |

E

| Tukey's HSD test- night parameters | | | |
|---|---|---|---|
| | Behavioural night | | Cryophase |
| Mean difference | midpoint of sleep rise | median bout duration | latency |
| ***early vs control*** | -1.0870 | -19.7064 | 4.2971 |
| ***early vs late*** | -1.6890 | -24.1221 | 13.3434 |
| ***control vs late*** | -0.6020 | -4.4157 | 9.0463 |
| Minimum significant difference | 0.6959 | 24.1126 | 12.2353 |
| q critical | 4.339 | 4.339 | 4.339 |
| K, N | 3, 4 | 3,4 | 3, 4 |

**Table S9. Midpoint of rise in sleep under LD12:12:** Midpoint of rise in sleep of virgin male chronotype flies under LD12:12. (A) Mean ± SEM midpoint (Zeitgeber time, h) and number of flies for each replicate population (block). (B) Two-way randomized block ANOVA (selection as a fixed factor and block as a random factor).

A

| Selection | Block | n flies | Midpoint_avg_ZT (h) | SEM_ZT (h) |
|---|---|---|---|---|
| *control* | 1 | 30 | 12.852 | 0.057 |
| *control* | 2 | 31 | 12.655 | 0.088 |
| *control* | 3 | 30 | 12.709 | 0.047 |
| *control* | 4 | 29 | 12.642 | 0.059 |
| *early* | 1 | 29 | 12.694 | 0.061 |
| *early* | 2 | 28 | 12.717 | 0.058 |
| *early* | 3 | 30 | 12.634 | 0.097 |
| *early* | 4 | 31 | 12.733 | 0.038 |
| *late* | 1 | 30 | 12.898 | 0.056 |
| *late* | 2 | 29 | 12.782 | 0.055 |
| *late* | 3 | 30 | 12.748 | 0.043 |
| *late* | 4 | 26 | 12.819 | 0.060 |

**B**

| Midpoint of rise in sleep (LD 12:12) ♂ | Effect | SS | $DF_n$ | MS | $DF_d$ | MS error | F | p |
|---|---|---|---|---|---|---|---|---|
| Intercept | Fixed | 1947.774 | 1 | 1947.774 | 3 | 0.008008 | 243242.2 | 0 |
| selection | Fixed | 0.031 | 2 | 0.016 | 6 | 0.00368 | 4.3 | **0.070021** |
| block | Random | 0.024 | 3 | 0.008 | 6 | 0.00368 | 2.2 | 0.191853 |
| selection*block | Random | 0.022 | 6 | 0.004 | 0 | 0 | | |
| Error | | | 0 | | | | | |

**Table S10. Phase advance:** Linear mixed-effects model (selection as a fixed factor, day as a repeated-measures factor and block as a random factor) for change from baseline in sleep parameters of male chronotype flies after a 9 h phase advance. (A) Type III ANOVA with Satterthwaite df. (B) Tukey's HSD post hoc comparisons for significant terms.

A

| Regime | Parameter | Term | $DF_n$ | $DF_d$ | SS | MS | F | p |
|---|---|---|---|---|---|---|---|---|
| Advance | Day.duration | Selection | 2 | 6 | 1470.2479 | 735.1239 | 3.0576 | 0.1215 |
| Advance | Day.duration | Day | 5 | 45 | 83441.1759 | 16688.2352 | 69.4104 | 0 |
| Advance | Day.duration | Selection:Day | 10 | 45 | 7356.4612 | 735.6461 | 3.0597 | 0.0048 |
| Advance | Day.BoutNumber | Selection | 2 | 51 | 2.0738 | 1.0369 | 1.2845 | 0.2856 |
| Advance | Day.BoutNumber | Day | 5 | 51 | 37.5327 | 7.5065 | 9.2989 | 0 |
| Advance | Day.BoutNumber | Selection:Day | 10 | 51 | 10.5653 | 1.0565 | 1.3088 | 0.2512 |
| Advance | Day.BoutDuration.Median | Selection | 2 | 9 | 2.2643 | 1.1321 | 0.4149 | 0.6724 |
| Advance | Day.BoutDuration.Median | Day | 5 | 45 | 146.8878 | 29.3776 | 10.7653 | 0 |
| Advance | Day.BoutDuration.Median | Selection:Day | 10 | 45 | 12.9916 | 1.2992 | 0.4761 | 0.8967 |
| Advance | Day.Latency | Selection | 2 | 51 | 1502.9058 | 751.4529 | 3.1052 | 0.0534 |
| Advance | Day.Latency | Day | 5 | 51 | 6919.0091 | 1383.8018 | 5.7182 | 0.0003 |
| Advance | Day.Latency | Selection:Day | 10 | 51 | 4986.2541 | 498.6254 | 2.0604 | 0.0456 |
| Advance | Night.duration | Selection | 2 | 51 | 2012.1884 | 1006.0942 | 15.1182 | 0 |
| Advance | Night.duration | Day | 5 | 51 | 14608.334 | 2921.6668 | 43.9029 | 0 |
| Advance | Night.duration | Selection:Day | 10 | 51 | 3142.3423 | 314.2342 | 4.7219 | 0.0001 |

| Regime | Parameter | Term | $DF_n$ | $DF_d$ | SS | MS | F | p |
|---|---|---|---|---|---|---|---|---|
| Advance | Night.BoutNumber | Selection | 2 | 9 | 3.3043 | 1.6522 | 1.8829 | 0.2074 |
| Advance | Night.BoutNumber | Day | 5 | 45 | 50.9849 | 10.197 | 11.6213 | 0 |
| Advance | Night.BoutNumber | Selection:Day | 10 | 45 | 16.7489 | 1.6749 | 1.9088 | 0.0689 |
| Advance | Night.BoutDuration.Median | Selection | 2 | 6 | 428.9807 | 214.4904 | 1.9612 | 0.2211 |
| Advance | Night.BoutDuration.Median | Day | 5 | 45 | 2243.0405 | 448.6081 | 4.1019 | 0.0037 |
| Advance | Night.BoutDuration.Median | Selection:Day | 10 | 45 | 1511.6175 | 151.1617 | 1.3822 | 0.2193 |
| Advance | Night.Latency | Selection | 2 | 9 | 0.4014 | 0.2007 | 0.2089 | 0.8153 |
| Advance | Night.Latency | Day | 5 | 45 | 299.0832 | 59.8166 | 62.2516 | 0 |
| Advance | Night.Latency | Selection:Day | 10 | 45 | 9.5169 | 0.9517 | 0.9904 | 0.4656 |

B

| Parameter | Comparison | Within | Contrast (A vs B) | Estimate (A − B) | SE | df | t ratio | p (Tukey) | Significance |
|---|---|---|---|---|---|---|---|---|---|
| Daytime sleep duration | Lines within each day | Day −3 | early vs control | 0.58 | 11.69 | 42.0 | 0.05 | 0.9987 | ns |
| Daytime sleep duration | Lines within each day | Day −3 | early vs late | 2.88 | 11.69 | 42.0 | 0.25 | 0.9671 | ns |
| Daytime sleep duration | Lines within each day | Day −3 | control vs late | 2.30 | 11.69 | 42.0 | 0.20 | 0.9788 | ns |
| Daytime sleep duration | Lines within each day | Day −2 | early vs control | 4.27 | 11.69 | 42.0 | 0.37 | 0.9293 | ns |
| Daytime sleep duration | Lines within each day | Day −2 | early vs late | 2.26 | 11.69 | 42.0 | 0.19 | 0.9795 | ns |
| Daytime sleep duration | Lines within each day | Day −2 | control vs late | -2.00 | 11.69 | 42.0 | -0.17 | 0.9839 | ns |
| Daytime sleep duration | Lines within each day | Day −1 | early vs control | -4.93 | 11.69 | 42.0 | -0.42 | 0.9068 | ns |
| Daytime sleep duration | Lines within each day | Day −1 | early vs late | -5.14 | 11.69 | 42.0 | -0.44 | 0.8990 | ns |
| Daytime sleep duration | Lines within each day | Day −1 | control vs late | -0.21 | 11.69 | 42.0 | -0.02 | 0.9998 | ns |
| Daytime sleep duration | Lines within each day | Day 1 | early vs control | 56.02 | 11.69 | 42.0 | 4.79 | **0.0001** | *** |
| Daytime sleep duration | Lines within each day | Day 1 | early vs late | 56.02 | 11.69 | 42.0 | 4.79 | **0.0001** | *** |
| Daytime sleep duration | Lines within each day | Day 1 | control vs late | 0.01 | 11.69 | 42.0 | 0.00 | 1.0000 | ns |
| Daytime sleep duration | Lines within each day | Day 2 | early vs control | 17.23 | 11.69 | 42.0 | 1.47 | 0.3132 | ns |
| Daytime sleep duration | Lines within each day | Day 2 | early vs late | 25.15 | 11.69 | 42.0 | 2.15 | 0.0917 | ns |
| Daytime sleep duration | Lines within each day | Day 2 | control vs late | 7.91 | 11.69 | 42.0 | 0.68 | 0.7780 | ns |
| Daytime sleep duration | Lines within each day | Day 3 | early vs control | -6.39 | 11.69 | 42.0 | -0.55 | 0.8487 | ns |
| Daytime sleep duration | Lines within each day | Day 3 | early vs late | 3.87 | 11.69 | 42.0 | 0.33 | 0.9415 | ns |
| Daytime sleep duration | Lines within each day | Day 3 | control vs late | 10.26 | 11.69 | 42.0 | 0.88 | 0.6573 | ns |
| Daytime sleep duration | Days within each line | early | Day −3 vs Day −2 | 1.47 | 10.96 | 45.0 | 0.13 | 1.0000 | ns |
| Daytime sleep duration | Days within each line | early | Day −3 vs Day −1 | 18.57 | 10.96 | 45.0 | 1.69 | 0.5428 | ns |
| Daytime sleep duration | Days within each line | early | Day −3 vs Day 1 | -116.51 | 10.96 | 45.0 | -10.63 | **<0.0001** | *** |
| Daytime sleep duration | Days within each line | early | Day −3 vs Day 2 | 8.44 | 10.96 | 45.0 | 0.77 | 0.9712 | ns |
| Daytime sleep duration | Days within each line | early | Day −3 vs Day 3 | 8.97 | 10.96 | 45.0 | 0.82 | 0.9627 | ns |
| Daytime sleep duration | Days within each line | early | Day −2 vs Day −1 | 17.11 | 10.96 | 45.0 | 1.56 | 0.6282 | ns |
| Daytime sleep duration | Days within each line | early | Day −2 vs Day 1 | -117.98 | 10.96 | 45.0 | -10.76 | **<0.0001** | *** |
| Daytime sleep duration | Days within each line | early | Day −2 vs Day 2 | 6.97 | 10.96 | 45.0 | 0.64 | 0.9876 | ns |
| Daytime sleep duration | Days within each line | early | Day −2 vs Day 3 | 7.50 | 10.96 | 45.0 | 0.68 | 0.9828 | ns |
| Daytime sleep duration | Days within each line | early | Day −1 vs Day 1 | -135.08 | 10.96 | 45.0 | -12.32 | **<0.0001** | *** |
| Daytime sleep duration | Days within each line | early | Day −1 vs Day 2 | -10.13 | 10.96 | 45.0 | -0.92 | 0.9382 | ns |
| Daytime sleep duration | Days within each line | early | Day −1 vs Day 3 | -9.60 | 10.96 | 45.0 | -0.88 | 0.9503 | ns |
| Daytime sleep duration | Days within each line | early | Day 1 vs Day 2 | 124.95 | 10.96 | 45.0 | 11.40 | **<0.0001** | *** |
| Daytime sleep duration | Days within each line | early | Day 1 vs Day 3 | 125.48 | 10.96 | 45.0 | 11.44 | **<0.0001** | *** |
| Daytime sleep duration | Days within each line | early | Day 2 vs Day 3 | 0.53 | 10.96 | 45.0 | 0.05 | 1.0000 | ns |
| Daytime sleep duration | Days within each line | control | Day −3 vs Day −2 | 5.16 | 10.96 | 45.0 | 0.47 | 0.9969 | ns |
| Daytime sleep duration | Days within each line | control | Day −3 vs Day −1 | 13.07 | 10.96 | 45.0 | 1.19 | 0.8384 | ns |
| Daytime sleep duration | Days within each line | control | Day −3 vs Day 1 | -61.07 | 10.96 | 45.0 | -5.57 | **<0.0001** | *** |
| Daytime sleep duration | Days within each line | control | Day −3 vs Day 2 | 25.10 | 10.96 | 45.0 | 2.29 | 0.2197 | ns |
| Daytime sleep duration | Days within each line | control | Day −3 vs Day 3 | 2.00 | 10.96 | 45.0 | 0.18 | 1.0000 | ns |
| Daytime sleep duration | Days within each line | control | Day −2 vs Day −1 | 7.91 | 10.96 | 45.0 | 0.72 | 0.9783 | ns |
| Daytime sleep duration | Days within each line | control | Day −2 vs Day 1 | -66.23 | 10.96 | 45.0 | -6.04 | **<0.0001** | *** |
| Daytime sleep duration | Days within each line | control | Day −2 vs Day 2 | 19.94 | 10.96 | 45.0 | 1.82 | 0.4645 | ns |
| Daytime sleep duration | Days within each line | control | Day −2 vs Day 3 | -3.16 | 10.96 | 45.0 | -0.29 | 0.9997 | ns |
| Daytime sleep duration | Days within each line | control | Day −1 vs Day 1 | -74.14 | 10.96 | 45.0 | -6.76 | **<0.0001** | *** |
| Daytime sleep duration | Days within each line | control | Day −1 vs Day 2 | 12.03 | 10.96 | 45.0 | 1.10 | 0.8798 | ns |

| Parameter | Comparison | Within | Contrast (A vs B) | Estimate (A – B) | SE | df | t ratio | p (Tukey) | Significance |
|---|---|---|---|---|---|---|---|---|---|
| Daytime sleep duration | Days within each line | control | Day −1 vs Day 3 | -11.06 | 10.96 | 45.0 | -1.01 | 0.9125 | ns |
| Daytime sleep duration | Days within each line | control | Day 1 vs Day 2 | 86.17 | 10.96 | 45.0 | 7.86 | **<0.0001** | *** |
| Daytime sleep duration | Days within each line | control | Day 1 vs Day 3 | 63.07 | 10.96 | 45.0 | 5.75 | **<0.0001** | *** |
| Daytime sleep duration | Days within each line | control | Day 2 vs Day 3 | -23.10 | 10.96 | 45.0 | -2.11 | 0.3024 | ns |
| Daytime sleep duration | Days within each line | late | Day −3 vs Day −2 | 0.85 | 10.96 | 45.0 | 0.08 | 1.0000 | ns |
| Daytime sleep duration | Days within each line | late | Day −3 vs Day −1 | 10.55 | 10.96 | 45.0 | 0.96 | 0.9273 | ns |
| Daytime sleep duration | Days within each line | late | Day −3 vs Day 1 | -63.37 | 10.96 | 45.0 | -5.78 | **<0.0001** | *** |
| Daytime sleep duration | Days within each line | late | Day −3 vs Day 2 | 30.71 | 10.96 | 45.0 | 2.80 | 0.0757 | ns |
| Daytime sleep duration | Days within each line | late | Day −3 vs Day 3 | 9.96 | 10.96 | 45.0 | 0.91 | 0.9424 | ns |
| Daytime sleep duration | Days within each line | late | Day −2 vs Day −1 | 9.70 | 10.96 | 45.0 | 0.88 | 0.9483 | ns |
| Daytime sleep duration | Days within each line | late | Day −2 vs Day 1 | -64.22 | 10.96 | 45.0 | -5.86 | **<0.0001** | *** |
| Daytime sleep duration | Days within each line | late | Day −2 vs Day 2 | 29.86 | 10.96 | 45.0 | 2.72 | 0.0903 | ns |
| Daytime sleep duration | Days within each line | late | Day −2 vs Day 3 | 9.10 | 10.96 | 45.0 | 0.83 | 0.9603 | ns |
| Daytime sleep duration | Days within each line | late | Day −1 vs Day 1 | -73.92 | 10.96 | 45.0 | -6.74 | **<0.0001** | *** |
| Daytime sleep duration | Days within each line | late | Day −1 vs Day 2 | 20.16 | 10.96 | 45.0 | 1.84 | 0.4523 | ns |
| Daytime sleep duration | Days within each line | late | Day −1 vs Day 3 | -0.59 | 10.96 | 45.0 | -0.05 | 1.0000 | ns |
| Daytime sleep duration | Days within each line | late | Day 1 vs Day 2 | 94.08 | 10.96 | 45.0 | 8.58 | **<0.0001** | *** |
| Daytime sleep duration | Days within each line | late | Day 1 vs Day 3 | 73.32 | 10.96 | 45.0 | 6.69 | **<0.0001** | *** |
| Daytime sleep duration | Days within each line | late | Day 2 vs Day 3 | -20.75 | 10.96 | 45.0 | -1.89 | 0.4196 | ns |
| Daytime sleep latency | Lines within each day | Day −3 | early vs control | -1.38 | 11.00 | 51.0 | -0.13 | 0.9914 | ns |
| Daytime sleep latency | Lines within each day | Day −3 | early vs late | -3.38 | 11.00 | 51.0 | -0.31 | 0.9493 | ns |
| Daytime sleep latency | Lines within each day | Day −3 | control vs late | -2.01 | 11.00 | 51.0 | -0.18 | 0.9818 | ns |
| Daytime sleep latency | Lines within each day | Day −2 | early vs control | 2.07 | 11.00 | 51.0 | 0.19 | 0.9807 | ns |
| Daytime sleep latency | Lines within each day | Day −2 | early vs late | 4.28 | 11.00 | 51.0 | 0.39 | 0.9202 | ns |
| Daytime sleep latency | Lines within each day | Day −2 | control vs late | 2.21 | 11.00 | 51.0 | 0.20 | 0.9781 | ns |
| Daytime sleep latency | Lines within each day | Day −1 | early vs control | -0.15 | 11.00 | 51.0 | -0.01 | 0.9999 | ns |
| Daytime sleep latency | Lines within each day | Day −1 | early vs late | -0.69 | 11.00 | 51.0 | -0.06 | 0.9978 | ns |
| Daytime sleep latency | Lines within each day | Day −1 | control vs late | -0.55 | 11.00 | 51.0 | -0.05 | 0.9986 | ns |
| Daytime sleep latency | Lines within each day | Day 1 | early vs control | -27.80 | 11.00 | 51.0 | -2.53 | **0.0383** | * |
| Daytime sleep latency | Lines within each day | Day 1 | early vs late | -53.36 | 11.00 | 51.0 | -4.85 | **<0.0001** | *** |
| Daytime sleep latency | Lines within each day | Day 1 | control vs late | -25.57 | 11.00 | 51.0 | -2.32 | 0.0614 | ns |
| Daytime sleep latency | Lines within each day | Day 2 | early vs control | 7.98 | 11.00 | 51.0 | 0.73 | 0.7494 | ns |
| Daytime sleep latency | Lines within each day | Day 2 | early vs late | -1.22 | 11.00 | 51.0 | -0.11 | 0.9933 | ns |
| Daytime sleep latency | Lines within each day | Day 2 | control vs late | -9.20 | 11.00 | 51.0 | -0.84 | 0.6822 | ns |
| Daytime sleep latency | Lines within each day | Day 3 | early vs control | 7.52 | 11.00 | 51.0 | 0.68 | 0.7742 | ns |
| Daytime sleep latency | Lines within each day | Day 3 | early vs late | -8.75 | 11.00 | 51.0 | -0.80 | 0.7076 | ns |
| Daytime sleep latency | Lines within each day | Day 3 | control vs late | -16.26 | 11.00 | 51.0 | -1.48 | 0.3096 | ns |
| Daytime sleep latency | Days within each line | early | Day −3 vs Day −2 | -11.08 | 11.00 | 51.0 | -1.01 | 0.9134 | ns |
| Daytime sleep latency | Days within each line | early | Day −3 vs Day −1 | -18.30 | 11.00 | 51.0 | -1.66 | 0.5617 | ns |
| Daytime sleep latency | Days within each line | early | Day −3 vs Day 1 | 16.40 | 11.00 | 51.0 | 1.49 | 0.6713 | ns |
| Daytime sleep latency | Days within each line | early | Day −3 vs Day 2 | -32.66 | 11.00 | 51.0 | -2.97 | **0.0490** | * |
| Daytime sleep latency | Days within each line | early | Day −3 vs Day 3 | -24.35 | 11.00 | 51.0 | -2.21 | 0.2495 | ns |
| Daytime sleep latency | Days within each line | early | Day −2 vs Day −1 | -7.22 | 11.00 | 51.0 | -0.66 | 0.9859 | ns |
| Daytime sleep latency | Days within each line | early | Day −2 vs Day 1 | 27.48 | 11.00 | 51.0 | 2.50 | 0.1438 | ns |
| Daytime sleep latency | Days within each line | early | Day −2 vs Day 2 | -21.58 | 11.00 | 51.0 | -1.96 | 0.3781 | ns |
| Daytime sleep latency | Days within each line | early | Day −2 vs Day 3 | -13.27 | 11.00 | 51.0 | -1.21 | 0.8317 | ns |
| Daytime sleep latency | Days within each line | early | Day −1 vs Day 1 | 34.70 | 11.00 | 51.0 | 3.15 | **0.0305** | * |
| Daytime sleep latency | Days within each line | early | Day −1 vs Day 2 | -14.36 | 11.00 | 51.0 | -1.31 | 0.7805 | ns |
| Daytime sleep latency | Days within each line | early | Day −1 vs Day 3 | -6.06 | 11.00 | 51.0 | -0.55 | 0.9937 | ns |
| Daytime sleep latency | Days within each line | early | Day 1 vs Day 2 | -49.06 | 11.00 | 51.0 | -4.46 | **0.0006** | *** |
| Daytime sleep latency | Days within each line | early | Day 1 vs Day 3 | -40.75 | 11.00 | 51.0 | -3.71 | **0.0065** | ** |
| Daytime sleep latency | Days within each line | early | Day 2 vs Day 3 | 8.31 | 11.00 | 51.0 | 0.76 | 0.9736 | ns |
| Daytime sleep latency | Days within each line | control | Day −3 vs Day −2 | -7.63 | 11.00 | 51.0 | -0.69 | 0.9818 | ns |
| Daytime sleep latency | Days within each line | control | Day −3 vs Day −1 | -17.07 | 11.00 | 51.0 | -1.55 | 0.6332 | ns |
| Daytime sleep latency | Days within each line | control | Day −3 vs Day 1 | -10.02 | 11.00 | 51.0 | -0.91 | 0.9421 | ns |

| Parameter | Comparison | Within | Contrast (A vs B) | Estimate (A − B) | SE | df | t ratio | p (Tukey) | Significance |
|---|---|---|---|---|---|---|---|---|---|
| Daytime sleep latency | Days within each line | control | Day −3 vs Day 2 | -23.30 | 11.00 | 51.0 | -2.12 | 0.2946 | ns |
| Daytime sleep latency | Days within each line | control | Day −3 vs Day 3 | -15.46 | 11.00 | 51.0 | -1.41 | 0.7237 | ns |
| Daytime sleep latency | Days within each line | control | Day −2 vs Day −1 | -9.43 | 11.00 | 51.0 | -0.86 | 0.9547 | ns |
| Daytime sleep latency | Days within each line | control | Day −2 vs Day 1 | -2.38 | 11.00 | 51.0 | -0.22 | 0.9999 | ns |
| Daytime sleep latency | Days within each line | control | Day −2 vs Day 2 | -15.66 | 11.00 | 51.0 | -1.42 | 0.7125 | ns |
| Daytime sleep latency | Days within each line | control | Day −2 vs Day 3 | -7.83 | 11.00 | 51.0 | -0.71 | 0.9797 | ns |
| Daytime sleep latency | Days within each line | control | Day −1 vs Day 1 | 7.05 | 11.00 | 51.0 | 0.64 | 0.9872 | ns |
| Daytime sleep latency | Days within each line | control | Day −1 vs Day 2 | -6.23 | 11.00 | 51.0 | -0.57 | 0.9928 | ns |
| Daytime sleep latency | Days within each line | control | Day −1 vs Day 3 | 1.61 | 11.00 | 51.0 | 0.15 | 1.0000 | ns |
| Daytime sleep latency | Days within each line | control | Day 1 vs Day 2 | -13.28 | 11.00 | 51.0 | -1.21 | 0.8312 | ns |
| Daytime sleep latency | Days within each line | control | Day 1 vs Day 3 | -5.44 | 11.00 | 51.0 | -0.49 | 0.9961 | ns |
| Daytime sleep latency | Days within each line | control | Day 2 vs Day 3 | 7.84 | 11.00 | 51.0 | 0.71 | 0.9796 | ns |
| Daytime sleep latency | Days within each line | late | Day −3 vs Day −2 | -3.42 | 11.00 | 51.0 | -0.31 | 0.9996 | ns |
| Daytime sleep latency | Days within each line | late | Day −3 vs Day −1 | -15.61 | 11.00 | 51.0 | -1.42 | 0.7156 | ns |
| Daytime sleep latency | Days within each line | late | Day −3 vs Day 1 | -33.58 | 11.00 | 51.0 | -3.05 | **0.0397** | * |
| Daytime sleep latency | Days within each line | late | Day −3 vs Day 2 | -30.50 | 11.00 | 51.0 | -2.77 | 0.0786 | ns |
| Daytime sleep latency | Days within each line | late | Day −3 vs Day 3 | -29.72 | 11.00 | 51.0 | -2.70 | 0.0925 | ns |
| Daytime sleep latency | Days within each line | late | Day −2 vs Day −1 | -12.19 | 11.00 | 51.0 | -1.11 | 0.8759 | ns |
| Daytime sleep latency | Days within each line | late | Day −2 vs Day 1 | -30.16 | 11.00 | 51.0 | -2.74 | 0.0844 | ns |
| Daytime sleep latency | Days within each line | late | Day −2 vs Day 2 | -27.07 | 11.00 | 51.0 | -2.46 | 0.1552 | ns |
| Daytime sleep latency | Days within each line | late | Day −2 vs Day 3 | -26.30 | 11.00 | 51.0 | -2.39 | 0.1789 | ns |
| Daytime sleep latency | Days within each line | late | Day −1 vs Day 1 | -17.97 | 11.00 | 51.0 | -1.63 | 0.5808 | ns |
| Daytime sleep latency | Days within each line | late | Day −1 vs Day 2 | -14.89 | 11.00 | 51.0 | -1.35 | 0.7539 | ns |
| Daytime sleep latency | Days within each line | late | Day −1 vs Day 3 | -14.11 | 11.00 | 51.0 | -1.28 | 0.7930 | ns |
| Daytime sleep latency | Days within each line | late | Day 1 vs Day 2 | 3.08 | 11.00 | 51.0 | 0.28 | 0.9998 | ns |
| Daytime sleep latency | Days within each line | late | Day 1 vs Day 3 | 3.86 | 11.00 | 51.0 | 0.35 | 0.9993 | ns |
| Daytime sleep latency | Days within each line | late | Day 2 vs Day 3 | 0.78 | 11.00 | 51.0 | 0.07 | 1.0000 | ns |
| Daytime bout number | Days (lines pooled) | – | Day −3 vs Day −2 | -0.08 | 0.37 | 51.0 | -0.22 | 0.9999 | ns |
| Daytime bout number | Days (lines pooled) | – | Day −3 vs Day −1 | -0.12 | 0.37 | 51.0 | -0.33 | 0.9995 | ns |
| Daytime bout number | Days (lines pooled) | – | Day −3 vs Day 1 | -1.58 | 0.37 | 51.0 | -4.30 | **0.0011** | ** |
| Daytime bout number | Days (lines pooled) | – | Day −3 vs Day 2 | -1.08 | 0.37 | 51.0 | -2.95 | 0.0515 | ns |
| Daytime bout number | Days (lines pooled) | – | Day −3 vs Day 3 | -1.71 | 0.37 | 51.0 | -4.67 | **0.0003** | *** |
| Daytime bout number | Days (lines pooled) | – | Day −2 vs Day −1 | -0.04 | 0.37 | 51.0 | -0.11 | 1.0000 | ns |
| Daytime bout number | Days (lines pooled) | – | Day −2 vs Day 1 | -1.49 | 0.37 | 51.0 | -4.07 | **0.0021** | ** |
| Daytime bout number | Days (lines pooled) | – | Day −2 vs Day 2 | -1.00 | 0.37 | 51.0 | -2.73 | 0.0870 | ns |
| Daytime bout number | Days (lines pooled) | – | Day −2 vs Day 3 | -1.63 | 0.37 | 51.0 | -4.45 | **0.0006** | *** |
| Daytime bout number | Days (lines pooled) | – | Day −1 vs Day 1 | -1.46 | 0.37 | 51.0 | -3.97 | **0.0030** | ** |
| Daytime bout number | Days (lines pooled) | – | Day −1 vs Day 2 | -0.96 | 0.37 | 51.0 | -2.62 | 0.1104 | ns |
| Daytime bout number | Days (lines pooled) | – | Day −1 vs Day 3 | -1.59 | 0.37 | 51.0 | -4.35 | **0.0009** | *** |
| Daytime bout number | Days (lines pooled) | – | Day 1 vs Day 2 | 0.49 | 0.37 | 51.0 | 1.35 | 0.7580 | ns |
| Daytime bout number | Days (lines pooled) | – | Day 1 vs Day 3 | -0.14 | 0.37 | 51.0 | -0.38 | 0.9989 | ns |
| Daytime bout number | Days (lines pooled) | – | Day 2 vs Day 3 | -0.63 | 0.37 | 51.0 | -1.72 | 0.5224 | ns |
| Daytime median bout duration | Days (lines pooled) | – | Day −3 vs Day −2 | -0.06 | 0.67 | 45.0 | -0.09 | 1.0000 | ns |
| Daytime median bout duration | Days (lines pooled) | – | Day −3 vs Day −1 | 1.12 | 0.67 | 45.0 | 1.66 | 0.5663 | ns |
| Daytime median bout duration | Days (lines pooled) | – | Day −3 vs Day 1 | 0.02 | 0.67 | 45.0 | 0.03 | 1.0000 | ns |
| Daytime median bout duration | Days (lines pooled) | – | Day −3 vs Day 2 | 2.97 | 0.67 | 45.0 | 4.41 | **0.0009** | *** |
| Daytime median bout duration | Days (lines pooled) | – | Day −3 vs Day 3 | 3.36 | 0.67 | 45.0 | 4.98 | **0.0001** | *** |
| Daytime median bout duration | Days (lines pooled) | – | Day −2 vs Day −1 | 1.18 | 0.67 | 45.0 | 1.75 | 0.5062 | ns |
| Daytime median bout duration | Days (lines pooled) | – | Day −2 vs Day 1 | 0.08 | 0.67 | 45.0 | 0.13 | 1.0000 | ns |
| Daytime median bout duration | Days (lines pooled) | – | Day −2 vs Day 2 | 3.04 | 0.67 | 45.0 | 4.50 | **0.0006** | *** |
| Daytime median bout duration | Days (lines pooled) | – | Day −2 vs Day 3 | 3.42 | 0.67 | 45.0 | 5.08 | **0.0001** | *** |
| Daytime median bout duration | Days (lines pooled) | – | Day −1 vs Day 1 | -1.10 | 0.67 | 45.0 | -1.63 | 0.5862 | ns |
| Daytime median bout duration | Days (lines pooled) | – | Day −1 vs Day 2 | 1.86 | 0.67 | 45.0 | 2.75 | 0.0848 | ns |
| Daytime median bout duration | Days (lines pooled) | – | Day −1 vs Day 3 | 2.24 | 0.67 | 45.0 | 3.33 | **0.0204** | * |
| Daytime median bout duration | Days (lines pooled) | – | Day 1 vs Day 2 | 2.95 | 0.67 | 45.0 | 4.38 | **0.0009** | *** |

| Parameter | Comparison | Within | Contrast (A vs B) | Estimate (A – B) | SE | df | t ratio | p (Tukey) | Significance |
|---|---|---|---|---|---|---|---|---|---|
| Daytime median bout duration | Days (lines pooled) | – | Day 1 vs Day 3 | 3.34 | 0.67 | 45.0 | 4.95 | **0.0001** | *** |
| Daytime median bout duration | Days (lines pooled) | – | Day 2 vs Day 3 | 0.39 | 0.67 | 45.0 | 0.58 | 0.9922 | ns |
| Nighttime sleep duration | Lines within each day | Day −3 | early vs control | -11.50 | 5.77 | 51.0 | -1.99 | 0.1243 | ns |
| Nighttime sleep duration | Lines within each day | Day −3 | early vs late | -7.95 | 5.77 | 51.0 | -1.38 | 0.3595 | ns |
| Nighttime sleep duration | Lines within each day | Day −3 | control vs late | 3.54 | 5.77 | 51.0 | 0.61 | 0.8130 | ns |
| Nighttime sleep duration | Lines within each day | Day −2 | early vs control | 6.60 | 5.77 | 51.0 | 1.14 | 0.4918 | ns |
| Nighttime sleep duration | Lines within each day | Day −2 | early vs late | 0.80 | 5.77 | 51.0 | 0.14 | 0.9894 | ns |
| Nighttime sleep duration | Lines within each day | Day −2 | control vs late | -5.80 | 5.77 | 51.0 | -1.01 | 0.5771 | ns |
| Nighttime sleep duration | Lines within each day | Day −1 | early vs control | 4.69 | 5.77 | 51.0 | 0.81 | 0.6971 | ns |
| Nighttime sleep duration | Lines within each day | Day −1 | early vs late | 6.98 | 5.77 | 51.0 | 1.21 | 0.4526 | ns |
| Nighttime sleep duration | Lines within each day | Day −1 | control vs late | 2.29 | 5.77 | 51.0 | 0.40 | 0.9166 | ns |
| Nighttime sleep duration | Lines within each day | Day 1 | early vs control | 12.72 | 5.77 | 51.0 | 2.21 | 0.0798 | ns |
| Nighttime sleep duration | Lines within each day | Day 1 | early vs late | 36.07 | 5.77 | 51.0 | 6.25 | **<0.0001** | *** |
| Nighttime sleep duration | Lines within each day | Day 1 | control vs late | 23.35 | 5.77 | 51.0 | 4.05 | **0.0005** | *** |
| Nighttime sleep duration | Lines within each day | Day 2 | early vs control | -2.62 | 5.77 | 51.0 | -0.45 | 0.8930 | ns |
| Nighttime sleep duration | Lines within each day | Day 2 | early vs late | 11.70 | 5.77 | 51.0 | 2.03 | 0.1156 | ns |
| Nighttime sleep duration | Lines within each day | Day 2 | control vs late | 14.32 | 5.77 | 51.0 | 2.48 | **0.0426** | * |
| Nighttime sleep duration | Lines within each day | Day 3 | early vs control | 9.42 | 5.77 | 51.0 | 1.63 | 0.2416 | ns |
| Nighttime sleep duration | Lines within each day | Day 3 | early vs late | 27.22 | 5.77 | 51.0 | 4.72 | **0.0001** | *** |
| Nighttime sleep duration | Lines within each day | Day 3 | control vs late | 17.81 | 5.77 | 51.0 | 3.09 | **0.0090** | ** |
| Nighttime sleep duration | Days within each line | early | Day −3 vs Day −2 | -15.67 | 5.77 | 51.0 | -2.72 | 0.0894 | ns |
| Nighttime sleep duration | Days within each line | early | Day −3 vs Day −1 | -21.21 | 5.77 | 51.0 | -3.68 | **0.0071** | ** |
| Nighttime sleep duration | Days within each line | early | Day −3 vs Day 1 | -26.95 | 5.77 | 51.0 | -4.67 | **0.0003** | *** |
| Nighttime sleep duration | Days within each line | early | Day −3 vs Day 2 | -45.17 | 5.77 | 51.0 | -7.83 | **<0.0001** | *** |
| Nighttime sleep duration | Days within each line | early | Day −3 vs Day 3 | -52.76 | 5.77 | 51.0 | -9.15 | **<0.0001** | *** |
| Nighttime sleep duration | Days within each line | early | Day −2 vs Day −1 | -5.55 | 5.77 | 51.0 | -0.96 | 0.9279 | ns |
| Nighttime sleep duration | Days within each line | early | Day −2 vs Day 1 | -11.28 | 5.77 | 51.0 | -1.96 | 0.3819 | ns |
| Nighttime sleep duration | Days within each line | early | Day −2 vs Day 2 | -29.50 | 5.77 | 51.0 | -5.11 | **0.0001** | *** |
| Nighttime sleep duration | Days within each line | early | Day −2 vs Day 3 | -37.09 | 5.77 | 51.0 | -6.43 | **<0.0001** | *** |
| Nighttime sleep duration | Days within each line | early | Day −1 vs Day 1 | -5.73 | 5.77 | 51.0 | -0.99 | 0.9179 | ns |
| Nighttime sleep duration | Days within each line | early | Day −1 vs Day 2 | -23.95 | 5.77 | 51.0 | -4.15 | **0.0017** | ** |
| Nighttime sleep duration | Days within each line | early | Day −1 vs Day 3 | -31.54 | 5.77 | 51.0 | -5.47 | **<0.0001** | *** |
| Nighttime sleep duration | Days within each line | early | Day 1 vs Day 2 | -18.22 | 5.77 | 51.0 | -3.16 | **0.0302** | * |
| Nighttime sleep duration | Days within each line | early | Day 1 vs Day 3 | -25.81 | 5.77 | 51.0 | -4.47 | **0.0006** | *** |
| Nighttime sleep duration | Days within each line | early | Day 2 vs Day 3 | -7.59 | 5.77 | 51.0 | -1.32 | 0.7750 | ns |
| Nighttime sleep duration | Days within each line | control | Day −3 vs Day −2 | 2.43 | 5.77 | 51.0 | 0.42 | 0.9982 | ns |
| Nighttime sleep duration | Days within each line | control | Day −3 vs Day −1 | -5.03 | 5.77 | 51.0 | -0.87 | 0.9514 | ns |
| Nighttime sleep duration | Days within each line | control | Day −3 vs Day 1 | -2.73 | 5.77 | 51.0 | -0.47 | 0.9969 | ns |
| Nighttime sleep duration | Days within each line | control | Day −3 vs Day 2 | -36.29 | 5.77 | 51.0 | -6.29 | **<0.0001** | *** |
| Nighttime sleep duration | Days within each line | control | Day −3 vs Day 3 | -31.85 | 5.77 | 51.0 | -5.52 | **<0.0001** | *** |
| Nighttime sleep duration | Days within each line | control | Day −2 vs Day −1 | -7.46 | 5.77 | 51.0 | -1.29 | 0.7874 | ns |
| Nighttime sleep duration | Days within each line | control | Day −2 vs Day 1 | -5.15 | 5.77 | 51.0 | -0.89 | 0.9464 | ns |
| Nighttime sleep duration | Days within each line | control | Day −2 vs Day 2 | -38.72 | 5.77 | 51.0 | -6.71 | **<0.0001** | *** |
| Nighttime sleep duration | Days within each line | control | Day −2 vs Day 3 | -34.27 | 5.77 | 51.0 | -5.94 | **<0.0001** | *** |
| Nighttime sleep duration | Days within each line | control | Day −1 vs Day 1 | 2.31 | 5.77 | 51.0 | 0.40 | 0.9986 | ns |
| Nighttime sleep duration | Days within each line | control | Day −1 vs Day 2 | -31.26 | 5.77 | 51.0 | -5.42 | **<0.0001** | *** |
| Nighttime sleep duration | Days within each line | control | Day −1 vs Day 3 | -26.82 | 5.77 | 51.0 | -4.65 | **0.0003** | *** |
| Nighttime sleep duration | Days within each line | control | Day 1 vs Day 2 | -33.56 | 5.77 | 51.0 | -5.82 | **<0.0001** | *** |
| Nighttime sleep duration | Days within each line | control | Day 1 vs Day 3 | -29.12 | 5.77 | 51.0 | -5.05 | **0.0001** | *** |
| Nighttime sleep duration | Days within each line | control | Day 2 vs Day 3 | 4.44 | 5.77 | 51.0 | 0.77 | 0.9713 | ns |
| Nighttime sleep duration | Days within each line | late | Day −3 vs Day −2 | -6.91 | 5.77 | 51.0 | -1.20 | 0.8354 | ns |
| Nighttime sleep duration | Days within each line | late | Day −3 vs Day −1 | -6.28 | 5.77 | 51.0 | -1.09 | 0.8836 | ns |
| Nighttime sleep duration | Days within each line | late | Day −3 vs Day 1 | 17.08 | 5.77 | 51.0 | 2.96 | 0.0500 | * |
| Nighttime sleep duration | Days within each line | late | Day −3 vs Day 2 | -25.51 | 5.77 | 51.0 | -4.42 | **0.0007** | *** |
| Nighttime sleep duration | Days within each line | late | Day −3 vs Day 3 | -17.58 | 5.77 | 51.0 | -3.05 | **0.0401** | * |

| Parameter | Comparison | Within | Contrast (A vs B) | Estimate (A – B) | SE | df | t ratio | p (Tukey) | Significance |
|---|---|---|---|---|---|---|---|---|---|
| Nighttime sleep duration | Days within each line | late | Day −2 vs Day −1 | 0.63 | 5.77 | 51.0 | 0.11 | 1.0000 | ns |
| Nighttime sleep duration | Days within each line | late | Day −2 vs Day 1 | 24.00 | 5.77 | 51.0 | 4.16 | **0.0016** | ** |
| Nighttime sleep duration | Days within each line | late | Day −2 vs Day 2 | -18.60 | 5.77 | 51.0 | -3.22 | **0.0254** | * |
| Nighttime sleep duration | Days within each line | late | Day −2 vs Day 3 | -10.67 | 5.77 | 51.0 | -1.85 | 0.4442 | ns |
| Nighttime sleep duration | Days within each line | late | Day −1 vs Day 1 | 23.36 | 5.77 | 51.0 | 4.05 | **0.0023** | ** |
| Nighttime sleep duration | Days within each line | late | Day −1 vs Day 2 | -19.23 | 5.77 | 51.0 | -3.33 | **0.0188** | * |
| Nighttime sleep duration | Days within each line | late | Day −1 vs Day 3 | -11.30 | 5.77 | 51.0 | -1.96 | 0.3793 | ns |
| Nighttime sleep duration | Days within each line | late | Day 1 vs Day 2 | -42.59 | 5.77 | 51.0 | -7.38 | **<0.0001** | *** |
| Nighttime sleep duration | Days within each line | late | Day 1 vs Day 3 | -34.67 | 5.77 | 51.0 | -6.01 | **<0.0001** | *** |
| Nighttime sleep duration | Days within each line | late | Day 2 vs Day 3 | 7.93 | 5.77 | 51.0 | 1.37 | 0.7417 | ns |
| Nighttime sleep latency | Days (lines pooled) | – | Day −3 vs Day −2 | -0.17 | 0.40 | 45.0 | -0.43 | 0.9980 | ns |
| Nighttime sleep latency | Days (lines pooled) | – | Day −3 vs Day −1 | 0.30 | 0.40 | 45.0 | 0.76 | 0.9734 | ns |
| Nighttime sleep latency | Days (lines pooled) | – | Day −3 vs Day 1 | 4.41 | 0.40 | 45.0 | 11.01 | **<0.0001** | *** |
| Nighttime sleep latency | Days (lines pooled) | – | Day −3 vs Day 2 | 4.01 | 0.40 | 45.0 | 10.01 | **<0.0001** | *** |
| Nighttime sleep latency | Days (lines pooled) | – | Day −3 vs Day 3 | 3.88 | 0.40 | 45.0 | 9.70 | **<0.0001** | *** |
| Nighttime sleep latency | Days (lines pooled) | – | Day −2 vs Day −1 | 0.47 | 0.40 | 45.0 | 1.19 | 0.8409 | ns |
| Nighttime sleep latency | Days (lines pooled) | – | Day −2 vs Day 1 | 4.58 | 0.40 | 45.0 | 11.44 | **<0.0001** | *** |
| Nighttime sleep latency | Days (lines pooled) | – | Day −2 vs Day 2 | 4.18 | 0.40 | 45.0 | 10.44 | **<0.0001** | *** |
| Nighttime sleep latency | Days (lines pooled) | – | Day −2 vs Day 3 | 4.05 | 0.40 | 45.0 | 10.13 | **<0.0001** | *** |
| Nighttime sleep latency | Days (lines pooled) | – | Day −1 vs Day 1 | 4.10 | 0.40 | 45.0 | 10.25 | **<0.0001** | *** |
| Nighttime sleep latency | Days (lines pooled) | – | Day −1 vs Day 2 | 3.70 | 0.40 | 45.0 | 9.25 | **<0.0001** | *** |
| Nighttime sleep latency | Days (lines pooled) | – | Day −1 vs Day 3 | 3.58 | 0.40 | 45.0 | 8.95 | **<0.0001** | *** |
| Nighttime sleep latency | Days (lines pooled) | – | Day 1 vs Day 2 | -0.40 | 0.40 | 45.0 | -1.00 | 0.9163 | ns |
| Nighttime sleep latency | Days (lines pooled) | – | Day 1 vs Day 3 | -0.52 | 0.40 | 45.0 | -1.31 | 0.7801 | ns |
| Nighttime sleep latency | Days (lines pooled) | – | Day 2 vs Day 3 | -0.12 | 0.40 | 45.0 | -0.31 | 0.9996 | ns |
| Nighttime bout number | Days (lines pooled) | – | Day −3 vs Day −2 | 0.61 | 0.38 | 45.0 | 1.59 | 0.6084 | ns |
| Nighttime bout number | Days (lines pooled) | – | Day −3 vs Day −1 | 0.63 | 0.38 | 45.0 | 1.65 | 0.5738 | ns |
| Nighttime bout number | Days (lines pooled) | – | Day −3 vs Day 1 | -0.60 | 0.38 | 45.0 | -1.58 | 0.6173 | ns |
| Nighttime bout number | Days (lines pooled) | – | Day −3 vs Day 2 | 2.10 | 0.38 | 45.0 | 5.48 | **<0.0001** | *** |
| Nighttime bout number | Days (lines pooled) | – | Day −3 vs Day 3 | 1.05 | 0.38 | 45.0 | 2.75 | 0.0854 | ns |
| Nighttime bout number | Days (lines pooled) | – | Day −2 vs Day −1 | 0.02 | 0.38 | 45.0 | 0.05 | 1.0000 | ns |
| Nighttime bout number | Days (lines pooled) | – | Day −2 vs Day 1 | -1.21 | 0.38 | 45.0 | -3.17 | **0.0309** | * |
| Nighttime bout number | Days (lines pooled) | – | Day −2 vs Day 2 | 1.49 | 0.38 | 45.0 | 3.89 | **0.0041** | ** |
| Nighttime bout number | Days (lines pooled) | – | Day −2 vs Day 3 | 0.44 | 0.38 | 45.0 | 1.16 | 0.8544 | ns |
| Nighttime bout number | Days (lines pooled) | – | Day −1 vs Day 1 | -1.23 | 0.38 | 45.0 | -3.22 | **0.0268** | * |
| Nighttime bout number | Days (lines pooled) | – | Day −1 vs Day 2 | 1.47 | 0.38 | 45.0 | 3.84 | **0.0048** | ** |
| Nighttime bout number | Days (lines pooled) | – | Day −1 vs Day 3 | 0.42 | 0.38 | 45.0 | 1.10 | 0.8777 | ns |
| Nighttime bout number | Days (lines pooled) | – | Day 1 vs Day 2 | 2.70 | 0.38 | 45.0 | 7.06 | **<0.0001** | *** |
| Nighttime bout number | Days (lines pooled) | – | Day 1 vs Day 3 | 1.65 | 0.38 | 45.0 | 4.33 | **0.0011** | ** |
| Nighttime bout number | Days (lines pooled) | – | Day 2 vs Day 3 | -1.05 | 0.38 | 45.0 | -2.74 | 0.0877 | ns |
| Nighttime median bout duration | Days (lines pooled) | – | Day −3 vs Day −2 | -3.46 | 4.27 | 45.0 | -0.81 | 0.9641 | ns |
| Nighttime median bout duration | Days (lines pooled) | – | Day −3 vs Day −1 | -6.44 | 4.27 | 45.0 | -1.51 | 0.6605 | ns |
| Nighttime median bout duration | Days (lines pooled) | – | Day −3 vs Day 1 | 2.68 | 4.27 | 45.0 | 0.63 | 0.9884 | ns |
| Nighttime median bout duration | Days (lines pooled) | – | Day −3 vs Day 2 | -15.00 | 4.27 | 45.0 | -3.51 | **0.0123** | * |
| Nighttime median bout duration | Days (lines pooled) | – | Day −3 vs Day 3 | -4.70 | 4.27 | 45.0 | -1.10 | 0.8783 | ns |
| Nighttime median bout duration | Days (lines pooled) | – | Day −2 vs Day −1 | -2.98 | 4.27 | 45.0 | -0.70 | 0.9812 | ns |
| Nighttime median bout duration | Days (lines pooled) | – | Day −2 vs Day 1 | 6.14 | 4.27 | 45.0 | 1.44 | 0.7043 | ns |
| Nighttime median bout duration | Days (lines pooled) | – | Day −2 vs Day 2 | -11.54 | 4.27 | 45.0 | -2.70 | 0.0945 | ns |
| Nighttime median bout duration | Days (lines pooled) | – | Day −2 vs Day 3 | -1.24 | 4.27 | 45.0 | -0.29 | 0.9997 | ns |
| Nighttime median bout duration | Days (lines pooled) | – | Day −1 vs Day 1 | 9.12 | 4.27 | 45.0 | 2.14 | 0.2881 | ns |
| Nighttime median bout duration | Days (lines pooled) | – | Day −1 vs Day 2 | -8.56 | 4.27 | 45.0 | -2.00 | 0.3557 | ns |
| Nighttime median bout duration | Days (lines pooled) | – | Day −1 vs Day 3 | 1.74 | 4.27 | 45.0 | 0.41 | 0.9985 | ns |
| Nighttime median bout duration | Days (lines pooled) | – | Day 1 vs Day 2 | -17.68 | 4.27 | 45.0 | -4.14 | **0.0020** | ** |
| Nighttime median bout duration | Days (lines pooled) | – | Day 1 vs Day 3 | -7.38 | 4.27 | 45.0 | -1.73 | 0.5212 | ns |
| Nighttime median bout duration | Days (lines pooled) | – | Day 2 vs Day 3 | 10.30 | 4.27 | 45.0 | 2.41 | 0.1737 | ns |

**Table S11. Phase delay:** Linear mixed-effects model (selection as a fixed factor, day as a repeated-measures factor and block as a random factor) for change from baseline in sleep parameters of male chronotype flies after a 9 h phase delay. (A) Type III ANOVA with Satterthwaite df. (B) Tukey's HSD post hoc comparisons for significant terms.

A

| Regime | Parameter | Term | $DF_n$ | $DF_d$ | SS | MS | F | p |
|---|---|---|---|---|---|---|---|---|
| Delay | Day.Duration | Selection | 2 | 51 | 1095.0537 | 547.5269 | 1.3115 | 0.2784 |
| Delay | Day.Duration | Day | 5 | 51 | 57237.209 | 11447.4418 | 27.4199 | 0 |
| Delay | Day.Duration | Selection:Day | 10 | 51 | 2648.1331 | 264.8133 | 0.6343 | 0.7776 |
| Delay | Day.BoutNumber | Selection | 2 | 6 | 0.0414 | 0.0207 | 0.0121 | 0.988 |
| Delay | Day.BoutNumber | Day | 5 | 45 | 103.8211 | 20.7642 | 12.1814 | 0 |
| Delay | Day.BoutNumber | Selection:Day | 10 | 45 | 12.0794 | 1.2079 | 0.7086 | 0.7115 |
| Delay | Day.BoutDuration.Median | Selection | 2 | 9 | 1.0495 | 0.5247 | 0.4418 | 0.6561 |
| Delay | Day.BoutDuration.Median | Day | 5 | 45 | 143.4663 | 28.6933 | 24.1578 | 0 |
| Delay | Day.BoutDuration.Median | Selection:Day | 10 | 45 | 16.3382 | 1.6338 | 1.3756 | 0.2223 |
| Delay | Day.Latency | Selection | 2 | 51 | 201.5899 | 100.795 | 0.3637 | 0.6969 |
| Delay | Day.Latency | Day | 5 | 51 | 13171.1707 | 2634.2341 | 9.5046 | 0 |
| Delay | Day.Latency | Selection:Day | 10 | 51 | 1577.473 | 157.7473 | 0.5692 | 0.8312 |
| Delay | Night.Duration | Selection | 2 | 9 | 217.3444 | 108.6722 | 0.9657 | 0.4169 |
| Delay | Night.Duration | Day | 5 | 45 | 8275.6808 | 1655.1362 | 14.7087 | 0 |
| Delay | Night.Duration | Selection:Day | 10 | 45 | 1085.2943 | 108.5294 | 0.9645 | 0.4866 |
| Delay | Night.BoutNumber | Selection | 2 | 9 | 2.7958 | 1.3979 | 0.8489 | 0.4595 |
| Delay | Night.BoutNumber | Day | 5 | 45 | 9.7428 | 1.9486 | 1.1833 | 0.3324 |
| Delay | Night.BoutNumber | Selection:Day | 10 | 45 | 16.0801 | 1.608 | 0.9765 | 0.4768 |
| Delay | Night.BoutDuration.Median | Selection | 2 | 51 | 126.7709 | 63.3854 | 0.1618 | 0.8511 |
| Delay | Night.BoutDuration.Median | Day | 5 | 51 | 10702.505 | 2140.501 | 5.4627 | 0.0004 |
| Delay | Night.BoutDuration.Median | Selection:Day | 10 | 51 | 853.4767 | 85.3477 | 0.2178 | 0.9935 |
| Delay | Night.Latency | Selection | 2 | 8.888 | 6.9136 | 3.4568 | 3.2762 | 0.0859 |
| Delay | Night.Latency | Day | 5 | 45.001 | 313.0374 | 62.6075 | 59.3364 | 0 |
| Delay | Night.Latency | Selection:Day | 10 | 45.001 | 17.4995 | 1.7499 | 1.6585 | 0.121 |

**B**

| Parameter | Comparison | Within | Contrast (A vs B) | Estimate (A – B) | SE | df | t ratio | p (Tukey) | Significance |
|---|---|---|---|---|---|---|---|---|---|
| Daytime sleep duration | Days (lines pooled) | – | Day −3 vs Day −2 | 4.63 | 8.34 | 51.0 | 0.56 | 0.9934 | ns |
| Daytime sleep duration | Days (lines pooled) | – | Day −3 vs Day −1 | 12.37 | 8.34 | 51.0 | 1.48 | 0.6762 | ns |
| Daytime sleep duration | Days (lines pooled) | – | Day −3 vs Day 1 | 74.55 | 8.34 | 51.0 | 8.94 | **<0.0001** | *** |
| Daytime sleep duration | Days (lines pooled) | – | Day −3 vs Day 2 | 15.74 | 8.34 | 51.0 | 1.89 | 0.4219 | ns |
| Daytime sleep duration | Days (lines pooled) | – | Day −3 vs Day 3 | -14.75 | 8.34 | 51.0 | -1.77 | 0.4947 | ns |
| Daytime sleep duration | Days (lines pooled) | – | Day −2 vs Day −1 | 7.74 | 8.34 | 51.0 | 0.93 | 0.9374 | ns |
| Daytime sleep duration | Days (lines pooled) | – | Day −2 vs Day 1 | 69.92 | 8.34 | 51.0 | 8.38 | **<0.0001** | *** |
| Daytime sleep duration | Days (lines pooled) | – | Day −2 vs Day 2 | 11.11 | 8.34 | 51.0 | 1.33 | 0.7663 | ns |
| Daytime sleep duration | Days (lines pooled) | – | Day −2 vs Day 3 | -19.38 | 8.34 | 51.0 | -2.32 | 0.2038 | ns |
| Daytime sleep duration | Days (lines pooled) | – | Day −1 vs Day 1 | 62.17 | 8.34 | 51.0 | 7.45 | **<0.0001** | *** |
| Daytime sleep duration | Days (lines pooled) | – | Day −1 vs Day 2 | 3.36 | 8.34 | 51.0 | 0.40 | 0.9985 | ns |
| Daytime sleep duration | Days (lines pooled) | – | Day −1 vs Day 3 | -27.12 | 8.34 | 51.0 | -3.25 | **0.0236** | * |
| Daytime sleep duration | Days (lines pooled) | – | Day 1 vs Day 2 | -58.81 | 8.34 | 51.0 | -7.05 | **<0.0001** | *** |
| Daytime sleep duration | Days (lines pooled) | – | Day 1 vs Day 3 | -89.30 | 8.34 | 51.0 | -10.71 | **<0.0001** | *** |
| Daytime sleep duration | Days (lines pooled) | – | Day 2 vs Day 3 | -30.49 | 8.34 | 51.0 | -3.65 | **0.0076** | ** |
| Daytime sleep latency | Days (lines pooled) | – | Day −3 vs Day −2 | -4.97 | 6.80 | 51.0 | -0.73 | 0.9771 | ns |
| Daytime sleep latency | Days (lines pooled) | – | Day −3 vs Day −1 | -11.95 | 6.80 | 51.0 | -1.76 | 0.5009 | ns |
| Daytime sleep latency | Days (lines pooled) | – | Day −3 vs Day 1 | -40.96 | 6.80 | 51.0 | -6.03 | **<0.0001** | *** |
| Daytime sleep latency | Days (lines pooled) | – | Day −3 vs Day 2 | -6.97 | 6.80 | 51.0 | -1.02 | 0.9074 | ns |
| Daytime sleep latency | Days (lines pooled) | – | Day −3 vs Day 3 | -5.62 | 6.80 | 51.0 | -0.83 | 0.9612 | ns |
| Daytime sleep latency | Days (lines pooled) | – | Day −2 vs Day −1 | -6.98 | 6.80 | 51.0 | -1.03 | 0.9067 | ns |
| Daytime sleep latency | Days (lines pooled) | – | Day −2 vs Day 1 | -35.99 | 6.80 | 51.0 | -5.29 | **<0.0001** | *** |
| Daytime sleep latency | Days (lines pooled) | – | Day −2 vs Day 2 | -1.99 | 6.80 | 51.0 | -0.29 | 0.9997 | ns |
| Daytime sleep latency | Days (lines pooled) | – | Day −2 vs Day 3 | -0.65 | 6.80 | 51.0 | -0.10 | 1.0000 | ns |

| Parameter | Comparison | Within | Contrast (A vs B) | Estimate (A – B) | SE | df | t ratio | p (Tukey) | Significance |
|---|---|---|---|---|---|---|---|---|---|
| Daytime sleep latency | Days (lines pooled) | – | Day −1 vs Day 1 | -29.01 | 6.80 | 51.0 | -4.27 | **0.0012** | ** |
| Daytime sleep latency | Days (lines pooled) | – | Day −1 vs Day 2 | 4.99 | 6.80 | 51.0 | 0.73 | 0.9768 | ns |
| Daytime sleep latency | Days (lines pooled) | – | Day −1 vs Day 3 | 6.33 | 6.80 | 51.0 | 0.93 | 0.9364 | ns |
| Daytime sleep latency | Days (lines pooled) | – | Day 1 vs Day 2 | 33.99 | 6.80 | 51.0 | 5.00 | **0.0001** | *** |
| Daytime sleep latency | Days (lines pooled) | – | Day 1 vs Day 3 | 35.34 | 6.80 | 51.0 | 5.20 | **0.0001** | *** |
| Daytime sleep latency | Days (lines pooled) | – | Day 2 vs Day 3 | 1.35 | 6.80 | 51.0 | 0.20 | 1.0000 | ns |
| Daytime bout number | Days (lines pooled) | – | Day −3 vs Day −2 | -0.44 | 0.53 | 45.0 | -0.83 | 0.9600 | ns |
| Daytime bout number | Days (lines pooled) | – | Day −3 vs Day −1 | -0.75 | 0.53 | 45.0 | -1.41 | 0.7215 | ns |
| Daytime bout number | Days (lines pooled) | – | Day −3 vs Day 1 | 0.48 | 0.53 | 45.0 | 0.89 | 0.9464 | ns |
| Daytime bout number | Days (lines pooled) | – | Day −3 vs Day 2 | -1.88 | 0.53 | 45.0 | -3.52 | **0.0121** | * |
| Daytime bout number | Days (lines pooled) | – | Day −3 vs Day 3 | -3.09 | 0.53 | 45.0 | -5.79 | **<0.0001** | *** |
| Daytime bout number | Days (lines pooled) | – | Day −2 vs Day −1 | -0.31 | 0.53 | 45.0 | -0.58 | 0.9921 | ns |
| Daytime bout number | Days (lines pooled) | – | Day −2 vs Day 1 | 0.92 | 0.53 | 45.0 | 1.72 | 0.5237 | ns |
| Daytime bout number | Days (lines pooled) | – | Day −2 vs Day 2 | -1.43 | 0.53 | 45.0 | -2.69 | 0.0976 | ns |
| Daytime bout number | Days (lines pooled) | – | Day −2 vs Day 3 | -2.64 | 0.53 | 45.0 | -4.96 | **0.0001** | *** |
| Daytime bout number | Days (lines pooled) | – | Day −1 vs Day 1 | 1.23 | 0.53 | 45.0 | 2.30 | 0.2150 | ns |
| Daytime bout number | Days (lines pooled) | – | Day −1 vs Day 2 | -1.13 | 0.53 | 45.0 | -2.11 | 0.3002 | ns |
| Daytime bout number | Days (lines pooled) | – | Day −1 vs Day 3 | -2.34 | 0.53 | 45.0 | -4.38 | **0.0009** | *** |
| Daytime bout number | Days (lines pooled) | – | Day 1 vs Day 2 | -2.35 | 0.53 | 45.0 | -4.41 | **0.0008** | *** |
| Daytime bout number | Days (lines pooled) | – | Day 1 vs Day 3 | -3.56 | 0.53 | 45.0 | -6.68 | **<0.0001** | *** |
| Daytime bout number | Days (lines pooled) | – | Day 2 vs Day 3 | -1.21 | 0.53 | 45.0 | -2.27 | 0.2277 | ns |
| Daytime median bout duration | Days (lines pooled) | – | Day −3 vs Day −2 | 0.89 | 0.44 | 45.0 | 1.99 | 0.3614 | ns |
| Daytime median bout duration | Days (lines pooled) | – | Day −3 vs Day −1 | 0.54 | 0.44 | 45.0 | 1.21 | 0.8277 | ns |
| Daytime median bout duration | Days (lines pooled) | – | Day −3 vs Day 1 | 3.79 | 0.44 | 45.0 | 8.52 | **<0.0001** | *** |
| Daytime median bout duration | Days (lines pooled) | – | Day −3 vs Day 2 | 3.09 | 0.44 | 45.0 | 6.95 | **<0.0001** | *** |
| Daytime median bout duration | Days (lines pooled) | – | Day −3 vs Day 3 | 2.62 | 0.44 | 45.0 | 5.88 | **<0.0001** | *** |
| Daytime median bout duration | Days (lines pooled) | – | Day −2 vs Day −1 | -0.35 | 0.44 | 45.0 | -0.78 | 0.9695 | ns |
| Daytime median bout duration | Days (lines pooled) | – | Day −2 vs Day 1 | 2.90 | 0.44 | 45.0 | 6.52 | **<0.0001** | *** |
| Daytime median bout duration | Days (lines pooled) | – | Day −2 vs Day 2 | 2.20 | 0.44 | 45.0 | 4.95 | **0.0001** | *** |
| Daytime median bout duration | Days (lines pooled) | – | Day −2 vs Day 3 | 1.73 | 0.44 | 45.0 | 3.89 | **0.0042** | ** |
| Daytime median bout duration | Days (lines pooled) | – | Day −1 vs Day 1 | 3.25 | 0.44 | 45.0 | 7.31 | **<0.0001** | *** |
| Daytime median bout duration | Days (lines pooled) | – | Day −1 vs Day 2 | 2.55 | 0.44 | 45.0 | 5.73 | **<0.0001** | *** |
| Daytime median bout duration | Days (lines pooled) | – | Day −1 vs Day 3 | 2.08 | 0.44 | 45.0 | 4.67 | **0.0004** | *** |
| Daytime median bout duration | Days (lines pooled) | – | Day 1 vs Day 2 | -0.70 | 0.44 | 45.0 | -1.57 | 0.6210 | ns |
| Daytime median bout duration | Days (lines pooled) | – | Day 1 vs Day 3 | -1.17 | 0.44 | 45.0 | -2.64 | 0.1094 | ns |
| Daytime median bout duration | Days (lines pooled) | – | Day 2 vs Day 3 | -0.47 | 0.44 | 45.0 | -1.06 | 0.8926 | ns |
| Nighttime sleep duration | Days (lines pooled) | – | Day −3 vs Day −2 | 0.36 | 4.33 | 45.0 | 0.08 | 1.0000 | ns |
| Nighttime sleep duration | Days (lines pooled) | – | Day −3 vs Day −1 | 2.24 | 4.33 | 45.0 | 0.52 | 0.9952 | ns |
| Nighttime sleep duration | Days (lines pooled) | – | Day −3 vs Day 1 | -14.61 | 4.33 | 45.0 | -3.37 | **0.0180** | * |
| Nighttime sleep duration | Days (lines pooled) | – | Day −3 vs Day 2 | -20.74 | 4.33 | 45.0 | -4.79 | **0.0003** | *** |
| Nighttime sleep duration | Days (lines pooled) | – | Day −3 vs Day 3 | -24.06 | 4.33 | 45.0 | -5.55 | **<0.0001** | *** |
| Nighttime sleep duration | Days (lines pooled) | – | Day −2 vs Day −1 | 1.88 | 4.33 | 45.0 | 0.43 | 0.9979 | ns |
| Nighttime sleep duration | Days (lines pooled) | – | Day −2 vs Day 1 | -14.97 | 4.33 | 45.0 | -3.46 | **0.0144** | * |
| Nighttime sleep duration | Days (lines pooled) | – | Day −2 vs Day 2 | -21.10 | 4.33 | 45.0 | -4.87 | **0.0002** | *** |
| Nighttime sleep duration | Days (lines pooled) | – | Day −2 vs Day 3 | -24.41 | 4.33 | 45.0 | -5.64 | **<0.0001** | *** |
| Nighttime sleep duration | Days (lines pooled) | – | Day −1 vs Day 1 | -16.85 | 4.33 | 45.0 | -3.89 | **0.0042** | ** |
| Nighttime sleep duration | Days (lines pooled) | – | Day −1 vs Day 2 | -22.98 | 4.33 | 45.0 | -5.31 | **<0.0001** | *** |
| Nighttime sleep duration | Days (lines pooled) | – | Day −1 vs Day 3 | -26.30 | 4.33 | 45.0 | -6.07 | **<0.0001** | *** |
| Nighttime sleep duration | Days (lines pooled) | – | Day 1 vs Day 2 | -6.14 | 4.33 | 45.0 | -1.42 | 0.7168 | ns |
| Nighttime sleep duration | Days (lines pooled) | – | Day 1 vs Day 3 | -9.45 | 4.33 | 45.0 | -2.18 | 0.2664 | ns |
| Nighttime sleep duration | Days (lines pooled) | – | Day 2 vs Day 3 | -3.31 | 4.33 | 45.0 | -0.76 | 0.9720 | ns |
| Nighttime sleep latency | Days (lines pooled) | – | Day −3 vs Day −2 | -0.27 | 0.42 | 45.0 | -0.64 | 0.9869 | ns |
| Nighttime sleep latency | Days (lines pooled) | – | Day −3 vs Day −1 | 0.21 | 0.42 | 45.0 | 0.51 | 0.9956 | ns |
| Nighttime sleep latency | Days (lines pooled) | – | Day −3 vs Day 1 | 5.85 | 0.42 | 45.0 | 13.95 | **<0.0001** | *** |
| Nighttime sleep latency | Days (lines pooled) | – | Day −3 vs Day 2 | 1.81 | 0.42 | 45.0 | 4.32 | **0.0011** | ** |

| Parameter | Comparison | Within | Contrast (A vs B) | Estimate (A – B) | SE | df | t ratio | p (Tukey) | Significance |
|---|---|---|---|---|---|---|---|---|---|
| Nighttime sleep latency | Days (lines pooled) | – | Day −3 vs Day 3 | 1.72 | 0.42 | 45.0 | 4.10 | **0.0022** | ** |
| Nighttime sleep latency | Days (lines pooled) | – | Day −2 vs Day −1 | 0.48 | 0.42 | 45.0 | 1.15 | 0.8572 | ns |
| Nighttime sleep latency | Days (lines pooled) | – | Day −2 vs Day 1 | 6.12 | 0.42 | 45.0 | 14.59 | **<0.0001** | *** |
| Nighttime sleep latency | Days (lines pooled) | – | Day −2 vs Day 2 | 2.08 | 0.42 | 45.0 | 4.97 | **0.0001** | *** |
| Nighttime sleep latency | Days (lines pooled) | – | Day −2 vs Day 3 | 1.99 | 0.42 | 45.0 | 4.75 | **0.0003** | *** |
| Nighttime sleep latency | Days (lines pooled) | – | Day −1 vs Day 1 | 5.64 | 0.42 | 45.0 | 13.44 | **<0.0001** | *** |
| Nighttime sleep latency | Days (lines pooled) | – | Day −1 vs Day 2 | 1.60 | 0.42 | 45.0 | 3.82 | **0.0052** | ** |
| Nighttime sleep latency | Days (lines pooled) | – | Day −1 vs Day 3 | 1.51 | 0.42 | 45.0 | 3.60 | **0.0098** | ** |
| Nighttime sleep latency | Days (lines pooled) | – | Day 1 vs Day 2 | -4.04 | 0.42 | 45.0 | -9.63 | **<0.0001** | *** |
| Nighttime sleep latency | Days (lines pooled) | – | Day 1 vs Day 3 | -4.13 | 0.42 | 45.0 | -9.85 | **<0.0001** | *** |
| Nighttime sleep latency | Days (lines pooled) | – | Day 2 vs Day 3 | -0.09 | 0.42 | 45.0 | -0.22 | 0.9999 | ns |
| Nighttime median bout duration | Days (lines pooled) | – | Day −3 vs Day −2 | -1.30 | 8.08 | 51.0 | -0.16 | 1.0000 | ns |
| Nighttime median bout duration | Days (lines pooled) | – | Day −3 vs Day −1 | -0.20 | 8.08 | 51.0 | -0.02 | 1.0000 | ns |
| Nighttime median bout duration | Days (lines pooled) | – | Day −3 vs Day 1 | 2.28 | 8.08 | 51.0 | 0.28 | 0.9997 | ns |
| Nighttime median bout duration | Days (lines pooled) | – | Day −3 vs Day 2 | -6.76 | 8.08 | 51.0 | -0.84 | 0.9592 | ns |
| Nighttime median bout duration | Days (lines pooled) | – | Day −3 vs Day 3 | -33.07 | 8.08 | 51.0 | -4.09 | **0.0020** | ** |
| Nighttime median bout duration | Days (lines pooled) | – | Day −2 vs Day −1 | 1.11 | 8.08 | 51.0 | 0.14 | 1.0000 | ns |
| Nighttime median bout duration | Days (lines pooled) | – | Day −2 vs Day 1 | 3.58 | 8.08 | 51.0 | 0.44 | 0.9977 | ns |
| Nighttime median bout duration | Days (lines pooled) | – | Day −2 vs Day 2 | -5.46 | 8.08 | 51.0 | -0.68 | 0.9839 | ns |
| Nighttime median bout duration | Days (lines pooled) | – | Day −2 vs Day 3 | -31.76 | 8.08 | 51.0 | -3.93 | **0.0033** | ** |
| Nighttime median bout duration | Days (lines pooled) | – | Day −1 vs Day 1 | 2.48 | 8.08 | 51.0 | 0.31 | 0.9996 | ns |
| Nighttime median bout duration | Days (lines pooled) | – | Day −1 vs Day 2 | -6.57 | 8.08 | 51.0 | -0.81 | 0.9639 | ns |
| Nighttime median bout duration | Days (lines pooled) | – | Day −1 vs Day 3 | -32.87 | 8.08 | 51.0 | -4.07 | **0.0022** | ** |
| Nighttime median bout duration | Days (lines pooled) | – | Day 1 vs Day 2 | -9.04 | 8.08 | 51.0 | -1.12 | 0.8714 | ns |
| Nighttime median bout duration | Days (lines pooled) | – | Day 1 vs Day 3 | -35.35 | 8.08 | 51.0 | -4.37 | **0.0008** | *** |
| Nighttime median bout duration | Days (lines pooled) | – | Day 2 vs Day 3 | -26.30 | 8.08 | 51.0 | -3.25 | **0.0234** | * |

**Table S12. Sleep rebound after sleep deprivation:** Two-way randomized block ANOVA for day (0–12 h), night (12–24 h) and 24 h sleep rebound after 12 h of night sleep deprivation in male chronotype flies, using standard (std) (A), long (B), intermediate (int) (C) and short (D) sleep definitions, with Tukey's HSD post hoc comparisons for night rebound of long and intermediate sleep.

A

| **std-day rebound** | **Effect** | **SS** | **$DF_n$** | **MS** | **$DF_d$** | **MS error** | **F** | **p** |
|---|---|---|---|---|---|---|---|---|
| Intercept | Fixed | 155014.8 | 1 | 155014.8 | 3 | 750.3942 | 206.578 | 0.001 |
| SELECTION | Fixed | 2534.8 | 2 | 1267.4 | 6 | 535.1237 | 2.368 | 0.175 |
| BLOCK | Random | 2251.2 | 3 | 750.4 | 6 | 535.1237 | 1.402 | 0.331 |
| SELECTION*BLOCK | Random | 3210.7 | 6 | 535.1 | 0 | 0 | | |
| Error | | | 0 | | | | | |
| **std- night rebound** | **Effect** | **SS** | **$DF_n$** | **MS** | **$DF_d$** | **MS error** | **F** | **p** |
| Intercept | Fixed | 10570.18 | 1 | 10570.18 | 3 | 519.7046 | 20.339 | 0.020 |
| SELECTION | Fixed | 1543.04 | 2 | 771.52 | 6 | 311.2532 | 2.479 | 0.164 |
| BLOCK | Random | 1559.11 | 3 | 519.7 | 6 | 311.2532 | 1.670 | 0.271 |
| SELECTION*BLOCK | Random | 1867.52 | 6 | 311.25 | 0 | 0 | | |
| Error | | | 0 | | | | | |
| **std - 24h rebound** | **Effect** | **SS** | **$DF_n$** | **MS** | **$DF_d$** | **MS error** | **F** | **p** |
| Intercept | Fixed | 251143.4 | 1 | 251143.4 | 3 | 1962.074 | 127.999 | 0.001 |
| SELECTION | Fixed | 4394.2 | 2 | 2197.1 | 6 | 1296.56 | 1.695 | 0.261 |
| BLOCK | Random | 5886.2 | 3 | 1962.1 | 6 | 1296.56 | 1.513 | 0.304 |
| SELECTION*BLOCK | Random | 7779.4 | 6 | 1296.6 | 0 | 0 | | |
| Error | | | 0 | | | | | |

B

| **long- day rebound** | **Effect** | **SS** | **$DF_n$** | **MS** | **$DF_d$** | **MS error** | **F** | **p** |
|---|---|---|---|---|---|---|---|---|
| Intercept | Fixed | 165599.8 | 1 | 165599.8 | 3 | 5130.955 | 32.275 | 0.011 |
| SELECTION | Fixed | 38.7 | 2 | 19.4 | 6 | 537.357 | 0.036 | 0.965 |
| BLOCK | Random | 15392.9 | 3 | 5131 | 6 | 537.357 | 9.549 | 0.011 |

| | | | | | | | | |
|---|---|---|---|---|---|---|---|---|
| SELECTION*BLOCK | Random | 3224.1 | 6 | 537.4 | 0 | 0 | | |
| Error | | | 0 | | | | | |
| **long- night rebound** | **Effect** | **SS** | **$DF_n$** | **MS** | **$DF_d$** | **MS error** | **F** | **p** |
| Intercept | Fixed | 12560.27 | 1 | 12560.27 | 3 | 3232.895 | 3.885 | 0.143 |
| SELECTION | Fixed | 14631.67 | 2 | 7315.84 | 6 | 702.255 | 10.418 | 0.011 |
| BLOCK | Random | 9698.68 | 3 | 3232.89 | 6 | 702.255 | 4.604 | 0.053 |
| SELECTION*BLOCK | Random | 4213.53 | 6 | 702.26 | 0 | 0 | | |
| Error | | | 0 | | | | | |
| **long- 24h rebound** | **Effect** | **SS** | **$DF_n$** | **MS** | **$DF_d$** | **MS error** | **F** | **p** |
| Intercept | Fixed | 269373.6 | 1 | 269373.6 | 3 | 16155.35 | 16.674 | 0.027 |
| SELECTION | Fixed | 13268.9 | 2 | 6634.5 | 6 | 2041.86 | 3.249 | 0.111 |
| BLOCK | Random | 48466.1 | 3 | 16155.4 | 6 | 2041.86 | 7.912 | 0.017 |
| SELECTION*BLOCK | Random | 12251.1 | 6 | 2041.9 | 0 | 0 | | |
| Error | | | 0 | | | | | |

C

| **int- day rebound** | **Effect** | **SS** | **$DF_n$** | **MS** | **$DF_d$** | **MS error** | **F** | **p** |
|---|---|---|---|---|---|---|---|---|
| Intercept | Fixed | 455.478 | 1 | 455.478 | 3 | 1253.556 | 0.363 | 0.589 |
| selection | Fixed | 315.958 | 2 | 157.979 | 6 | 438.827 | 0.360 | 0.712 |
| block | Random | 3760.668 | 3 | 1253.556 | 6 | 438.827 | 2.857 | 0.127 |
| selection*block | Random | 2632.96 | 6 | 438.827 | 0 | 0 | | |
| Error | | | 0 | | | | | |
| **int- night rebound** | **Effect** | **SS** | **$DF_n$** | **MS** | **$DF_d$** | **MS error** | **F** | **p** |
| Intercept | Fixed | 1140.016 | 1 | 1140.016 | 3 | 3222.738 | 0.354 | 0.594 |
| selection | Fixed | 2817.475 | 2 | 1408.737 | 6 | 155.352 | 9.068 | 0.015 |
| block | Random | 9668.213 | 3 | 3222.738 | 6 | 155.352 | 20.745 | 0.001 |
| selection*block | Random | 932.109 | 6 | 155.352 | 0 | 0 | | |
| Error | | | 0 | | | | | |
| **int-24h rebound** | **Effect** | **SS** | **$DF_n$** | **MS** | **$DF_d$** | **MS error** | **F** | **p** |
| Intercept | Fixed | 154.31 | 1 | 154.311 | 3 | 6790.964 | 0.023 | 0.890 |
| selection | Fixed | 2172.37 | 2 | 1086.184 | 6 | 217.89 | 4.985 | **0.053** |
| block | Random | 20372.89 | 3 | 6790.964 | 6 | 217.89 | 31.167 | 0.000 |
| selection*block | Random | 1307.34 | 6 | 217.89 | 0 | 0 | | |
| Error | | | 0 | | | | | |

D

| **short- day rebound** | **Effect** | **SS** | **$DF_n$** | **MS** | **$DF_d$** | **MS error** | **F** | **p** |
|---|---|---|---|---|---|---|---|---|
| Intercept | Fixed | 626.871 | 1 | 626.871 | 3 | 2084.509 | 0.301 | 0.622 |
| selection | Fixed | 949.924 | 2 | 474.962 | 6 | 345.322 | 1.375 | 0.322 |
| block | Random | 6253.528 | 3 | 2084.509 | 6 | 345.322 | 6.036 | 0.030 |
| selection*block | Random | 2071.934 | 6 | 345.322 | 0 | 0 | | |
| Error | | | 0 | | | | | |
| **short- night rebound** | **Effect** | **SS** | **$DF_n$** | **MS** | **$DF_d$** | **MS error** | **F** | **p** |
| Intercept | Fixed | 79.932 | 1 | 79.932 | 3 | 277.9336 | 0.288 | 0.629 |
| selection | Fixed | 2243.016 | 2 | 1121.508 | 6 | 368.8359 | 3.041 | 0.122 |
| block | Random | 833.801 | 3 | 277.934 | 6 | 368.8359 | 0.754 | 0.559 |
| selection*block | Random | 2213.015 | 6 | 368.836 | 0 | 0 | | |
| Error | | | 0 | | | | | |
| **short-24h rebound** | **Effect** | **SS** | **$DF_n$** | **MS** | **$DF_d$** | **MS error** | **F** | **p** |
| Intercept | Fixed | 259.111 | 1 | 259.111 | 3 | 2478.293 | 0.105 | 0.768 |
| selection | Fixed | 4729.404 | 2 | 2364.702 | 6 | 826.289 | 2.862 | 0.134 |
| block | Random | 7434.878 | 3 | 2478.293 | 6 | 826.289 | 2.999 | 0.117 |
| selection*block | Random | 4957.733 | 6 | 826.289 | 0 | 0 | | |
| Error | | | 0 | | | | | |

| Tukey's HSD test | | |
|---|---|---|
| **Mean difference** | **long- night rebound** | **int-night rebound** |
| ***early vs control*** | 13.11436 | -27.4168 |
| ***early vs late*** | 79.75477 | -35.9073 |
| ***control vs late*** | 66.64041 | -8.4906 |
| Minimum significant difference | 57.4920 | 27.0407 |
| q critical | 4.339 | 4.339 |
| K, N | 3, 4 | 3, 4 |

**Table S13. Sleep fragmentation before and after sleep deprivation:** Linear mixed-effects model (selection as a fixed factor, day as a repeated-measures factor and block as a random factor) for nighttime number of bouts and median bout duration of sleep-deprived male flies on Day 1 (pre-deprivation) and Day 3 (post-deprivation). (A) Type III ANOVA with Satterthwaite df. (B) Tukey's HSD post hoc comparisons for significant terms.

A

| Parameter | Term | SS | $DF_n$ | $DF_d$ | MS error | F | p |
|---|---|---|---|---|---|---|---|
| Nighttime bout number | Selection | 3.9354 | 2 | 6 | 0.4099 | 4.8008 | 0.0569 |
| Nighttime bout number | Day | 5.4492 | 1 | 9 | 0.4099 | 13.2948 | 0.0053 |
| Nighttime bout number | Selection × Day | 11.7408 | 2 | 9 | 0.4099 | 14.3223 | 0.0016 |
| Nighttime median bout duration | Selection | 2241.9931 | 2 | 15 | 85.3041 | 13.1412 | 0.0005 |
| Nighttime median bout duration | Day | 22.4365 | 1 | 15 | 85.3041 | 0.263 | 0.6155 |
| Nighttime median bout duration | Selection × Day | 112.8805 | 2 | 15 | 85.3041 | 0.6616 | 0.5304 |

B

| Parameter | Comparison | Within | Contrast | Estimate | SE | df | t | p |
|---|---|---|---|---|---|---|---|---|
| Nighttime bout number | Lines within each day | Day 1 | *early vs control* | 1.0144 | 0.7267 | 8.8932 | 1.3959 | 0.3834 |
| Nighttime bout number | Lines within each day | Day 1 | *early vs late* | 2.9883 | 0.7267 | 8.8932 | 4.1121 | 0.0068 |
| Nighttime bout number | Lines within each day | Day 1 | *control vs late* | 1.9739 | 0.7267 | 8.8932 | 2.7162 | 0.0568 |
| Nighttime bout number | Lines within each day | Day 3 | *early vs control* | 2.1621 | 0.7267 | 8.8932 | 2.9752 | 0.038 |
| Nighttime bout number | Lines within each day | Day 3 | *early vs late* | 0.7661 | 0.7267 | 8.8932 | 1.0543 | 0.564 |
| Nighttime bout number | Lines within each day | Day 3 | *control vs late* | -1.396 | 0.7267 | 8.8932 | -1.9209 | 0.1889 |
| Nighttime bout number | Days within each line | *early* | Day 1 vs Day 3 | -0.5948 | 0.4527 | 9 | -1.314 | 0.2214 |
| Nighttime bout number | Days within each line | *control* | Day 1 vs Day 3 | 0.5529 | 0.4527 | 9 | 1.2212 | 0.253 |
| Nighttime bout number | Days within each line | *late* | Day 1 vs Day 3 | -2.817 | 0.4527 | 9 | -6.2226 | 0.0002 |
| Nighttime median bout duration | Lines (days pooled) | - | *early vs control* | -14.2124 | 4.618 | 15 | -3.0776 | 0.0197 |
| Nighttime median bout duration | Lines (days pooled) | - | *early vs late* | -23.5037 | 4.618 | 15 | -5.0896 | 0.0004 |
| Nighttime median bout duration | Lines (days pooled) | - | *control vs late* | -9.2913 | 4.618 | 15 | -2.012 | 0.1435 |
